# Regulatory T cells clonally expand and contribute to stromal cell function in fibrotic response to synthetic implants

**DOI:** 10.64898/2026.01.05.697727

**Authors:** Joscelyn C. Mejías, Kavita Krishnan, Anna Ruta, A. Shri Ramanujam, Elise F. Gray-Gaillard, Locke Davenport Huyer, David R. Maestas, Sushma Nagaraj, Frank Haoning Yu, Alexandra N. Rindone, Peter Abraham, Maria Browne, Elana J. Fertig, Drew M. Pardoll, Jennifer H. Elisseeff

**Affiliations:** Translational Tissue Engineering Center, Department of Chemical and Biomolecular Engineering and Biomedical Engineering, Johns Hopkins School of Medicine, Baltimore, MD, USA; Wallace H. Coulter Department of Biomedical Engineering, Georgia Institute of Technology and Emory University; Parker H. Petit Institute for Bioengineering and Bioscience, Georgia Institute of Technology, Atlanta, GA, USA; Bloomberg-Kimmel Institute for Cancer Immunotherapy, Sidney Kimmel Comprehensive Cancer Center, Johns Hopkins School of Medicine, Baltimore, MD, USA; Department of Biomaterials & Applied Oral Sciences, School of Biomedical Engineering, Department of Microbiology & Immunology, Dalhousie University, Halifax, NS, Canada; Institute for Genome Sciences, University of Maryland School of Medicine, Baltimore, MD, USA; Greenebaum Comprehensive Cancer Center, Department of Medicine, and Department of Epidemiology, University of Maryland School of Medicine, Baltimore, MD, USA; Department of Chemical Engineering, Johns Hopkins University, Baltimore, MD, USA

## Abstract

Fibrosis plays a key role in both chronic disease progression and failure of synthetic biomaterial implants. However, the contribution of adaptive immunity to fibrotic development remains incompletely understood, particularly for regulatory T cells (Tregs). Here, we used single-cell multiomic profiling, integrating transcriptomics with T cell receptor (TCR) sequencing, to map Treg heterogeneity and clonal dynamics in a synthetic material-induced model of fibrosis. We uncovered progressive Treg clonal expansion accompanied by TCR activation signatures and an increasingly immunosuppressive phenotype along a continuous transcriptional trajectory. These Tregs suppressed immune responses and influenced extracellular matrix and vascular gene expression. Cell-cell communication inference predicted Treg-driven activation of pro-fibrotic and vasculogenic transcriptional programs in fibroblasts and endothelial cells, including Sox-family transcription factors. Functional Treg depletion increased inflammation and significantly reduced neovascularization. Together, these findings identify Treg functions in the fibro-vascular niche through stromal cell modulation, highlighting immune-stromal interactions as an important axis in fibrosis.

## Introduction

Fibrosis, the pathological deposition of extracellular matrix (ECM) that compromises tissue and organ function, is a common characteristic of numerous chronic diseases and a major feature of age-related decline.^1–3^ Although fibrotic diseases arise from diverse causes—such as idiopathic pulmonary fibrosis (IPF) and non-alcoholic fatty liver disease (NAFLD)—they share core underlying mechanisms.^4,5^ These include dysregulated immune responses, abnormal vascular remodeling, and persistent fibroblast activation.^6^ Dissecting these intertwined processes in complex disease states, including immune-stromal signaling, remains a significant challenge.^7^ The foreign body response (FBR) to implanted materials is characterized by fibrosis and provides a robust and reproducible *in vivo* model to interrogate the development of fibrosis.^7^

The FBR is an inflammatory response to implantation of foreign material. This cascade is initiated by protein adsorption to the material surface, which rapidly recruits innate immune cells, including neutrophils, monocytes, and macrophages.^7^ A core component of the fibrotic response lies in a signaling axis between immune cells and resident fibroblasts, driving their differentiation into ECM-secreting myofibroblasts.^6,8^ While macrophages are critical drivers of this process, often fusing into multinucleated giant cells at the material interface, T lymphocytes are also present in the fibrotic niche.^9–11^ Our previous work demonstrated that T helper 17 (Th17) cells, in conjunction with senescent cells, establish a pro-fibrotic feed forward signaling network contributing to biomaterial induced fibrosis and post-traumatic osteoarthritis.^12,13^ However, the role of other T cell subsets, such as regulatory T cells (Tregs), is less understood. Furthermore, the unique function of T cells introduces questions of antigen recognition, T cell receptor (TCR) signaling, and possible clonal expansion of T cells in fibrotic tissue environments. Advances in single-cell RNA sequencing (scRNAseq) with paired single-cell TCR sequencing (scTCRseq) enables detailed characterization of T cells including phenotype and clonal expansion. When combined with predictive algorithms for cell-to-cell communication, these methods help reveal immune–stromal networks, including those driven by Tregs.

Tregs, characterized by the expression of transcription factor (TF) forkhead box P3 (Foxp3), are essential for maintaining immune homeostasis.^14^ They exert their suppressive functions through diverse mechanisms, including secretion of cytokines such as Interleukin-10 (IL-10) and transforming growth factor-β (TGF-β), direct contact-dependent inhibition of effector cells, and disruption of adenosine metabolism.^15,16^ Based on their immunosuppressive properties, Treg-based therapies are being explored for autoimmune diseases and transplant tolerance. ^17^ In the context of cancer, Treg immunosuppression hinders the immune response to tumors.^18,19^ More recently, researchers found that Tregs modulate the ECM and vascular tissue structure in the tumor microenvironment, suggesting that Tregs contribute to more than immunosuppression.^20^ The recognition that Tregs play a vital role in muscle tissue repair, along with the discovery that they undergo clonal expansion, sheds light on their diverse functions and contributions to tissue structure.^21,22^ In the case of dysregulated repair fibrosis, the contribution of Tregs is less understood. Although Tregs are known to contribute to chronic inflammation and age-related fibrosis during muscle repair, the composition and kinetics of Treg populations within the fibrotic niche remain poorly defined.^23^ It also remains unclear whether Treg clonal selection occurs during fibrosis, or how such selection might influence stromal activation, matrix production, and vascularization.

In this study, we investigated the functional importance and clonal dynamics of Tregs in a biomaterial-induced murine model of fibrosis. Using paired single-cell RNA and TCR sequencing (scRNA/TCRseq), we observed clonal expansion of differentiated and immunosuppressive Tregs in the fibrotic environment. Bioinformatics analysis predicted communication between Treg populations and key stromal cells, fibroblasts and endothelial cells, in part via activation of multiple Sox-family TFs. *In vitro* co-culture studies supported Treg communication with fibroblasts with significant increases in collagen gene expression observed in fibroblasts cultured with Tregs. *In vivo* Treg depletion confirmed the immunosuppressive effects of the Tregs in the fibrotic environment in addition to significant reductions in neovascularization. Together, these findings demonstrate a Treg clonal response in the fibrotic niche and reveal a role for Tregs in influencing immune-stromal interactions that drive fibrotic remodeling and vascular development.

## Results

### Immunosuppressive regulatory T cells accumulate during fibrosis

To model the role of Tregs in fibrosis induced by the FBR, we used a volumetric muscle loss (VML) injury in C57BL/6 mice treated with polycaprolactone (PCL), a synthetic polymer that consistently induces a robust and reproducible fibrotic response (schematic Fig 1A).^12,24^ Using spectral flow cytometry, we characterized the kinetics of Treg accumulation surrounding the material implant, comparing PCL-treated and saline-treated muscle at 1, 3, and 6 weeks post-surgery (n = 5 per group; STable 1, SFig 1). Total T cell numbers peaked at 1 week after surgery with PCL-implanted tissues containing significantly more T cells than saline controls (31,207 ± 7,529 cells vs 6,817 ± 1,521 cells; Fig 1B). In the PCL implant group, T cell numbers plateaued at 3 weeks and remained stable through 6 weeks post implantation, whereas saline controls showed decreasing numbers at 3 weeks with negligible counts by 6 weeks.

**Figure 1:**
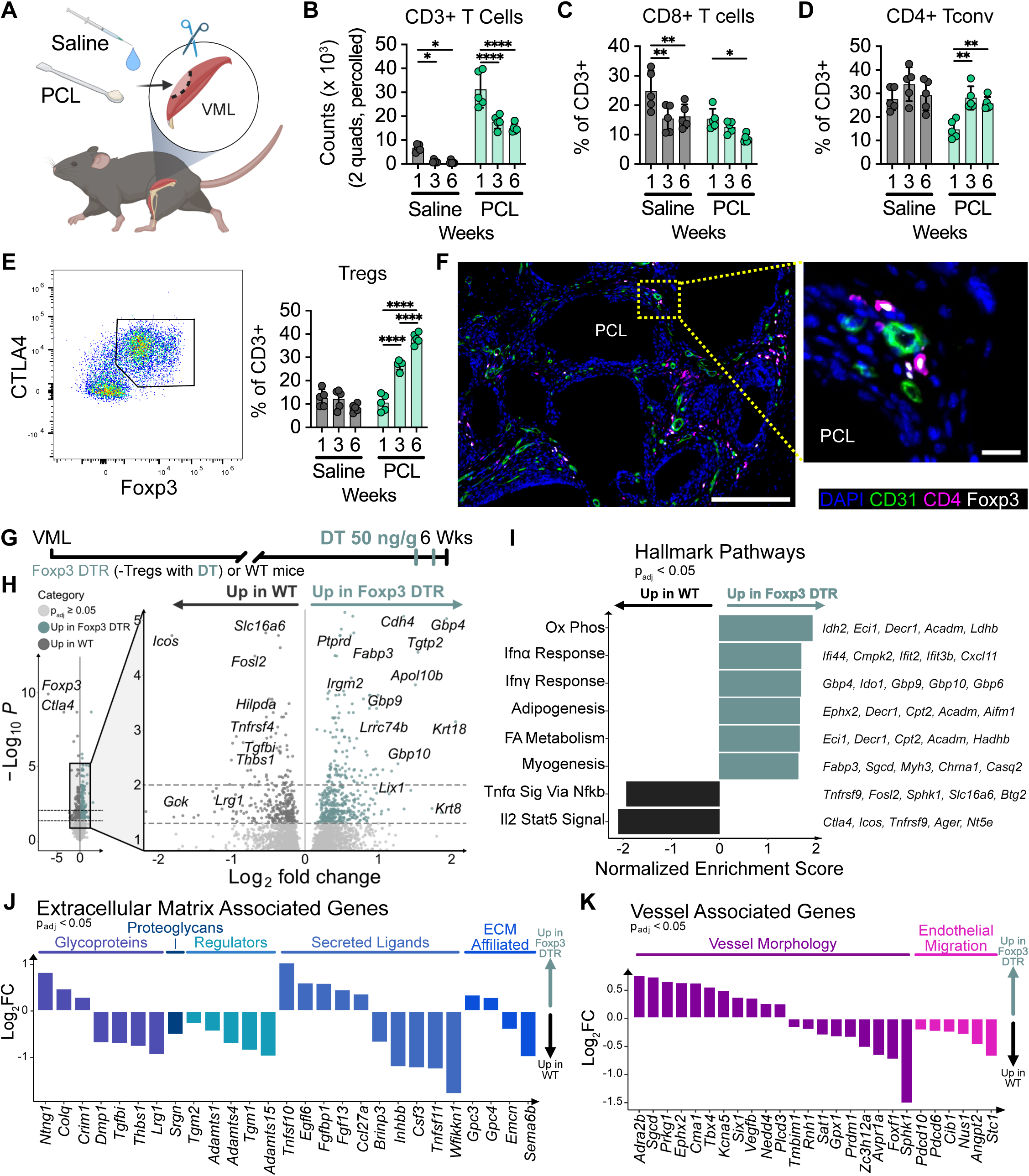
Immunosuppressive regulatory T cells accumulate in fibrotic tissue and modulate transcriptional changes in the stroma. A) Schematic depicting volumetric muscle loss (VML) injury in the quadriceps (quads) with poly-caprolactone (PCL) implant or saline control used to model the foreign body response (FBR). B) Flow cytometry counts of percolled tissue for CD3+ T cells at 1, 3, and 6 weeks after saline or PCL implant (n = 5 per group). C) Percentage of CD8+ and (D) CD4+ conventional T cells. E) Representative flow plot and percentage of regulatory T cells (Tregs) in CD3+ T cells. F) Representative immunofluorescence images of quads 6 weeks after PCL implant stained for DAPI (blue), CD31 (green), CD4 (magenta) and Foxp3 (white). Scale bars: overview = 200 µm, inset = 20 µm. G) Timeline of Treg depletion model with diphtheria toxin (DT) treatment 3 and 1 days before harvest 6 weeks after PCL implant in wildtype (WT) and Foxp3 DTR mice (n = 5 per group). H) Volcano plot displaying differential gene expression between WT and Foxp3 DTR mice from bulk RNA sequencing. I) Normalized enrichment score of significantly enriched Hallmark pathways from gene set enrichment analysis of (B). J) Log fold change of significantly differentially expressed genes associated with extracellular matrix and (K) vasculature. Bar plots indicate mean ± SD. Statistical analysis was performed using two-way ANOVA with Tukey’s post hoc test within sample group (B-E). p < 0.05 (*), p < 0.01 (**), p < 0.001 (***), p < 0.0001(****).

As fibrosis progressed around the PCL, the T cell compartment shifted. The proportion of CD8+ T cells decreased over time, while conventional CD4+ T cells increased modestly (Fig 1C-D). Notably, Tregs, defined by expression of Foxp3+ and cytotoxic T-lymphocyte associated protein 4 (CTLA4), increased in proportion from 11 ± 3.6 % at week 1 to become the predominant subset by week 6, representing 38 ± 2.4% of the total T cell population (Fig 1E). Tregs in the fibrotic tissue uniformly expressed activation markers CTLA4 and glucocorticoid-induced TNFR-related protein (GITR), in contrast with inguinal lymph node Tregs (STable 1, SFig 2). Immunofluorescence imaging confirmed the presence of CD4+ Foxp3+ Tregs within the fibrotic capsule and associated vasculature (Fig 1F, SFig 3).

**Figure 2:**
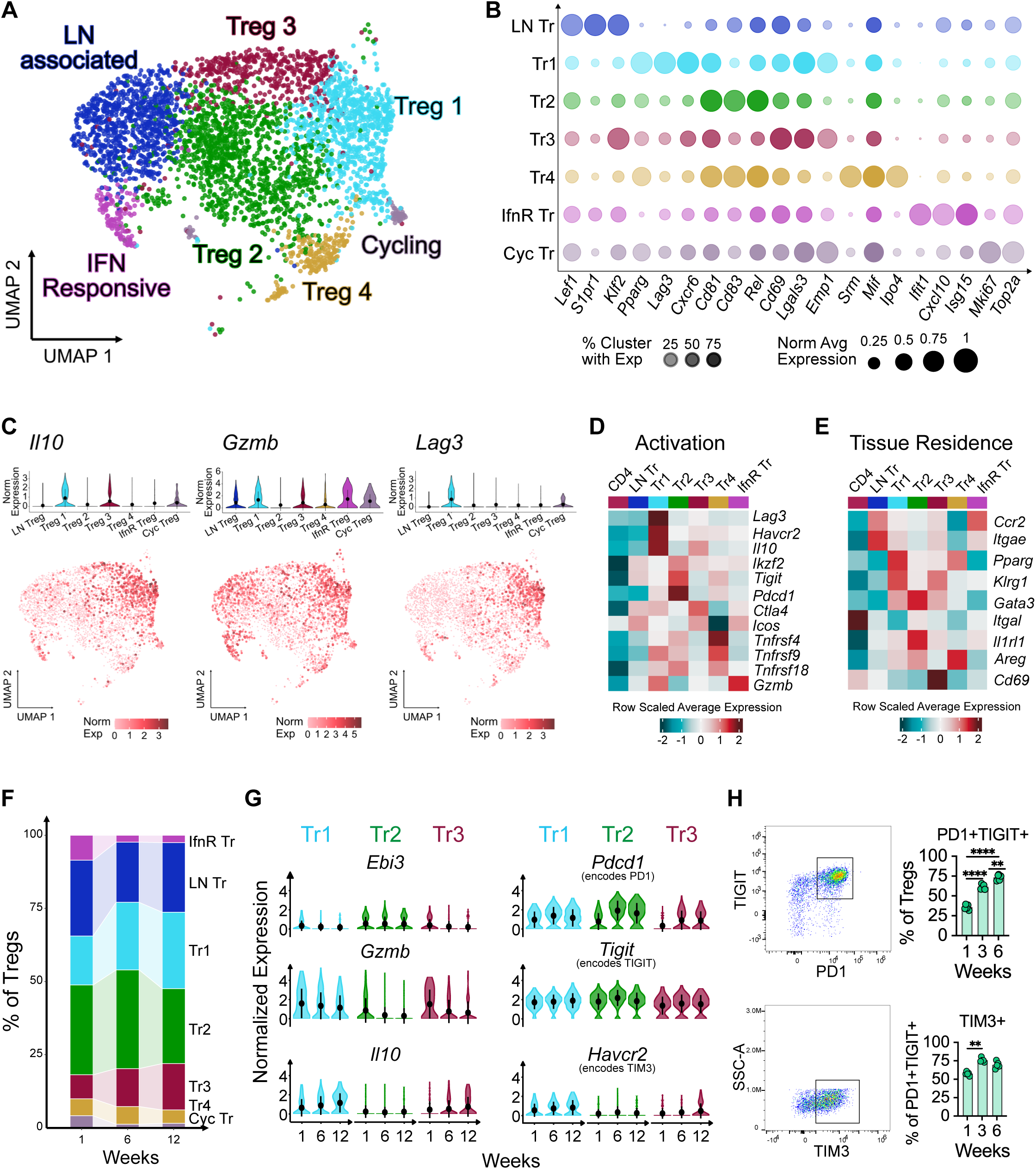
Activated, tissue-resident Tregs increase suppressive phenotype as fibrosis progresses. A) UMAP of 3,678 Tregs isolated from CD3 scRNAseq, colored by cluster. B) Dot plot displaying normalized average expression (dot size) and percentage of expressing cells (dot transparency) for genes defining each Treg cluster. C) Violin plots by cluster (top) and feature plots on UMAP (bottom) showing expression of effector Treg genes. D) Row scaled average expression of Treg activation genes and (E) tissue residence markers across cluster, including CD4 cluster from CD3 scRNAseq as reference. F) Percentage of Treg subsets at 1, 6, and 12 weeks after PCL implant. G) Normalized expression of genes across 1, 6, and 12 weeks after PCL implant within Tregs 1, 2, and 3. H) Representative flow cytometry gating (left) and quantification (right) of PD1+ TIGIT+ Tregs and TIM3+ Tregs 1, 3, and 6 weeks after PCL implant. Violin plots show data distribution with overlaid point and line indicating mean ± SD. Bar plots indicate mean ± SD. Statistical analysis was performed using two-way ANOVA with Tukey’s post hoc test within sample group. p < 0.05 (*), p < 0.01 (**), p < 0.001 (***), p < 0.0001(****).

**Figure 3:**
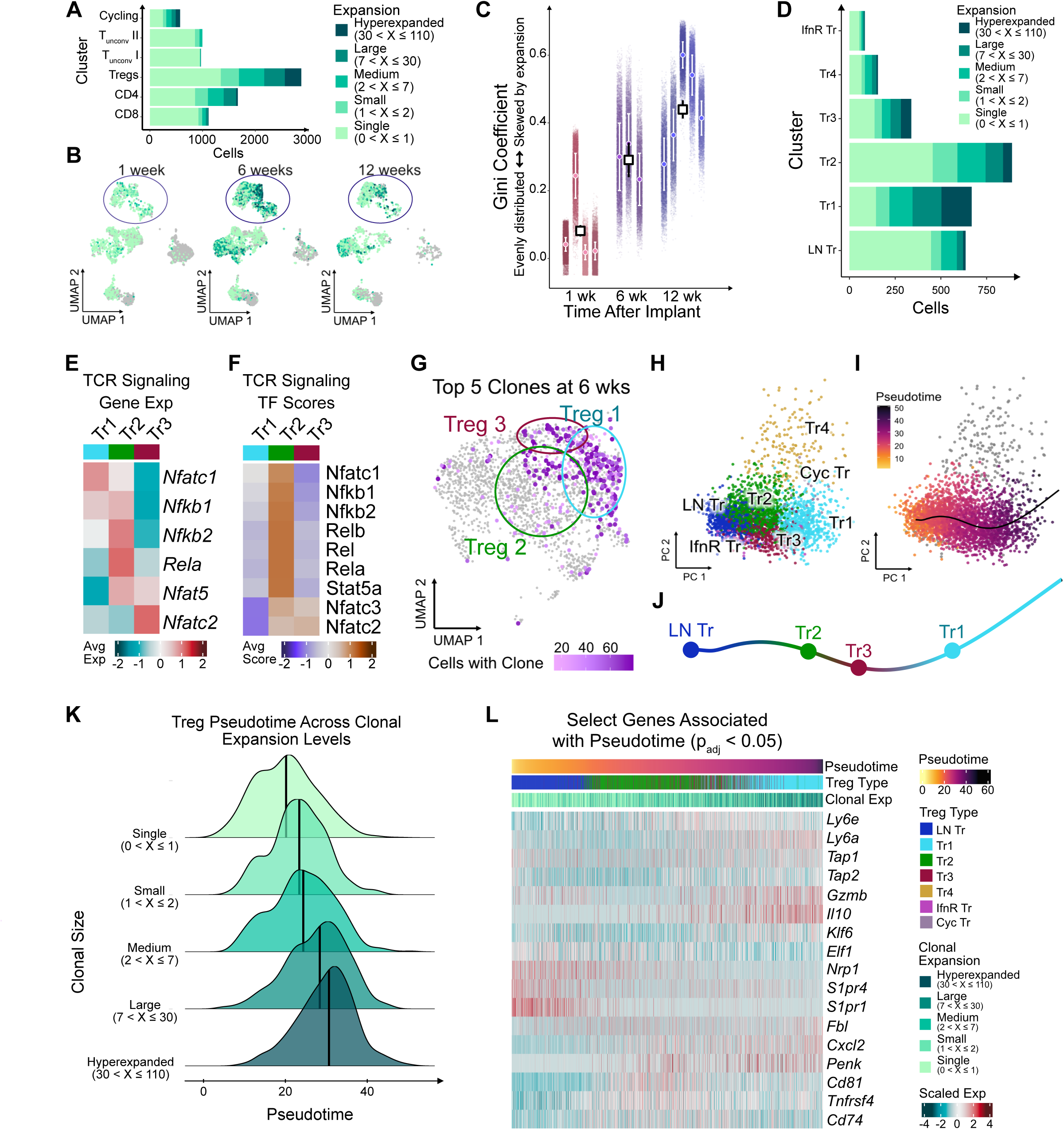
Immunosuppressive Tregs undergo clonal expansion with TCR activation during fibrotic progression. A) Cell counts for T cell clones categorized by expansion state across populations in CD3 scRNAseq. B) UMAP of CD3 scRNAseq split by time point and colored by clonal expansion category. Circles highlight emergence of expanded clones in Tregs over time. C) Gini coefficient of the CD3 clonal repertoire calculated per replicate where individual points represent 10000 random downsamples; circles indicate replicate mean ± SD, squares represent timepoint mean ± SEM. D) Cell counts for T cell clones categorized by expansion state across Treg clusters. E) Scaled average gene expression and (F) transcription factor (TF) activation associated with TCR signaling in Tregs 1, 2, and 3. G) UMAP of Tregs highlighting top 5 clones (ties allowed) 6 weeks after PCL implant, with color indicating clone size. H) Treg subpopulations displayed on batch corrected principle components, as used for trajectory inference. I) Principal components colored by pseudotime along main pseudotime trajectory. J) Position of Treg clusters along main trajectory. K) Ridge plot showing distribution of Tregs across pseudotime, split by clonal expansion category; lines indicate median of distribution. L) Row scaled expression of select genes significantly associated with pseudotime of main trajectory in each cell.

To evaluate the functional significance of this expanding Treg population in fibrosis, we performed a short Treg depletion using a Foxp3 diphtheria toxin receptor (DTR) model 3 days prior to harvest at 6 weeks post-surgery (Fig 1G).^25^ Flow cytometric analysis confirmed depletion of Tregs which produced a significant decrease in the number of CD3+ T cells including both CD4+ and CD8+ subsets (STable 2, SFig 4). The proportion of CD8+ T cells expressing activation marker CD25 increased, as did the proportion expressing programmed cell death protein 1 (PD1), also upregulated on highly activated T cells. Treg depletion also resulted in a significant decrease to the number of B220+ B cells, though no changes were observed to myeloid populations (STable 3, SFig 5).

**Figure 4:**
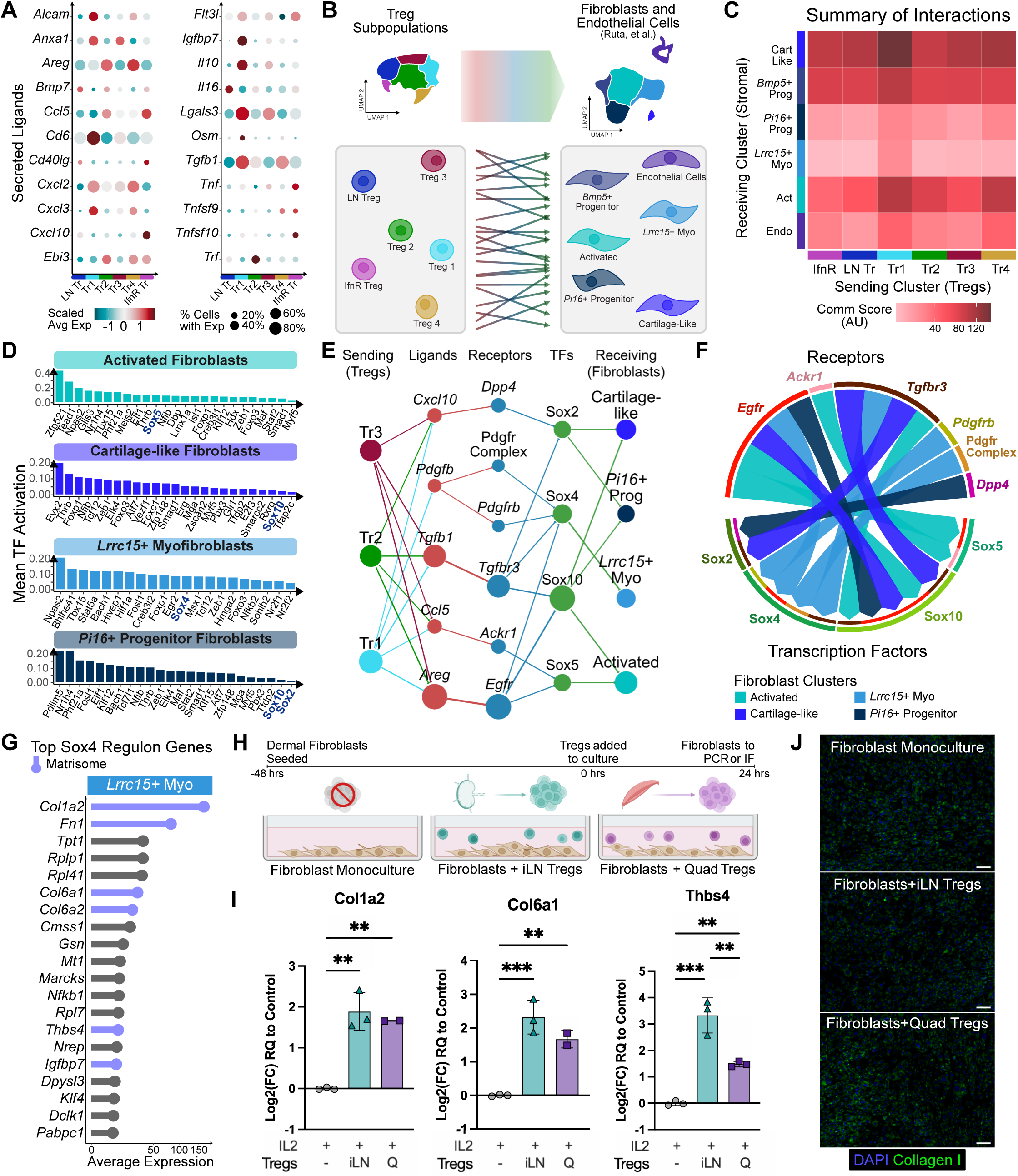
Inferred Treg activation of Sox TFs in fibroblasts aligns with increased collagen production in fibroblasts cocultured with Tregs. A) Scaled average expression (color) and percentage of expressing cells (dot size) of secreted ligands in each Treg cluster. B) Schematic of communication analysis from Tregs to fibroblasts and endothelial cells from the CD45-enriched scRNAseq data from Ruta et al. C) Communication score summarizing inferred signaling from Treg clusters to fibroblast and endothelial populations from (B). D) Activation of TFs in Treg to fibroblast network by fibroblast subcluster. E) Network plot illustrating inferred Treg to fibroblast signaling mediated by Sox family TFs with nodes representing sending clusters, ligands, receptors, TFs, and receiving clusters. Node size and edge thickness correspond to number of connections. F) Chord diagram showing connections from receptors and Sox family TFs for each fibroblast population. G) Mean expression of most highly expressed genes in Sox4 regulon in *Lrrc15*+ myofibroblast cluster, with matrisome associated genes indicated in blue. H) Schematic of fibroblast co-culture, performed with fibroblasts alone, or fibroblasts with Tregs from iLN or quadricep 6 weeks after PCL implant. I) Quantification via PCR of fibroblasts RNA after co-culture for *Col1a2*, *Col6a1*, and *Thbs4*. J) Representative images of immunofluorescence staining of DAPI (white) and collagen I (green) in fibroblast coculture wells. Scale bar = 100µm. Statistical analysis was performed for technical replicates using one-way ANOVA with Tukey’s post hoc test. p < 0.05 (*), p < 0.01 (**), p < 0.001 (***).

**Figure 5:**
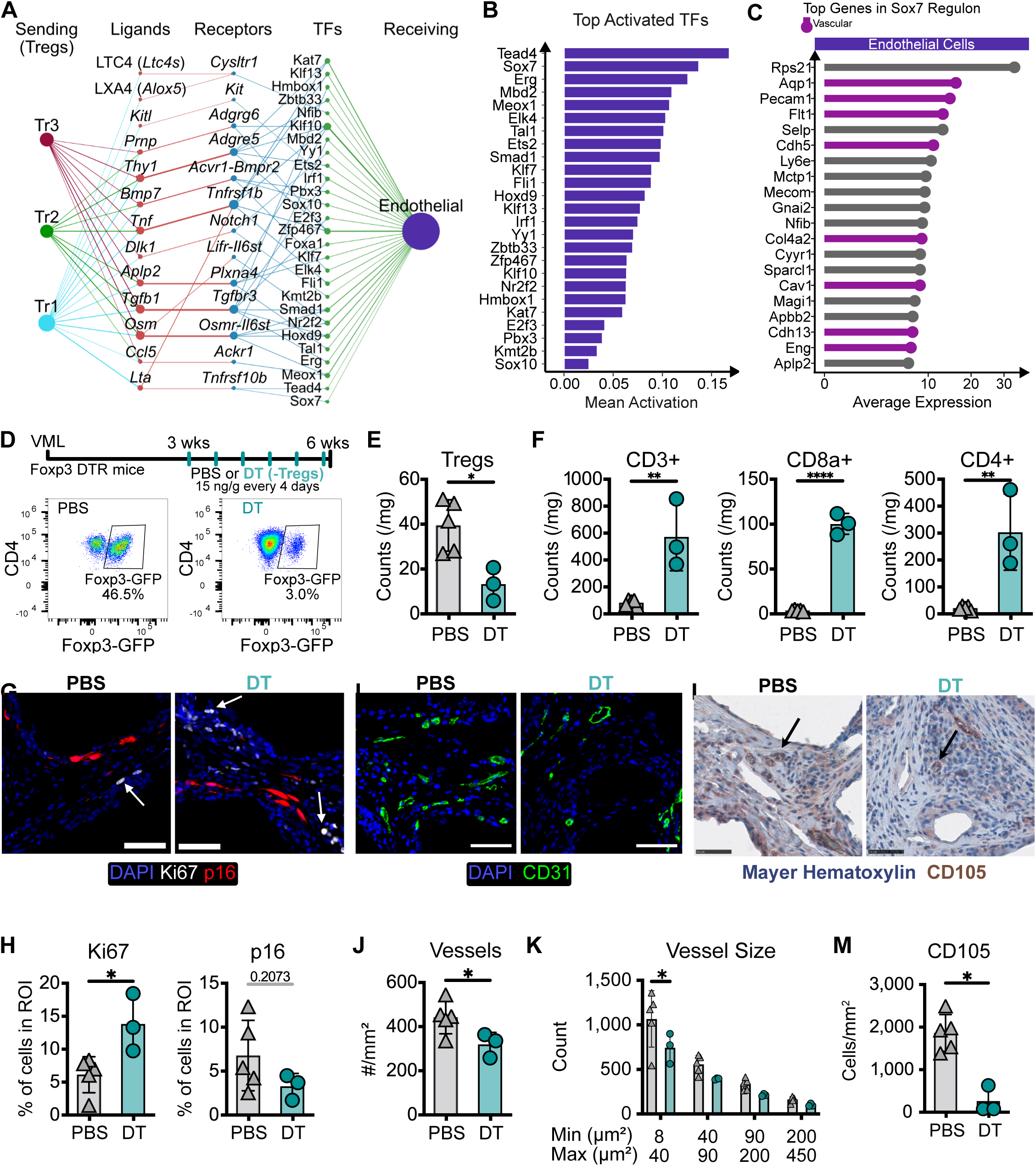
Predicted Treg to endothelial cell signaling corresponds to decreased neovascularization upon Treg depletion. A) Network plot of inferred communication from Tregs to endothelial cells with nodes representing sending clusters, ligands, receptors, TFs, and receiving clusters. Node size and edge thickness correspond to number of connections. B) Activation of TFs in in Treg to endothelial signaling network. C) Mean expression of most highly expressed genes in the endothelial cells within the Sox7 regulon, with genes associated with vasculature highlighted in purple. D) Timeline of Treg depletion model with PBS (n = 5) or DT (n = 3) administered every 4 days from 3 – 6 weeks after PCL implant (top) and representative flow cytometry gating of Tregs (bottom); percentages shown are for representative plots. E) Flow cytometry counts of Tregs in PBS and DT treated mice. F) Flow cytometry counts of CD3+ T cells and subsets (CD8+, CD4+ conventional). G) Immunofluorescence images of quads 6 weeks after PCL implant stained for DAPI (blue), Ki67 (white) and p16 (red). H) Quantification of percentage of Ki67+ cells (left) and p16+ cells (right) in regions of interest. I) Immunofluorescence staining of DAPI (blue) and CD31 (green) in PBS and DT treated mice. J) Quantification of vessel number by area K) Comparison of vessel size distribution based on CD31 staining; full histogram in SFig 26. L) Images immunohistochemistry staining for Mayer hematoxylin (blue) and CD105 (brown). M) Quantification of CD105+ cells by area. All images are representative. Scale bars = 50µm. Bar plots mean ± SD. Statistical significance was assessed with unpaired two-sided t-test (D-J, M) or two-way ANOVA with Sidak’s multiple comparison post hoc test (K): p < 0.05 (*), p < 0.01 (**).

We then examined the tissue-level consequences of the Treg depletion with bulk RNA sequencing of the fibrotic tissue. We identified 857 differentially expressed genes upon Treg depletion including decreased expression of canonical Treg-associated genes (*Foxp3*, *Ctla4*), and significant changes to genes associated with cell adhesion, immune activity, and metabolism (Fig 1H). Ranked gene set enrichment analysis revealed clear differences between the Treg-depleted and control fibrotic tissues (Fig 1I). Treg depletion resulted in significant enrichment of interferon response pathways (both alpha and gamma), consistent with the loss of immunosuppression. Additionally, in the absence of Tregs we observed enrichment of metabolic pathways (oxidative phosphorylation, fatty acid metabolism) and tissue development programs (adipogenesis, myogenesis), suggesting changes beyond immune modulation.

We next sought to determine whether Treg depletion induced changes in gene expression associated with properties of the fibrotic tissue such as ECM and vascularity. We used the Matrisome Project database to identify significantly differentially expressed ECM-related genes in the fibrotic tissue upon acute Treg depletion (Fig 1J).^26^ Treg depletion shifted the balance of ECM components. Glycoprotein *Colq*, which plays a structural role in neuromuscular junctions, increased.^27^ Other glycoproteins decreased, such as *Thbs1*, involved in matrix mechanotransduction, and *Lrg1,* which activates TGFβ signaling in skin fibroblasts to encourage ECM deposition.^28,29^ Secreted ligands also shifted with Treg depletion, with increased expression of growth factors (*Egfl6*, *Fgfpb1*, *Fgf13*) and reduced expression of ligands associated with TGFβ (*Wfikkn1*) and tumor necrosis superfamily member RANKL (*Tnfsf11*). ECM affiliated proteins and ECM regulators also decreased with Treg depletion, such as *Emcn*, which impacts cell interaction with ECM and is important for vascular function and angiogenesis.^30^ We used Gene Ontology annotations to identify genes associated with vasculature (Fig 1K).^31,32^ Vascular regulatory factor *Vegfb* was among the genes that increased with Treg depletion, while genes associated with vascular development (*Foxf1*) and endothelial migration (*Angpt2*) significantly decreased. These results suggest that Tregs contribute to formation and modulation of the ECM composition and vascularity in fibrosis.

### Tregs are activated and increase immunosuppressive phenotype during fibrosis progression

The significant accumulation of Tregs, and the impact of their acute depletion, prompted further investigation into their phenotype and stromal cell interactions. To enable a high-resolution analysis of Treg heterogeneity, we subclustered Tregs in scRNAseq data we obtained from VML-PCL quadriceps at 1-, 6-, and 12-weeks post-implant (n = 3-5 per timepoint for PCL; n = 1 for saline;, STable 4).^33^ As we described in Ruta et al (2025), after quality control excluded contaminating myeloid cells, our dataset comprised 14,995 T cells with an average of 3,553 reads and 1,850 genes detected per cell. Unsupervised clustering identified 7 distinct T cell clusters, which were annotated using established cell markers; Tregs (*Cd3e, Cd4, Foxp3),* CD4 (*Cd3e, Cd4,* no *Foxp3*), CD8 (*Cd3e, Cd8a*), GD (*Cd3e, Trdc*), two unconventional T cell clusters (*Cd3e; Zbtb16, Trdc* and/or *Klrg1*), and proliferative cells (*Mki67, Top2a*). The T cell subsets in this single-cell dataset showed a modest decrease in CD8 T cells (16% to 12%), an increase in conventional CD4 T cells (7% to 19%), and a notable expansion of Tregs (14% to 35%) from 1 to 6 weeks (SFig 6), paralleling the shifts seen by flow cytometry in this study.

To further characterize Treg phenotype and heterogeneity, we performed additional subclustering of the scRNAseq data for the Treg cluster (3,678 cells combining all time points) and identified 7 distinct subclusters (Fig 2A, SFig 7). These subclusters included populations with gene expression profiles similar to known Treg subsets, such lymph node-associated Tregs (LN Tregs, expressing *Lef1, S1pr1,* and *Klf2*; SFig 8), interferon-responsive Tregs (IfnR Tregs, expressing *Ifit1, Cxcl10,* and *Isg15),* and cycling Tregs (expressing *Mki67* and *Top2a*; Fig 2B). We also identified four effector Treg clusters (Tregs 1-4) characterized by increased expression of immunosuppressive markers. These effector Treg subclusters broadly expressed *Il10*, *Gzmb*, and *Lag3*, highlighting their shared suppressive regulatory program albeit with varying enrichment across the different clusters (Fig 2C, SFig 9). For example, Treg 1 expressed higher levels of *Pparg* (associated with potent immunosuppression), as well as *Lag3* and *Cxcr6* (associated with tissue resident memory cells) compared to the other effectors Tregs.^34–36^ Tregs 2 and 4 expressed higher levels of genes associated with early effector Treg activation, such as *Cd81, Cd83,* and *Rel*, while Treg 4 was enriched in polyamine metabolism-related genes, including *Srm,* which is implicated in T cell differentiation.^37–39^ Along with Treg 1, Treg 3 was notable for high expression of suppressive cytokine *Il10* and of *Cd69* and *Lgals3*, both of which can promote IL10 secretion in T cells, suggesting similar suppressive function at a lower intensity.^40^

We next examined markers of Treg activation and tissue residence (SFig 10A). All Treg populations broadly expressed genes associated with activation and tissue residence. The effector Tregs (Tregs 1–4) expressed higher levels of Treg activation-associated genes than conventional CD4+ T cells, but each effector subset displayed distinct enrichment patterns (Fig 2D). For example, Tregs 1 and 3 enriched *Il10* and *Havcr2* (encodes TIM3); Tregs 2–4 enriched *Ikzf2* (encodes HELIOS); and Tregs 1, 2, and 4 enriched *Pdcd1* (encodes PD1). Among these, Treg 1 expressed the highest levels of *Lag3*, *Il10*, and *Havcr2*, even compared with other effector Treg subsets. Treg clusters also differed in their expression of tissue residency markers, including *Itgae*, *Il1rl1*, and *Areg* (Fig 2E).^35^ Treg 1 showed the highest *Pparg* expression, associated with visceral adipose tissue Tregs; Tregs 1 and 3 highly expressed *Klrg1,* expressed on non-lymphoid Tregs; and Treg 2 expressed more *Gata3,* involved in Treg accumulation at sites of inflammation, and *Il1rl1* (encoding ST2, the IL-33 receptor), which is important for Treg accumulation in skeletal muscle.^41,42^ Together, these patterns suggest the four effector populations represent related regulatory states along gradients of suppressive potential, activation state, and tissue adaptation.

During fibrosis progression, effector Treg populations, specifically Tregs 1 and 3, increased in frequency and increased expression of immunosuppressive markers. These subsets expanded in frequency over time, while Treg 4 remained stable and the IfnR and cycling Tregs that predominated at 1 week declined sharply by 6 weeks (Fig 2F, SFig 10B). Within the three largest effector Treg populations (Tregs 1–3), we observed time-dependent changes in key immunoregulatory genes, including *Il10*, *Ebi3*, and *Gzmb*, as well as increased expression of immune checkpoint molecules *Pdcd1* (PD1), *Tigit*, and *Havcr2* (TIM3; Fig 2G, SFig 10C). Interestingly, the expanding Treg 1 and 3 subsets showed progressive *Il10* upregulation but reduced expression of *Ebi3* (IL-35) and *Gzmb*, indicating a shift toward IL-10–driven suppression. Immune checkpoint genes *Pdcd1*, *Tigit*, and *Havcr2* also increased over time in these subsets. Flow cytometry confirmed these changes at the protein level, with more TIGIT+ and PD1+ Tregs, and a rising fraction of cells co-expressing TIM-3, as fibrosis advanced (Fig 2H, STable 1). Together, these findings show that Tregs acquire a progressively increased suppressive phenotype during fibrosis development represented by increasing Treg effector clusters 1 and 3.

### Clonal expansion and TCR activation increase in suppressive effector Tregs in the fibrotic niche

The TCR is central to adaptive immunity, enabling T cells to recognize specific peptide antigens. To probe clonal dynamics, we performed TCR repertoire analysis using the paired scTCRseq data from Ruta et al.^33^ In the previous study, we identified γδ T cells using the expression of γ or δ TCR variable region genes. In this study, we focused on the analysis of α or β chains in other T cell subpopulations. TCR sequencing achieved a high degree of coverage, with 80% of cells assigned at least an α or β chain (SFig 11A). Clones were defined by identical V gene and CDR3 nucleotide sequences for both the α and β chains, as clonal expansion results in T cells bearing fully matched TCRs. When the TCR sequence information was incomplete, cells were grouped only with others sharing the same available sequence and pattern of missing data to prevent ambiguous or inflated clonal assignments. We first assessed clonal space homeostasis within the CD3 scRNAseq data by categorizing clones based on expansion level (single, small, medium, large, or hyperexpanded) according to clonal frequency distributions. When stratified by cluster, we found the highest number of hyperexpanded clones in Tregs compared to all other T cell populations (Fig 3A, SFig 11B). Visualization of the clones on UMAP plots, split by time point, further highlighted the accumulation of expanded Treg clones as fibrosis progressed (Fig 3B, SFig 11C).

The pattern of progressive clonal expansion in T cells during fibrosis development persists regardless of sampling depth or diversity metric. To confirm that clonal expansion was not a technical artifact from uneven sequencing coverage, we repeatedly subsampled each biological replicate to control for variation in cell numbers and TCR detection. Gini coefficient analysis revealed increasing clonality over time in all CD3+ T cells (Fig 3C), with a similar trend in Tregs despite smaller cell numbers (SFig 12A). Additional diversity measures, such as Shannon entropy and Simpson’s index, showed the same trend toward reduced repertoire diversity and greater clonal dominance, reinforcing the conclusion that clonal expansion intensifies as fibrosis develops (SFig 12B–C).

We next evaluated clonal enrichment within the Treg clusters, focusing on the 3 large effector populations (Tregs 1-3), as only a small proportion of cells were in Treg 4. Effector Treg populations Treg 1 and Treg 3 contained the largest proportions of expanded and hyperexpanded clones, while almost half the clones in Treg 2 were expanded (Fig 3D, SFig 13). To predict whether clonal expansion was accompanied by TCR signaling, we examined the expression of TCR signaling-associated genes. All three effector Treg populations expressed genes associated with receptor signaling (such as *Nfkb1*, *Rela*, *Nfatc1)* with Treg 2 expressing the highest levels (Fig 3E, SFig 14A). We next applied single-cell regulatory network inference and clustering (SCENIC)^43^ to infer TF activation scores for TFs linked to TCR signaling, including *Nfatc1*, *Nfkb1*, and *Rel*. All effector Tregs showed activation of TFs associated with TCR signaling, with Treg 2 exhibiting the highest relative TF activation of the effector Tregs (Fig 3F, SFig 14B). Together, these results support an antigen-specific Treg response in the fibrotic environment in the absence of a protein antigen in the biomaterial implant.

### Integrated clonal and transcriptional analyses identifies trajectory to suppressive Tregs

To identify possible cell trajectories, we compared patterns of clonal sharing with gene expression profiles across Treg clusters. Treg 2, marked by moderate clonal expansion, expressed early activation markers, while Tregs 1 and 3, with high proportions of expanded clones, displayed more suppressive phenotypes characteristic of differentiated cells. To explore the connection between these populations, we visualized the distribution of the 5 largest clones by size across subclusters and timepoints (SFig 14C). At 6 weeks, when fibrosis is well developed, Treg 2 predominantly contained smaller clonal populations, while larger clonal populations were found within Tregs 1 and 3 (Fig 3G). This apparent gradient of clonal expansion prompted an analysis of clonal sharing. We identified TCR sequences shared across at least two Treg subsets and quantified the cells with that sequence in each subset. Hierarchical clustering based on these cell numbers grouped Tregs 1 and 3 together, indicating a highly similar incidence of shared clones between the populations (SFig 15A). We identified 32 clones present in all three large effector Treg populations (SFig 15B). Half of these clones were more abundant in Treg 1 and 3, with the remainder either biased toward Treg 2 or evenly distributed across subclusters. The presence of overlapping TCRs among distinct transcriptional clusters suggested a trajectory towards a highly differentiated Treg phenotype.

Given the presence of shared clones and the activation of suppressive gene programs that increased over time, we next examined whether these Treg populations existed along a continuum of differentiation states. To assess phenotypic transitions, we performed pseudotime analysis of all Tregs using the comparatively undifferentiated LN Tregs as the root population. Visualization in principal component space confirmed separation of the Treg subpopulations by transcriptional state (Fig 3H). Trajectory inference identified 4 paths, one of which aligned with the observed patterns of clonal expansion and differentiation status (SFig 16). Coloring cells by pseudotime along this trajectory highlighted the progression of cellular states independent of cluster boundaries (Fig 3I). Overlaying cells on the principal curves, we observed an increase in pseudotime from the LN Tregs through Treg 2 and Treg 3 to Treg 1 (Fig 3J). When cells were stratified by clonal expansion status along the predicted trajectory, cells with higher degrees of expansion also had greater pseudotime values, supporting a link between clonal expansion and progressive differentiation (Fig 3K, SFig 17).

We next sought to further define the relationship between Treg phenotype, clonal status, and transcriptional dynamics along the inferred trajectory. We visualized scaled expression of key pseudotime-associated genes (p_adj_ < 0.05), ordering cells by pseudotime and annotating cluster identity, clonal expansion category, and pseudotime value (Fig 3L). This aligns the changing gene expression patterns with cell differentiation and clonality. Genes associated with lymphoid Tregs (*Nrp, S1pr1*) were most highly expressed at the start of the trajectory (LN Tregs). Early activation genes (*Cd81, Cd74)* emerged mid-trajectory (primarily in Treg 2), and effector-suppressive markers (*Gzmb, Il10*) peaked at trajectory end, coinciding with greater clonal expansion and spanning effector populations Tregs 1 and 3. Overall, the integrated clonal and transcriptional analyses reveal that Treg clonal expansion is closely associated with progressive differentiation and the acquisition of suppressive phenotypes during fibrosis

### Cell communication inference identifies pathways for Treg interactions with fibroblasts and endothelial cells

Given the accumulation of clonally expanded, immunosuppressive Tregs during fibrosis, we next sought to define the pathways through which these cells may interact with the stromal cells responsible for the altered ECM and vascular tissue structure in fibrosis, specifically fibroblasts and endothelial cells. To this end, we applied ligand-receptor cell communication inference of the scRNAseq dataset to examine Treg transcriptomic profiles in conjunction with stromal cells in the fibrotic microenvironment.

We first profiled the secreted ligands expressed by Tregs across all time points using CellPhoneDB.^44^ After filtering for ligands expressed by at least 5% of cells in any cluster, we identified 22 molecules (Fig 4A, SFig 18). Canonical Treg ligands, such as *Areg* and *Tgfb1*, were broadly expressed across all Treg subtypes. In addition, each effector Treg population displayed a distinct ligand profile. For example, Treg 1 expressed *Igfbp7* and *Osm*, which are linked to fibrogenesis; Treg 2 expressed *Ebi3* (a subunit of IL-35), known to alleviate lung fibrosis, and *Trf,* which can serve as a biomarker for liver fibrosis; and Treg 3 expressed high levels of the suppressive molecules *Il10* and *Lgals3*.^45,46^ To predict Treg cell communication networks with the fibroblasts and endothelial cells, we focused on the primary effector Treg populations (Tregs 1-3) at the 6 week timepoint when fibrosis is well established.

To create cell networks from Treg-generated ligands, we performed further analysis in conjunction with the additional, independent CD45-enriched scRNAseq dataset from fibrotic tissue 6 weeks post-injury and PCL implantation from Ruta et al.^33^ As previously described, clustering revealed multiple stromal and immune subpopulations, including fibroblasts (*Col1a1, Col3a1, Dcn*), endothelial cells (*Pecam1, Flt1, Cdh5*), lymphatic endothelial cells (*Lyve1*, *Ccl21a, Prox1*), perivascular cells (*Rgs5*, *Myh11, Tagln*), myeloid cells (*Itgam, Csf1r, Cd68*) and T and natural killer cells (*Cd3e, Nkg7*). Fibroblasts were subclustered to resolve heterogeneity and communication with subpopulations (Fig 4B). These subclusters included two progenitor subpopulations, *Pi16*+ (*Pi16*, *Dpp4*) and *Bmp5*+ (*Bmp5*, *Bmp4, Penk*), which resemble the stem and fibroadipogenic progenitors previously reported in fibroblasts and murine skeletal muscle.^47,48^ The other subclusters included activated fibroblasts, which were delineated by their expression of inflammatory, Wnt signaling, and matrix remodeling genes (*Sfrp1*, *Sfrp2*, *Mmp3*), as well as terminally differentiated *Lrrc15*+ myofibroblasts (*Acta2, Lrrc15, Tnc*) and cartilage-like fibroblasts (*Tnmd, Comp, Fmod*).^49–51^

To estimate signaling pathways between Tregs and the fibroblast and endothelial populations in this current study, we employed dominoSignal, a bioinformatics analysis pipeline that goes beyond simple ligand-receptor mapping by integrating ligand expression in sender cells, cognate receptor expression in receiver cells, and downstream activation of TFs based on inferred regulons.^52,53^ For each interaction, we calculated a composite communication score reflecting ligand, receptor, and TF components. We aggregated scores across all predicted interactions between each Treg subset and each stromal population as a quantitative overview of potential intercellular signaling strength (Fig 4C). We found that Treg 1, the most clonally expanded and phenotypically suppressive population, sent the strongest collective signals to fibroblasts and endothelial cells. On the receiving side, cartilage-like fibroblasts exhibited the highest combined communication score, suggesting increased connection from Tregs to this population.

### Treg signaling activates multiple transcription factors including Sox family in fibroblasts

To resolve individual signaling interactions driving fibroblast responses, we constructed an inferred Treg to fibroblast communication network. Across all Treg subsets, this network included 15 unique ligands, 19 unique receptors, and 78 unique TFs (SFig 19). In these networks, nodes correspond to cell types and edges to signaling interactions. Their sizes and thicknesses scale with the number of pathways involved, emphasizing key mediators of predicted Treg to fibroblast communication, such as *Areg* to *Egfr* and *Tgfb1* to *Tgfbr3*, both implicated in fibrotic signaling cascades.^54–56^

Examining downstream consequences of the predicted communication pathways, we analyzed the top 25 TFs activated in each receiving fibroblast cluster (Fig 4D). We found Treg-induced activation of TFs including *Jun*, *Hif1a*, and *Tead1* in the fibroblast subclusters, all of which have been linked to fibrosis pathogenesis.^57–59^ Furthermore, there was activation of multiple members of the activator protein 1 (AP-1), Kruppel-like factor (KLF), and SRY-related HMG-box (Sox) families of TFs, which have established or emerging roles in fibrotic remodeling.^60–62^ Though many TFs were inferred in the signaling network, the frequency and breadth of the Sox family was notable. Sox TFs were involved in 6.8% of predicted interactions with the major Treg effector populations and were activated by effector Tregs 1-3 across four of the five fibroblast clusters.

While Sox family members are recognized as contributors to fibrosis, and can be induced by TGFβ, the potential for Treg-mediated Sox activation in stromal cell types has not been established in fibrotic disease contexts, motivating further analysis of Sox-associated pathways. To further probe the contributions of Sox family TFs, we focused on interactions predicted between the three largest effector Treg subsets (Tregs 1-3) and fibroblasts. Central predicted ligand-receptor pairs such as *Areg* to *Egfr* and *Tgfb1* to *Tgfbr3* remained prominent, alongside additional interactions including *Ccl5* to *Ackr1* and *Pdgfb*-Pdgfr complex (Fig 4E, SFig 20A).

Mapping predicted receptor-Sox TF activation at the fibroblast subcluster level revealed differences in subtypes of Sox (Fig 4F). Sox4 activation via Treg signaling through multiple receptors was specific to *Lrrc15+* myofibroblasts, while Sox5 was selectively activated in the activated fibroblast population. Sox10 activation was restricted to receptors *Egfr* and *Tgfbr3* but was present across activated fibroblasts, cartilage-like fibroblasts, and *Pi16*+ progenitor fibroblasts. Interestingly, Sox2 activation in *Pi16+* progenitor fibroblasts was linked to receptor *Dpp4. Pi16* and *Dpp4* are linked to progenitor fibroblasts across multiple organs;^63^ however, *Dpp4* is also a marker of FAPs in skeletal muscle that may promote fibroblast activation when overexpressed, and its inhibition in systemic sclerosis may reduce fibrosis.^64,65^ In the cartilage-like fibroblasts, Sox2 activation was associated with TGFβ signaling. There was no evidence of Sox TF activation by the large Treg effector populations in the *Bmp5+* progenitor fibroblasts.

### Treg communication with fibroblasts promotes collagen production

To investigate potential downstream consequences of Sox-family TF activation, we analyzed the inferred regulons driving activation in each fibroblast population, leveraging the matrisome database to annotate genes associated with ECM.^26^ In the *Lrrc15+* myofibroblasts, Sox4 activation scores were driven by expression of ECM-associated genes such as *Col1a2, Fn1,* and *Col6a1*, along with other fibrosis modulators such as *Nfkb1* and *Klf4,* consistent with their role in ECM deposition and drivers of fibrosis (Fig 4G).^66–68^. Similar analysis revealed that Sox5 in activated fibroblasts was driven by *Fn1*, while highly expressed Sox10 target genes included proteoglycan *Gpc3* (in both activated and *Pi16+* progenitor fibroblasts), and glycoprotein *Chrdl1* (in the cartilage-like fibroblasts). Activation of Sox2 in both cartilage-like and *Pi16+* progenitor fibroblasts was associated with high expression of genes such as *Crispld1* and *Sema3b*, which are implicated as a biomarker and fibrosis inhibitor in IPF, respectively (SFig 20B-D).^69,70^ These data suggest a direct mechanism by which Treg signals may influence fibroblast ECM production, with strong expression of matrix associated genes present in the Sox4 regulon of myofibroblasts.

We used an *in vitro* Treg-fibroblast coculture model to experimentally validate predicted Treg influence on fibroblasts. Tregs isolated from the quadricep and inguinal lymph nodes (iLN) of mice 6 weeks after PCL implantation were cultured with dermal fibroblasts seeded 48 hours prior. A fibroblast monoculture served as a control (Fig 4H). Quantitative RT-PCR revealed significant increases in expression of *Col1a2*, *Col6a1*, and *Thbs4* in fibroblasts cocultured with Tregs compared to monoculture. These genes were identified as leading genes in the Sox4 regulon in myofibroblasts (Fig 4I). Inguinal LN Tregs stimulated higher *Thbs4* expression levels than quadricep Tregs. Immunofluorescence imaging of Collagen I confirms fibroblast morphology and collagen protein production (Fig 4J, SFig 21).

### Tregs are predicted to communicate with endothelial cells

We extended our Treg communication analysis to endothelial cells, given the established link between fibrosis and altered tissue vasculature.^71,72^ The inferred network comprised of 182 distinct signaling pathways, suggesting multiple routes through which Treg-derived signals might influence endothelial cells and subsequent vascular structure (Fig 5A, SFig 22). Included among these pathways were well-characterized regulators of endothelial function in fibrosis, such as *Tgfb1* to *Tgfbr3*.^73,74^ The network also included signaling through lipid mediators leukotriene C4 and lipoxin A4 to *Cysltr1*, associated with endothelial cell proliferation and *Dlk1*-*Notch1*, an interaction that promotes neovascularization.^75,76^ The predicted interactions also encompassed signals involved in endothelial cell inflammation, such as *Tnf*-*Tnfrsf1b* (TNFα-TNFR2) and *Osm*-Osm receptor complex, indicating Tregs may impact both vascular development and inflammation.

Downstream of the ligand-receptor signaling, we analyzed the most highly activated TFs in endothelial cells. Activated TFs included members of the KLF and Sox families, which have been linked to vascular health and development (Fig 5B).^77,78^ Other highly activated TFs included Tead4, which integrates mechanical cues for vessel growth in conjunction with YAP/TAZ, and Erg, linked to regulation of endothelial homeostasis and vascular development.^79,80^ The Tead4 regulon is associated with communication from both gamma delta T cells and Tregs as we described previously, and has high expression of genes essential for vascular development and structural integrity, such as *Pecam1* (CD31), *Flt1* (VEGFR), and *Cdh5* (VE-Cadherin).^33,81–83^ Sox7, most highly activated after Tead4, similarly regulated key endothelial genes involved in vascular development and differentiation. Its regulon, like that of Tead4, encompassed *Pecam1*, *Flt1*, and *Cdh5*.^78,84^ It also included genes such as *Col4a1*, a basement membrane collagen; *Cav1*, an inhibitor of endothelial cell permeability; and *Eng*, which plays a crucial role in angiogenesis (Fig 5C).^85–87^ These predicted signaling pathways suggest that Treg-derived signals promote activation of vascular development and homeostasis programs.

### Treg depletion reduces neovasculogenesis in the fibrotic tissue environment

To further characterize the impact of Tregs in fibrosis development and validate the stromal effects predicted from the scRNAseq data, we performed a longer-term depletion. Previous studies identified 3 weeks as a critical transition when the muscle injury heals and biomaterial-induced inflammation, senescence and fibrosis increase.^12^ We therefore started Treg reduction 3 weeks after material implantation using the Foxp3-DTR mice and administered low dose diphtheria toxin (DT) every 4 days for 3 weeks (until 6 weeks harvest; Fig 5D, SFig 23A-D). This low dose regimen was previously reported to reduce Treg numbers while avoiding systemic complications caused by their depletion.^88^

Flow cytometric analysis confirmed a significant decrease in absolute Treg counts as well as a decrease in the proportion of Tregs among CD3+ cells (Fig 5E, SFig 23E-H, STable 5). Chronic Treg reduction led to significant increases in the total number of T cells in both the CD8+ and CD4+ compartments (Fig 5F). We observed changes in other immune populations, including decreases in monocytes and natural killer cells and increased eosinophils. (SFig 23I-M). In addition, macrophages decreased as a proportion of immune cells (62% ± 2.2% to 52% ± 4.7%). The bulk macrophage population displayed increased median fluorescence intensity (MFI) of CD206 and MHCII with Treg depletion, while CD86 MFI decreased (SFig 24A). However, within a subset of CD206+CD301b+ macrophages, often associated with repair, CD206 and CD86 both decreased with Treg depletion (SFig 24B).^89^ Because this subset represented only a minor fraction of macrophages, it did not alter the population level trend of increased CD206 MFI and MHCII MFI along with decreased CD86 MFI (SFig 24C-D).

Beyond changes to immune cell composition, we examined the impact of Treg depletion on the overall tissue. As cellular senescence plays a known role in fibrosis, we assessed markers of proliferation (Ki67) and senescence (p16).^90^ Immunohistochemical analysis revealed significantly increased Ki67+ cells in Treg-depleted tissues compared to controls, with a decreasing trend in p16+ cells (Fig 5G-H, SFig 25, STable 6). This shift between the balance of proliferation and senescence, alongside the altered immune populations, may influence the progression and structure of the fibrotic response.

To directly assess Treg influence on tissue structure, we next evaluated vascular changes in the tissue. Immunostaining for the endothelial marker CD31 revealed altered vascular architecture following Treg depletion (Fig 5I, SFig 26, STable 6). The vessel density was significantly reduced, with a significant decrease in the smaller annotated structures (Fig 5J-K). There was no difference in larger vessels. Further assessment of vasculogenesis using CD105 (Endoglin, encoded by *Eng*) immunohistochemistry showed a significant decrease in CD105+ cells in Treg-depleted tissues, indicating a reduction in neovascularization in the absence of Tregs (Fig 5L-M, SFig 27). These functional studies support the role of Tregs on the neovascular response to fibrosis previously inferred from our scRNAseq data.

## Discussion

This study demonstrates that Tregs play a role in shaping the fibrotic microenvironment in a material-induced model of fibrosis. We find that Tregs within fibrotic lesions are phenotypically diverse, clonally expanded, and exert influence over matrix composition and vascular architecture. Accumulating Tregs exhibit an activated, tissue-resident phenotype, characterized by distinct transcriptional profiles, including interferon response, metabolic activity, and effector function. Suppressive Tregs undergo clonal expansion and express genes involved in TCR signaling, consistent with the potential for antigen-specific responses. Temporal clonal analysis reveals a progressive shift from LN associated gene expression, through TCR activation, and culminating in a suppressive effector phenotype. Short-term Treg depletion altered the expression of genes related to ECM and vasculature, while sustained depletion modulated immune cell composition, increased inflammation, and significantly reduced neovascularization. Fibroblast coculture with Tregs resulted in increased expression of ECM associated genes. Collectively, our work positions Tregs as an important population in influencing both immune and stromal compartments, thereby shaping the structure of fibrotic tissue.

These findings build on our recent analysis of γδ T cells from the same dataset, which characterized type-1 and type-17 γδ T cell subsets and highlighted the association of type-17 γδ T cells with chronic inflammation. The current work focuses on the Treg compartment, a population with a distinct TCR repertoire that is broadly implicated in modulating inflammatory responses. By examining this separate immune axis, we identify a regulatory layer of adaptive immunity that shapes the tissue environment surrounding synthetic implants. While both studies point to a common theme in which T cell subsets influence ECM composition and vascular features of fibrotic tissues, this parallel reveal emphasizes the role of T cell to stromal interactions within the fibrotic niche. Recognizing these complementary roles highlights the broader range of T cell driven pathways in the foreign body response and motivates the Treg focused analysis in this study.

A key finding of this work is the progressive accumulation of Tregs with an activated phenotype, which ultimately dominate the T cell compartment in the fibrotic lesion. This is consistent with observations from other biomaterial-induced fibrosis models, where Treg abundance has been correlated with reduced capsular contracture symptoms in silicone mammary implants and with Tregs and T helper cells present at levels comparable to macrophages in synthetic abdominal mesh implants.^91,92^ These accumulating Tregs display a tissue-resident phenotype, evidenced by expression of genes such as *Areg, Il1rl1,* and *Pparg*, that aligns with tissue-resident Tregs described in muscle and visceral adipose tissue.^42^ This suggests either tissue-resident Tregs expand in response to injury or circulating Tregs recruited to the lesion adopt a tissue-adapted profile within the fibrotic environment. The universal activation state we observe likely reflects the ongoing inflammation characteristic of the chronic response to the implanted material. While Tregs in organ-specific fibrotic diseases results in varied outcomes, ranging from anti-fibrotic effect in hepatitis B-related liver fibrosis to promotion of fibrosis via *Areg* secretion in non-alcoholic steatohepatitis, our work adds to the growing consensus that a robust, functionally engaged Treg population can influence fibrotic tissue structure.^93,94^

An unexpected observation is the clonal expansion of Tregs in response to implantation of a synthetic material lacking obvious peptide antigen. This complements previous work demonstrating CD4+ T cell IL17 production in response to synthetic implants is dependent on TCR repertoire diversity.^12^ Similarly, scRNAseq analysis from synthetic polymer implants revealed elevated expression of T cell activation and TCR signaling genes.^95^ Beyond biomaterial-induced responses, this parallels observations in a murine model of acute muscle injury, where injury associated Tregs show clonal expansion, and adoptively transferred Tregs bearing specific TCRs preferentially accumulate at the site of injury.^21,22^ In our model, greater clonal expansion is associated with a suppressive Treg phenotype, in line with previous evidence linking TCR engagement to the differentiation and function of suppressive Tregs.^96,97^ The specificity of the antigens driving this process remains unclear. One possibility is recognition of self-peptides or damage-associated molecular patterns exposed by the tissue injury, or even neo-epitopes generated at the implant surface. This may frame the response in a similar context to autoimmunity, where Treg dysfunction in response to self-antigens drives pathology. The emergence of cartilage-like fibroblasts in the fibrotic lesion raises the further possibility that immune-privileged ECM components present novel antigens for adaptive immune recognition.^98,99^ Alternatively, bystander activation in the setting of persistent inflammation and tissue remodeling cannot be ruled out.

Our data support a model in which Tregs orchestrate crosstalk to influence multiple cellular compartments. Inferred communication networks between Tregs and fibroblasts via pathways including the *Areg*-*Egfr* axis and the *Tgfb1-Tgfbr3* axis align with known mechanisms of fibrosis.^54,74,94^ Extending these findings, our analysis identifies the Sox family of TFs as possible downstream effectors. Sox2 serves as a non-canonical regulator of collagen deposition, and Sox4 has been suggested as a therapeutic target for cardiac hypertrophy, indicating that these TFs contribute to fibroblast function in fibrosis.^100,101^ Sox5 is known for its role in cartilage development, which is relevant given our identification of a cartilage-like fibroblast phenotype in this niche.^100–102^ The increased gene expression of collagens by fibroblasts cultured with Tregs as compared to a fibroblast monoculture validate the impact of Treg signaling on transcriptional profile of fibroblasts. These signaling pathways may be conserved in other contexts where immune-stromal interactions drive tissue remodeling.

Our scRNAseq analyses further predict Treg communication with endothelial cells, activating TFs including Tead4, associated with mechanical signaling to vessels, Sox7, involved in early angiogenesis, and Erg, linked to vascular homeostasis and development.^79,84,103^ This inferred Treg influence on vascular development is supported by the significantly reduced neovasculogenesis observed upon Treg depletion. In a murine model of lung adenocarcinoma, Treg depletion resulted in increased expression of genes related to vascularization, a contrast to our results, emphasizing the context-dependent nature of Treg activity.^20^ While the reduced microvessel density in our model of fibrosis may reflect either impaired reparative angiogenesis or prevention of aberrant leaky vasculature, our data establish Tregs as a determinant of vascular structure in fibrosis. These targeted communication pathways likely work in concert with broader mechanisms, including metabolic and developmental programs suggested by our bulk transcriptomic data, to influence the tissue environment. This vascular role is particularly relevant for aging, where impaired angiogenesis and chronic inflammation contribute to tissue dysfunction.

Despite these insights, our study does have limitations. Single-cell transcriptomic approaches may underrepresent rare cell types, and sequencing depth constrains the detection of low-frequency TCR clones. Our cell-cell communication analysis is based on ligand-receptor-transcription factor network inference of transcriptomic data; while immunofluorescence staining and flow cytometric analysis serve as evidence of protein expression, additional study at the protein and functional levels is required. Finally, while the biomaterial-induced model provides a robust and reproducible platform for exploring fibrotic mechanisms, the translation of these findings to human fibrotic diseases, whether systemic or organ-specific fibrotic syndromes, requires further investigation.

Several important questions remain for future research. The antigens driving Treg clonal expansion remain undefined, with no public clones detected, suggesting a need for increased sequencing depth or systemic TCR analyses for further elucidation. Comprehensive TCR sequencing could define the diversity and tissue-restriction of the Treg repertoire over time. Advanced motif analysis may reveal shared predispositions for survival or selection of TCR clones within the fibrotic niche. Further characterization of Treg-mediated signaling, especially its effects on compartments such as myeloid populations, will be critical for understanding how Tregs might indirectly influence the fibrotic stroma. Importantly, we find that broad depletion of Tregs is unlikely to improve outcomes, as Treg loss exacerbates inflammation without clear reductions in the developing fibrosis despite the changes to ECM and vasculature. This suggests a need for a nuanced approach to therapeutics, where approaches targeting specific Treg functions or communication pathways, rather than enhancement or depletion, may prove more beneficial.

Collectively, these results underscore the importance of Tregs as not only modulators of the immune system but of immune-stromal networks impacting fibrotic tissue structure. Our study establishes the impact of these cells on matrix composition and vasculature in biomaterial induced fibrosis, providing a greater understanding of the role of adaptive immunity in shaping fibrotic microenvironments that may serve to inform development of therapeutics for pathological fibrotic disease contexts.

## Materials and Methods

### Animal Welfare

All animals were housed in a pathogen-free facility, and all procedures were performed in accordance with protocols approved by the Johns Hopkins University Animal Care and Use Committee. Female 6-8-week of age were used from the following strains: C57BL6/J (Jackson Laboratory, strain 000664), FoxP3-DTR (Jackson Laboratory, strain 016958), and FoxP3-GFP (Jackson Laboratory, strain 006772).

### Volumetric Muscle Loss and Implantation

The VML injury was performed as previously described and done bilaterally.^104^ Mice received 5 mg/kg of carprofen (Rimadyl 54771-8507-1 or OstiFen 510510) analgesic and were anesthetized with 2-3% isofluorane for the surgery. An incision was made in the epidermis, dermis, and underlying fascia above the muscle, approximately 3 mm^3^ segment of the quadricep was resected and the remaining space filled with approximately 25-30 mg of PCL particulate (Polysciences 25090) or 50 µL of saline. The wound was then sutured closed with 5-0 nylon sutures. For mice receiving DT therapy, DT (Sigma) resuspended as recommended and under sterile conditions was injected for a short course treatment at 50 ng/g at days –3 and –1 to tissue collection and for long-course treatment 15 ng/g every 4 days from week 3 to week 6 as previously described.^88^ Control mice were treated with PBS, all mice maintained satisfactory weight relative to starting weight up to euthanasia, however, 2 mice in the DT group were euthanized for non-experimental reasons. Mice were euthanized by isoflurane overdose at the specified time intervals post-surgery.

### Tissue Dissociation

Quadriceps femoris were collected, finely diced into ∼1 mm pieces and digested for 45 minutes at 37°C with 1.67 Wünsch U/mL Liberase TL (Roche Diagnostics) and 0.2 mg/mL DNase I (Roche Diagnostics) in RPMI 1640 medium (Gibco) supplemented with 12 mM HEPES (Gibco). Digested tissues were neutralized with cold 1% bovine serum albumin (BSA; Sigma) in RPMI, then ground through a 70 µm strainer (Falcon) and rinsed with PBS (Gibco). For pan-immune staining the cells were further filtered in a 40 µm strainer before washing with 1x PBS before plating in a 96 well plate for RBC lysis (BD). For the T cell focused panels (Checkpoint and Treg) lymphoid cells were concentrated using a percoll gradient. Cells were resuspended in 80% percoll (percoll [GE Healthcare] diluted with PBS) then 40% percoll (diluted in RPMI) and 20% percoll (diluted in PBS) were carefully layered on top then centrifuged at 2100xg for 30 minutes. Cells were collected from the 80-40 interface and washed in PBS before plating in a 96 well plate for staining. Inguinal lymph nodes (iLN) were mechanically dissociated with the end of a 1mL syringe against a 70 µm strainer (Miltenyi) directly into PBS, washed with PBS, and plated in a 96 well plate.

### Flow Cytometry

All panels’ antibody dilutions and fixatives can be found in STables 1-5 and 10. After isolation cells were stained with Zombie NIR for 30 min on ice, washed twice with staining buffer (1% w/v BSA + 1mM EDTA in PBS) and stained for 45 minutes with surface antibodies. For surface only panels cells were fixed with Fluorofix (BioLegend) for 15 min, washed and stored in staining buffer at 4°C. For TF staining, the Tru-Nuclear Transcription Staining Buffer Kit (BioLegend) was used according to manufacturer protocols. Briefly, cells were fixed for 1 hour at room temperature, washed 3x in perm buffer, stained for 45 minutes, washed 2x in perm buffer and 1x in staining buffer before storing at 4°C. Samples were run within 24 hours of staining and buffer was exchanged to PBS before data were collected on a 4L (V, B, YG, R) Cytek Aurora. Data were unmixed in SpectroFlow (Cytek) before analysis in FlowJo (FlowJo LLC).

### Histology

Tissues were collected and fixed in 10% neutral buffered formalin (Sigma-Aldrich) for 24-48 hours before a graded ethanol dehydration 70%, 80%, 95%, 100%, cleared for 1.5 hours in Xylenes (Fisher), and stored in melted paraffin (Tissue-Tek) overnight at ∼60°C. Tissues were embedded in paraffin blocks the following day. Samples were sectioned at 7 µm through the Johns Hopkins Oncology Tissue Services SKCCC core facility and one slide per sample was stained for Masson’s Trichrome by the core facility. Masson’s trichrome was imaged using a digital slide scanner (Hamamatsu NanoZoomer-XR).

### Immunofluorescence

Slides were deparaffinized in xylenes and rehydrated in ethanol concentrations 100%, 95%, 80%, 70% followed by Type 1 water. Heat-induced epitope retrieval was performed using AR6 (Akoya Biosciences) in a steamer for 15 min. 3% H2O2 (Sigma) was used to quench endogenous peroxidase for 15 min. Slides were blocked in 10% BSA with 0.05% Tween 20 in PBS for 30 minutes. Opal was used for the secondary antibodies, so Opal manufacturer protocols were used. Briefly, primary antibodies (STable 6) were incubated for 30 min at room temperature, washed in 1x TBST. Slides were then incubated with horseradish peroxidase (HRP) polymer-conjugated secondary for rabbit-on-mouse IgG (Biocare Medical) was used for 30 minutes, rinsed in TBST before reacting with tyramide signal amplification reagents: Opal 520 (Akoya), Opal 570 (Akoya), Opal 650 (Akoya) for 10 minutes. Subsequently samples were stripped by heat (steamer) with AR6 for 15 minutes before staining with the next primary antibody and Opal dye. Nuclei were counterstained with 4’,6-diami-dino-2-phenylindole (Spectral DAPI, Akoya) for 5 min before mounting cover slips using DAKO mounting medium (Agilent). Imaging was performed on an Axio Imager.A2 (Carl Zeiss Microscopy, LLC) and Zeiss Zen software. Images were linearly contrasted with isotype or primary delete controls for each channel setting the contrast levels. Images were quantified using QuPath-0.5.1. To only quantify within the FBR, an ROI was drawn around the PCL particulate without the quadricep, a mask was created on the DAPI channel using a pixel classifier (settings were adjusted to outline just the tissue without the empty PCL area), cells were identified with Cell Detection, positive cell detection was quantified using an object classifier and vessels sizes were quantified using a pixel classifier (minimum area was set to 8 µm^2^ to capture the smallest potential capillaries.

### Dermal Fibroblast Isolation

3– to 5-week-old mice (C57BL/6) were euthanized and their backs shaved and depilated with Nair for 1 min. Nair was removed with gauze and 70% ethanol. The mouse was sterilized using 70% ethanol and transferred into a biosafety cabinet where a dry 1×2 cm section of skin was cut, avoiding the peritoneum and adipose. The tissues were serially washed in RPMI-1640 medium with L-Glutamine (Gibco), then 1x DPBS (Gibco). They were transferred to RPMI to be finely diced, then enzymatically digested with 10 mL of 0.05% (w/v) Liberase TM (Roche Diagnostics) in RPMI. Any pellet formation was disrupted and the samples laid flat on a shaker set to 100-200rpm at 37°C for 45min. Pellets were vigorously disrupted again and diluted in complete RPMI (cRPMI; 10% FBS [Gibco], 1% Pen Strep [Gibco], 1% Sodium Pyruvate 100x [Gibco]) before centrifuging at 300xg for 5 min. The supernatants were aspirated and pellets resuspended vigorously in cRPMI. The skin solutions were added to T-175 flasks equilibrated with cRPMI, then shaken gently to distribute cells evenly prior to a 4-day incubation at 37°C. Then, the flasks were washed with DPBS and media switched to complete MEM (cMEM; MEM with L-Glutamine [Gibco], 10% FBS, 1% Pen Strep) to select for fibroblasts.

### Treg-Fibroblast Co-Culture

48 hours prior to the Treg sort, dermal fibroblasts were seeded into one 24-well plate (Falcon, 353047) at an initial density of 50,000 cells per well and five 8-well culture slides (Ibidi, 80841) at an initial density of 3000 cells per well in cMEM. Inguinal lymph nodes and quadriceps femoris were harvested from 6-week VML-PCL mice (FoxP3-GFP) processed and stained in accordance with the flow cytometry procedure detailed previously, and Tregs were sorted into 1% BSA in PBS (BD Fusion Sorter; Live ➜ CD45^+^ ➜ CD3^+^SSC^low^ ➜ Foxp3^+^). Tregs were added to fibroblasts at a ratio of 1:100 assuming fibroblasts were confluent. All wells received the same base media of cMEM supplemented with 10 ng/mL IL2 (BioLegend, 575402). Fibroblasts were either untreated, treated with inguinal lymph node Tregs, or treated with quadricep Tregs. They were incubated for 24 hours at 37°C then washed 3x with DPBS. The 24-well plate was dissociated in TRIzol (Invitrogen) for 5 min before transfer to Eppendorf tubes on dry ice and long-term storage in –80°C for RNA extraction. The culture slides were fixed with 4% paraformaldehyde (Santa Cruz, sc-281692) and stored in DPBS at 4°C for immunofluorescence staining.

### RT-qPCR

RNA was extracted from samples stored in TRIzol using chloroform (Sigma, C2432) followed by isolation using the RNeasy Plus Micro Kit (Qiagen) according to manufacturer instructions and quantified with a NanoDrop 2000 (ThermoFisher Scientific). RNA was converted into cDNA using SuperScript IV VILO Master Mix (ThermoFisher Scientific) and a C100 Touch Thermocycler (BioRad). For each sample, 100 ng of cDNA was plated per well and RT-qPCR was performed using the StepOne Plus Real-Time PCR System (Applied Biosystems) with TaqMan Gene Expression Master Mix (Applied Biosystems) according to manufacturer instructions. Murine TaqMan gene expression probes were used: Rer1 (Mm00471276_m1), Col1a2 (Mm00483888_m1), Col6a1 (Mm00487160_m1), and Thbs4 (Mm00449057_m1). Samples were normalized to the control (monoculture) group, and all data were analyzed using the 2^−ΔΔCt^ method.^105^

### Tissue Culture Immunofluorescence

Cells were permeabilized with 0.2% Triton X-100 (Sigma) in DPBS for 10 min at room temperature. After a 5 min DPBS wash, cells were blocked with 10% BSA with 0.05% Tween 20 in DPBS for 30 minutes. Primary-conjugated anti-Collagen 1 antibody (AF647, ab309367) was incubated overnight at 4°C. Cells were then washed once in blocking buffer before a 1-hour incubation with Phalloidin (AF488, A12379). Subsequently, cells were washed three times with DPBS, counterstained with 10x Spectral DAPI (Akoya, FP1490), washed again with DPBS, and then stored in DPBS at 4°C. Imaging was performed on an Axio Imager.A2 (Carl Zeiss Microscopy, LLC) and Zen software (Zeiss). Images were linearly contrasted for clarity using primary delete controls to set the contrast levels.

### Bulk RNA sequencing sample collection

Whole tissues were isolated and immediately placed in RNAlater (Sigma) at 4°C for at least 24 hours and at maximum 1-week before being transferred to Trizol (ThermoFisher Scientific) and –80°C storage. Tissues were homogenized with ceramic beads (2.8 mm; OMNI international) on a Bead Ruptor 12 (OMNI international) at the highest speed for 15 seconds 2-3 times; between rounds samples were kept on ice. Chloroform (Sigma) extraction was performed followed by RNA isolation and purification using the RNeasy Plus Mini Kit (Qiagen) following manufacturer instructions.

### Bulk RNA sequencing analysis

FASTQ files were aligned to the mouse genome (GRCm38/mm10) and GENCODE vM23 transcriptome annotation using STAR aligner (v2.7.84).^106–108^ Gene expression quantification was performed concurrently with alignment using STAR’s built-in gene counting feature. Differential gene expression analysis was conducted with DESeq2, incorporating a design matrix that accounted for mouse strain.^109^ Statistical significance for differentially expressed genes was determined using a false discovery rate (FDR) – adjusted p-value threshold of 0.05. For plotting, gene models, ribosomal genes, and immunoglobulins were excluded. Ranked gene set enrichment analysis (GSEA) was performed with fgsea (v1.25.1), ranking genes by the product of the log2 fold change and the negative log10 of the adjusted p-value, using the Hallmark gene sets.^110,111^ Pathways with an FDR-adjusted p-value below 0.05 were deemed significantly enriched. Matrisome-associated genes were identified by selecting those present within the matrisome gene sets.^112^ Genes associated with vascular development were selected from the gene ontology biological process gene sets containing “angiogenesis,” “venous_blood,” “vasodilation,” “vascular_wound_healing,” or “blood_vessel” in the name. All gene sets were sourced from the Molecular Signatures Database.

## scRNAseq and scTCRseq analysis

### Data preprocessing

scRNAseq and scRNAseq data were obtained, quality controlled, filtered, and preprocessed as described previously in Ruta et al. and are available at GEO accession number GSE306147.^33^ Unless stated otherwise, differential expression analysis compared each cluster against all other cells. Within the Treg subclustering, 3 small clusters of doublets were removed and 2 clusters that differed only by genes associated with processing artifacts were combined for downstream analysis.

### T cell receptor analysis

TCR sequencing data were integrated with gene expression data using the scRepertoire package (version 2.0.0).^113^ Cells with sequences for both αβ and γδ TCRs were regarded as doublets and removed from further analysis. Clonotypes were defined by the presence of identical gene and nucleotide sequences in both chains for αβ T cell analyses. In cases where gene, nucleotide, or chain information was incomplete, TCRs were grouped only with others sharing the same available sequence and pattern of missing data to prevent inflation of clonal numbers or ambiguous assignment of clones. To categorize clonal expansion levels in αβ T cells, cell counts were stratified based on the 75th, 90th, 99th, and 99.99th percentiles of clone sizes. These thresholds were adjusted within scRepertoire for optimal fit to the observed data distribution.^113^ Clones were determined using the αβ TCR repertoire of every individual animal within the CD3 dataset. Subsampled repertoire analysis involved randomly selecting 150 cells (CD3 dataset) or 50 cells (Treg dataset) from each sample’s repertoire. Diversity coefficients were then calculated for each downsampled repertoire 10,000 times utilizing DescTools (version 0.99.54) or vegan (v2.6.4).^114,115^ Subsequently the mean and standard deviation for every sample was determined and the mean and standard error of the mean for each time point was calculated. Two samples from the 1-week time point, and an additional 1-week sample from the Treg analysis, possessed insufficient cell counts for robust analysis and were thus excluded. Clustering of shared clones was performed by identifying all clones shared across at least 2 Treg clusters and quantifying the number of cells within each cluster with a given shared clone. Hierarchical clustering with complete linkages was performed using the incidence of each clone within each cluster to group the Treg populations and clones by similar incidence profile.

### Trajectory Inference

Pseudotime inference was performed using slingshot v2.18.0 using the corrected principal components returned by Harmony after batch effect correction.^116^ The LN Tregs were selected as the starting cluster due to their undifferentiated phenotype, and Treg 1 was identified as the end cluster due to its clonality and effector state. As implemented in the tradeSeq v3.2.2 package, a general additive model was used to model relationships between gene expression and pseudotime.^117^ A test was performed to assess the association of gene expression to pseudotime, and p-values were adjusted using the FDR method.

### Communication analysis

The cell communication pipeline previously described in Ruta et al. was implemented for this data.^33^ Briefly, intercellular signaling pathway analysis employed CellPhoneDB v4.1.0.^44^ Gene orthologs between human and mouse were determined using Ensembl 111, GRCh38.p14.^118^ Genes denoted as secreted and as ligands which were expressed in at least 5% of cells in a Treg cluster were identified and visualized.

TF regulons and activation were inferred using pySCENIC v0.12.1. These regulons and TF activity scores, along with CellPhoneDB v4 ligand-receptor database, were provided as input to dominoSignal v0.99 to predict communication between clusters.^43,44,53^ Analyses were performed independently for each 6-week timepoint replicate, then subsetted to include only those pathways consistently identified across all samples within each analysis type (CD45-enriched or CD3+ T cell). The CD45-enriched data and the T cell data were connected using ligand-receptor pairings from CellPhoneDB. These networks were then filtered to include only the receptor-TF correlations within a cluster greater than 0.25 with a p-value below 0.05. For Treg interactions with fibroblasts and endothelial cells, the thresholds for ligand and receptor expression were set to 0.05, and a threshold of 0.01 was implemented for TF activation.

Interaction strength was quantified using a composite communication score. Ligand expression, receptor expression, and TF activation were each scaled to a range of 1 to 10 using v1.3.0 of the scales package.^119^ These scaled values were multiplied, thereby equally weighting each pathway component. For clarity, the resulting scores were adjusted by order of magnitude as the units are arbitrary.

## Data visualization

Schematics used in figures were created with Biorender.com. Volcano plots were created using EnhancedVolcano v1.16.0 and chord diagrams were constructed using circlize v0.4.16 and refined in Affinity Designer.^120,121^ Heatmaps were created using ComplexHeatmap v2.14.0 or ggplot2 from the tidyverse v2.0.0.^122–124^ Network communication plots were produced with igraph.^125,126^ Supplementary packages included ggnewscale v0.4.10, ggrepel v 0.9.5, and svglite v2.1.3 for visualizations, and stringdist v0.9.12, reshape2 v1.4.4 and plyr v1.8.9 for data manipulation.^127–132^

## Data Availability

All sequencing data generated for this analysis are available for download from NCBI GEO, at accession numbers GSE306145, GSE306147, and GSE306253 (Foxp3-DTR bulk RNA sequencing, CD3 scRNA/TCRseq, and CD45-enriched scRNAseq, respectively). The scRNAseq/TCRseq and scRNAseq datasets at GSE306147 and GSE306253 are also used in a separate manuscript (see Ruta et al), with this study focusing on Tregs and their relation to stromal cells.^33^

## Code Availability

Communication inference analysis was performed using a pre-release branch of dominoSignal to implement experimental features not yet included in the package (branch krishnan_analysis, commit: de6b93f).^133^ This version can be accessed in the dominoSignal Github repository at https://github.com/FertigLab/dominoSignal. Custom scripts used in this analysis were written in R v4.2.1.^134^ Code for analyses will be made publicly accessible in the Elisseeff lab’s Github repository at www.github.com/Elisseeff-Lab/regulatory_t_cells_in_fibrosis upon publication.

## Author Contributions

JCM, KK, and JHE conceptualized and designed the studies. JCM, AR, ASR, EGG, LDH, DRM, ANR, PA, and MB developed and performed experimental work. KK and FHY performed bioinformatics analyses. KK and SN developed software. ANR annotated data. KK, JCM, ASR, and JHE interpreted results. EJF supervised software development. DMP and JHE supervised the study. KK, JCM, and JHE wrote the manuscript. All authors reviewed and approved the final manuscript.

## Acknowledgements

We would like to thank the Sidney Kimmel Cancer Center Flow Cytometry Technology Development Center for performing cell sorting (with funding from NCI CCSG P30 CA006973), C M Cherry Consulting for processing the CD45-enriched scRNAseq data, the Johns Hopkins University Transcriptomics and Deep Sequencing core facility for performing bulk and scRNAseq, the JHU Oncology Tissue Services SKCCC core facility for histology, and Dr. Katlin Stiver for flow cytometry. This study was funded by NIH Pioneer Award DP1AR076959 (JHE), NIH NIA K99/R00-AG081564 (JCM), and NSF GRFP DGE1746891 (AR).

