## Supplemental Figures and Tables for "Regulatory T cells clonally expand and contribute to stromal cell function in fibrotic response to synthetic implants"


Supplemental Figure 1. T cell checkpoint flow cytometry panel and quantification of populations 1
Supplemental Figure 2. Identification of Tregs in muscle vs inguinal lymph node by flow cytometry 2
Supplemental Figure 3. Immunofluorescence staining controls and single channel images for Figure 1F inset 3
Supplemental Figure 4. Gating strategy and quantification in Foxp3 DTR vs Wildtype mice using GFP Checkpoint Panel 4
Supplemental Figure 5. Gating strategy and quantification in Foxp3 DTR vs Wildtype mice using GFP Pan-Immune Panel 5
Supplemental Figure 6. Mean and standard deviation of cluster proportions across time points for T cell scRNAseq. 6
Supplemental Figure 7. Single-cell RNAseq analysis of Treg subsets 7
Supplemental Figure 8. Detection of CD25 low Tregs in scRNAseq and flow cytometry 8
Supplemental Figure 9. Differential gene expression identifies genes distinguishing effector Treg subpopulations. 9
Supplemental Figure 10. Treg populations all express high levels of activation and tissue residency markers with temporal shifts to population proportion and effector gene expression 10
Supplemental Figure 11. Single-cell TCR sequencing coverage and expansion analysis for sorted CD3+ T cells 11
Supplemental Figure 12. Downsampling analysis of TCR repertoire diversity in scTCRseq data 12
Supplemental Figure 13. Quantification and visualization of clonal expansion within Treg subclusters 13
Supplemental Figure 14. TCR signaling and top clones in Treg subclusters. 14
Supplemental Figure 15. Clonal overlap among Treg subpopulations 15
Supplemental Figure 16. Trajectory inference identifies multiple pathways among Treg populations 16
Supplemental Figure 17. Association of pseudotime with Treg clonal expansion categories 17
Supplemental Figure 18. Secreted ligand expression profiles across Treg subclusters 18
Supplemental Figure 19. Comprehensive Treg-Fibroblast dominoSIgnal communication network 19
Supplemental Figure 20. Sox-family mediated Treg-fibroblast network and target gene analysis 20
Supplemental Figure 21. Imaging for phalloidin and collagen 1 after Treg-Fibroblast Coculture. 21
Supplemental Figure 22. Treg to endothelial cell communication network. 22
Supplemental Figure 23. Q4D in vivo Treg depletion timeline, weights, histology, and immune quantification 23
Supplemental Figure 24. Macrophage subset quantification following Q4D Treg depletion 24
Supplemental Figure 25. Imaging for proliferation and senescence markers following Q4D Treg depletion 25
Supplemental Figure 26. Imaging of vascular and stromal markers after Q4D Treg depletion 26
Supplemental Figure 27. Immunohistochemistry staining of neovascular marker CD105 after Q4D Treg depletion 27
Supplemental Table 1: T Cell Checkpoint Panel I
Supplemental Table 2: GFP T Cell Checkpoint Panel II
Supplemental Table 3: GFP Pan Immune Panel III
Supplemental Table 4: T Cell Sorting Panel IV
Supplemental Table 5: GFP Pan Stromal Immune Panel V
Supplemental Table 6: Immunofluorescence Staining Primary and Secondary VI
Supplemental Table 7: TotalSeq-C for Hashing VII
Supplemental Table 8: Custom Primers for γδ VDJ sequencing VIII
Supplemental Table 9: HTO Sequences for Demultiplexing Samples IX
Supplemental Table 10: Treg Canonical Markers Panel X

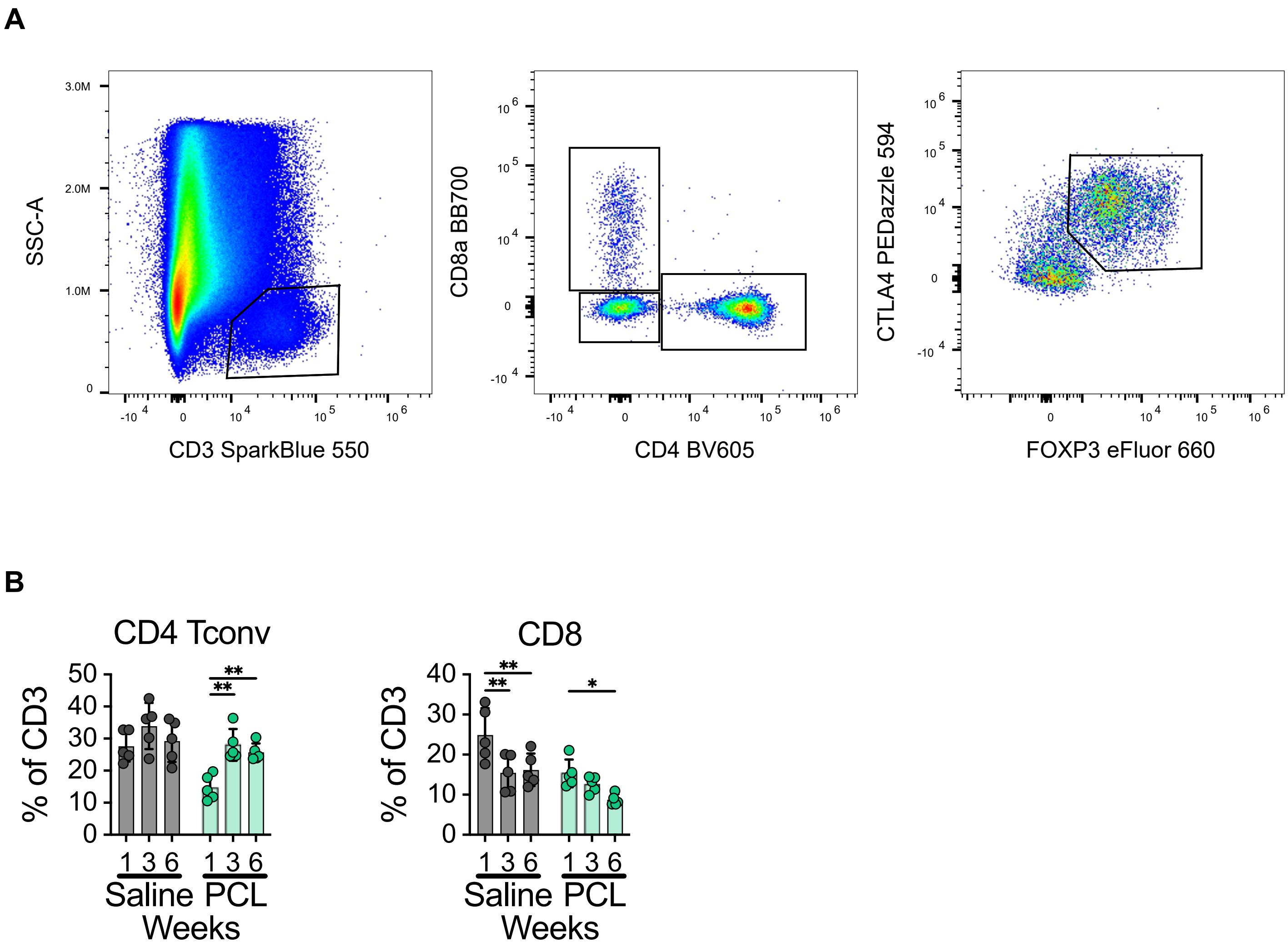


**Supplemental Figure 1. T cell checkpoint flow cytometry panel and quantification of populations**
A) Gating scheme for T cell immune checkpoint panel used in flow cytometry analysis. B) Quantification of the two T cell populations identified via gating strategy not included in main figure.

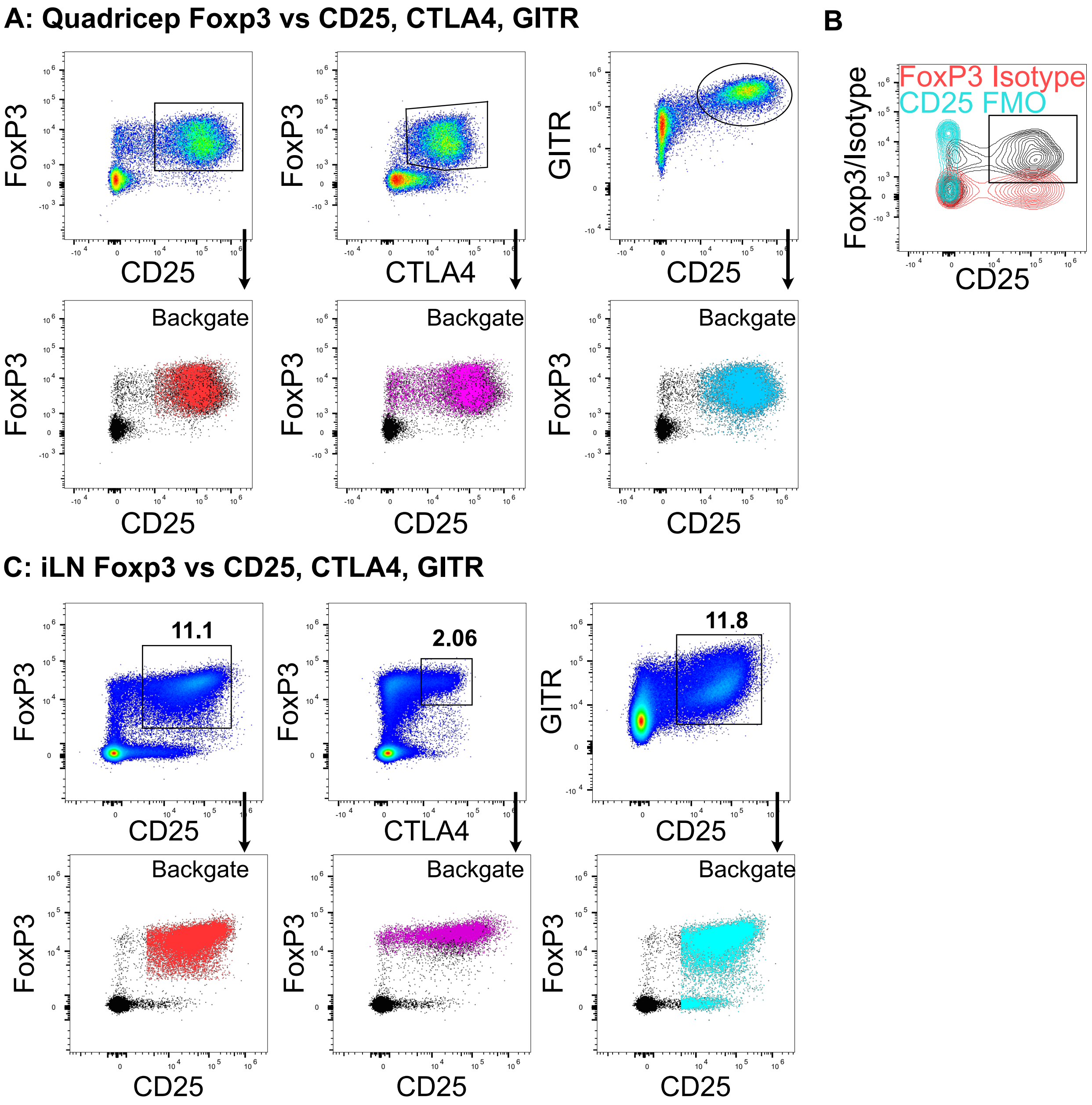


**Supplemental Figure 2. Identification of Tregs in muscle vs inguinal lymph node by flow cytometry**
A) Representative flow cytometry plots illustrating three strategies for Treg identification in quadriceps after CD4+ gating: double positive CD25+Foxp3+, CTLA4+Foxp3+, or CD25+GITR+. Top-row gates are backgated onto canonical Foxp3 CD25 gate in the second row. CTLA4 captures a subset of low-CD25 Tregs. See STable 10 for panel. B) Overlay of the Foxp3 Isotype and the CD25 FMO staining controls on a representative sample. C) Similar gating strategy in the inguinal lymph nodes shows  Foxp3+CD25+ population is not captured by CTLA4+ (only 2% relative to 11%) and GITR collects the Foxp3 negative CD25+ cells.

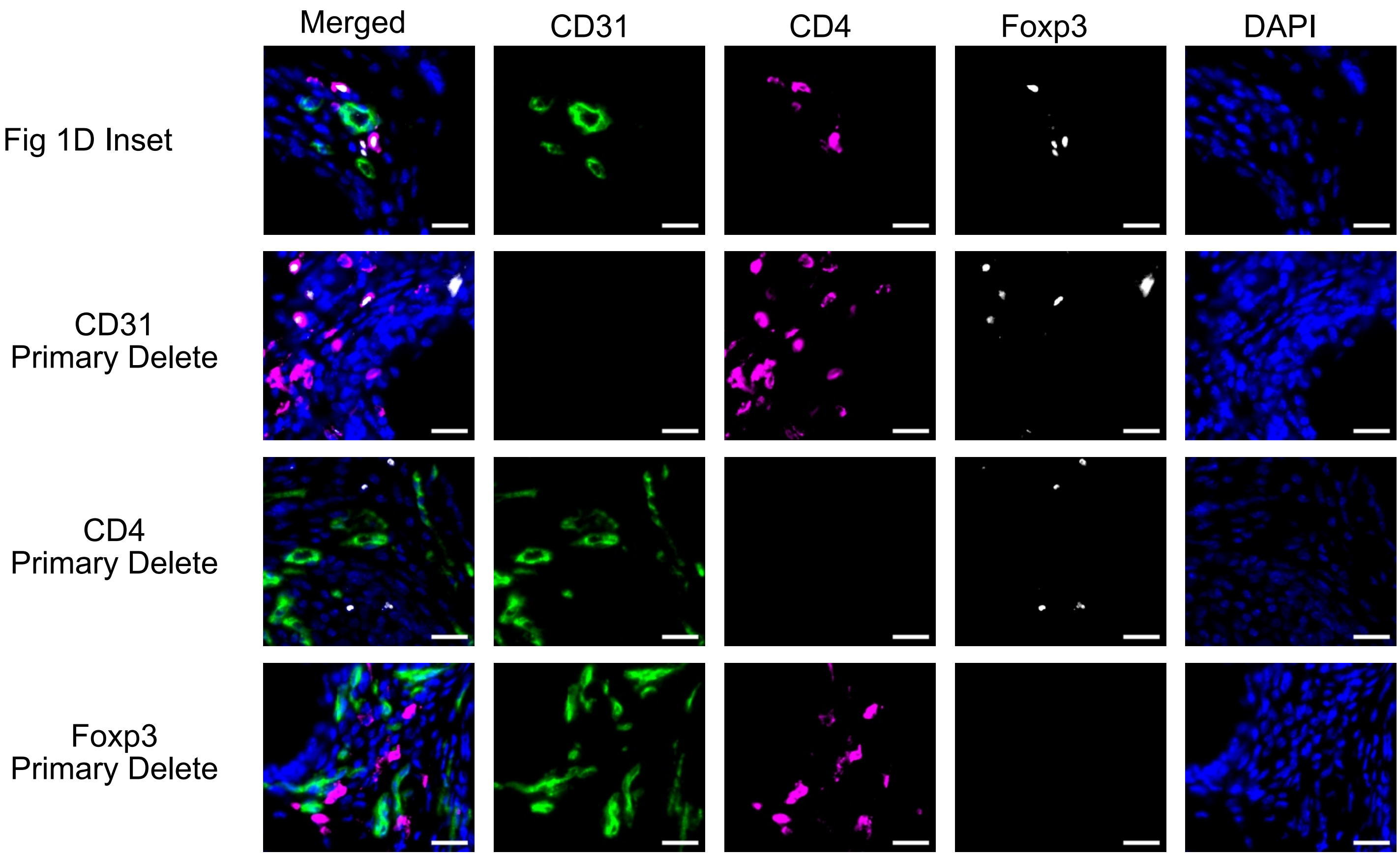


**Supplemental Figure 3. Immunofluorescence staining controls and single channel images for Figure 1F inset**
All images were acquired and linearly contrasted identically. Scale bars = 20μm.

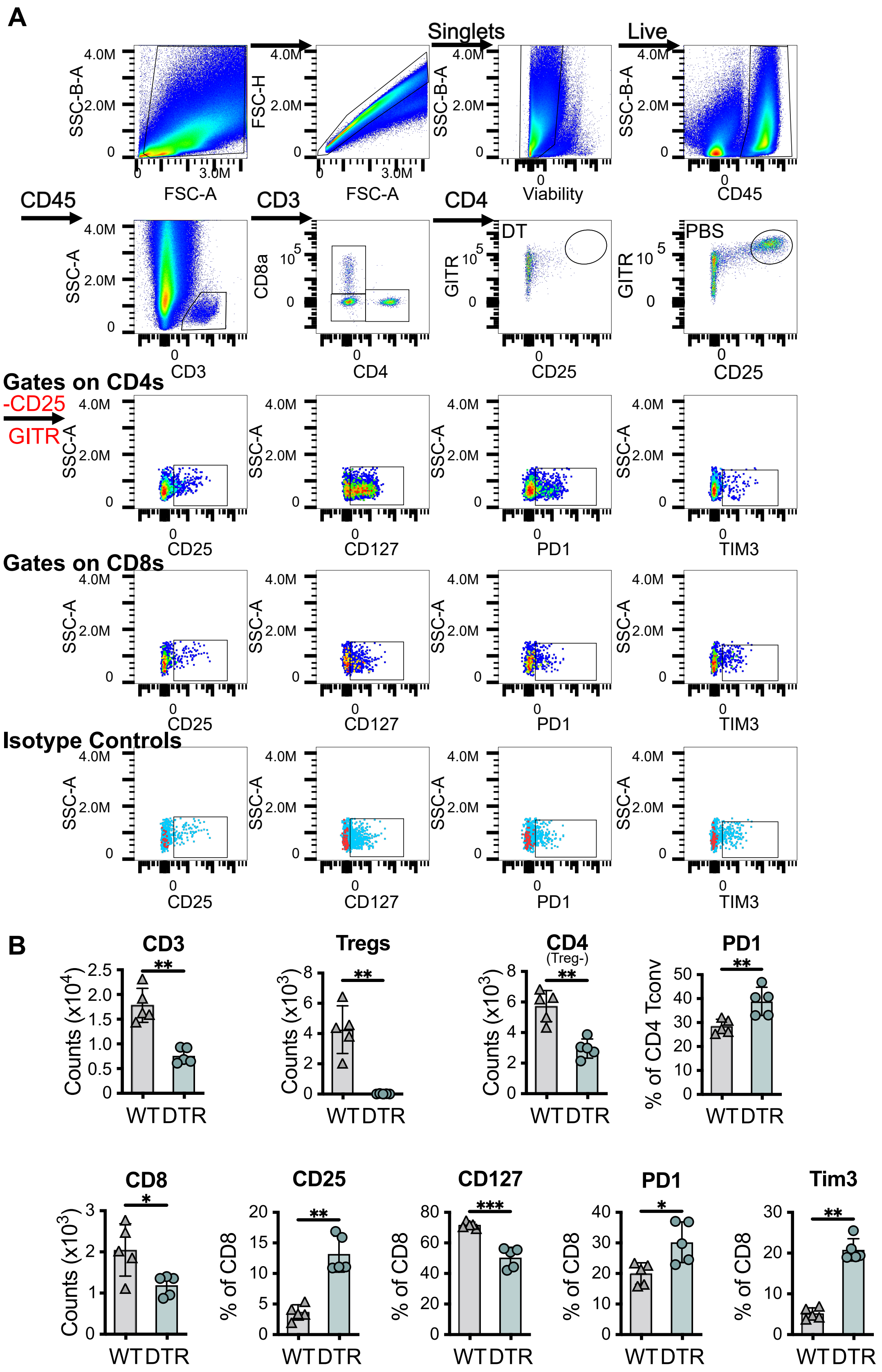


**Supplemental Figure 4. Gating strategy and quantification in Foxp3 DTR vs Wildtype mice using GFP Checkpoint Panel**
A) Gating strategy for cells collected from two quadriceps digested and processed with a percoll gradient; not gates indicated by red text over arrows. CD4+ and CD8+ T cells were gated from CD3. To remove the majority of Tregs from CD4+ analysis, Tregs were gated as CD25 GITR bright and a not gate was used to identify the conventional CD4+ T cells. CD25, CD127, PD1, and Tim3 were then quantified from CD4+ Tconv (no Tregs) or CD8+ T cells. B) Quantification of T cell subsets and T cell markers in WT and Foxp3 DTR mice with DT treatment 3 days before harvest. Statistical significance was assessed with unpaired two-sided t-test. $p < 0.05$ (*), $p < 0.01$ (**), $p < 0.001$ (***), $p < 0.0001$(****).

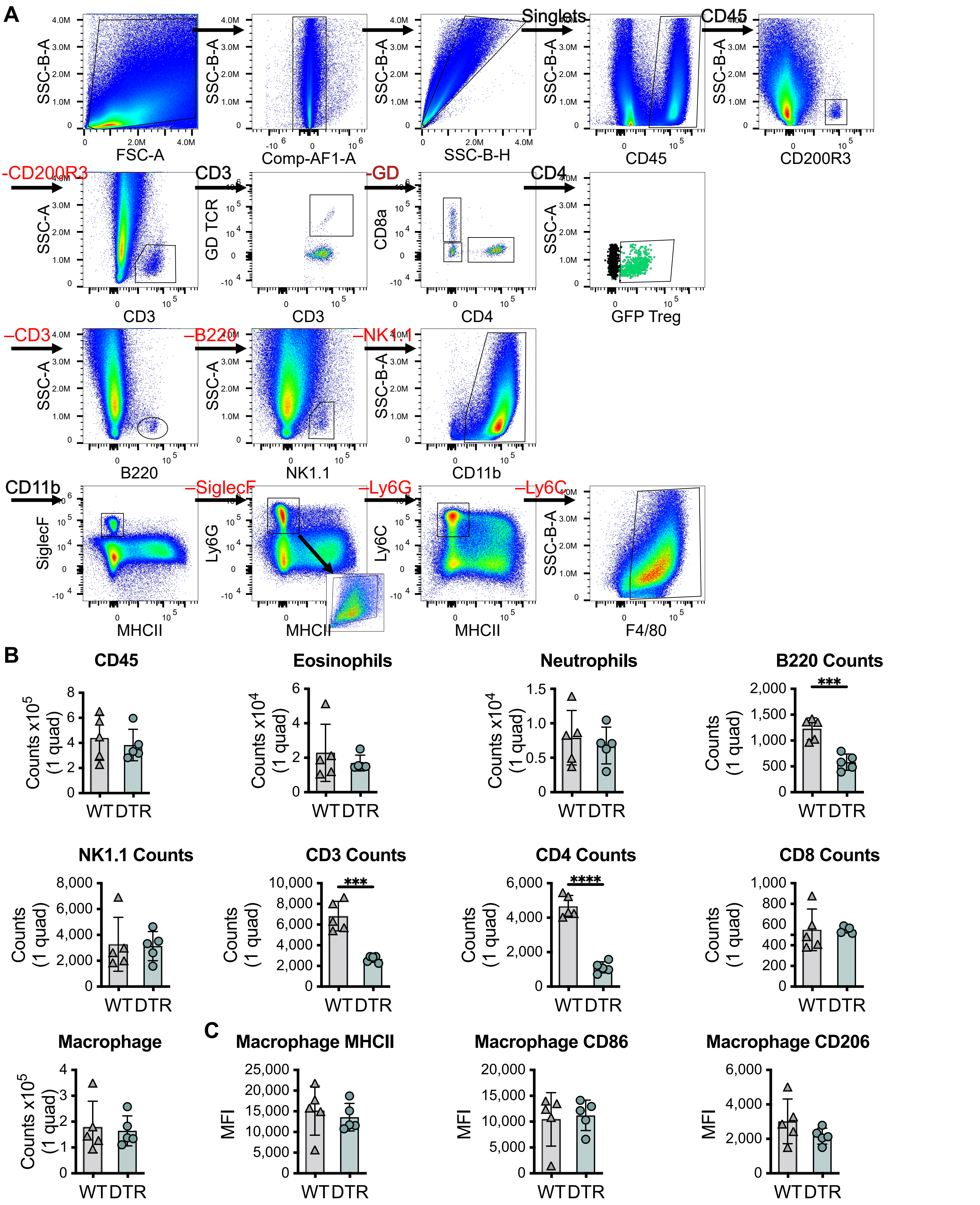


**Supplemental Figure 5. Gating strategy and quantification in Foxp3 DTR vs Wildtype mice using GFP Pan-Immune Panel**
A) Gating strategy for cells collected from a single quadricep digested without percoll gradient; not gates are indicated by red text over arrows. B) Comparison of immune population counts between WT and Foxp3 DTR mice for Treg depletion with DT treatment 3 and 1 days prior to harvest. C) Comparison of MFI in macrophage population between WT and Foxp3 DTR mice. Statistical significance was assessed with unpaired two-sided t-test: $p < 0.05$ (*), $p < 0.01$ (**), $p < 0.001$ (***), $p < 0.0001$ (****).

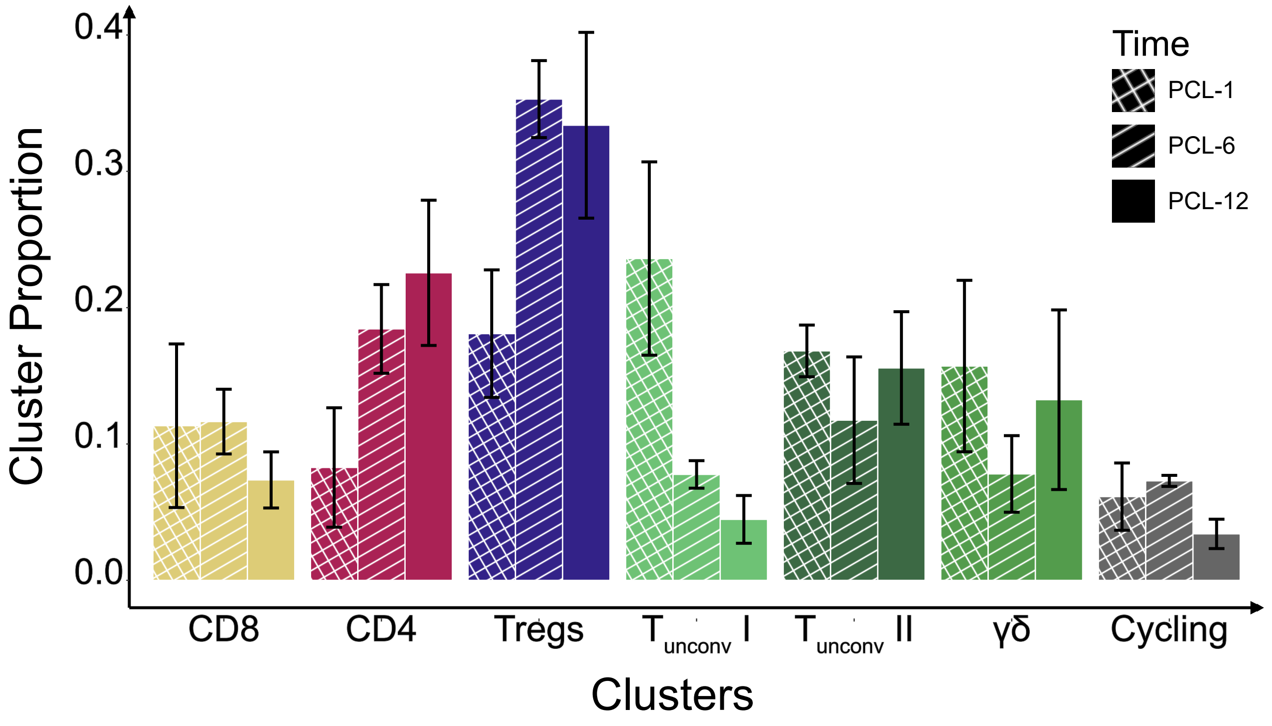


**Supplemental Figure 6. Mean and standard deviation of cluster proportions across time points for T cell scRNAseq.**

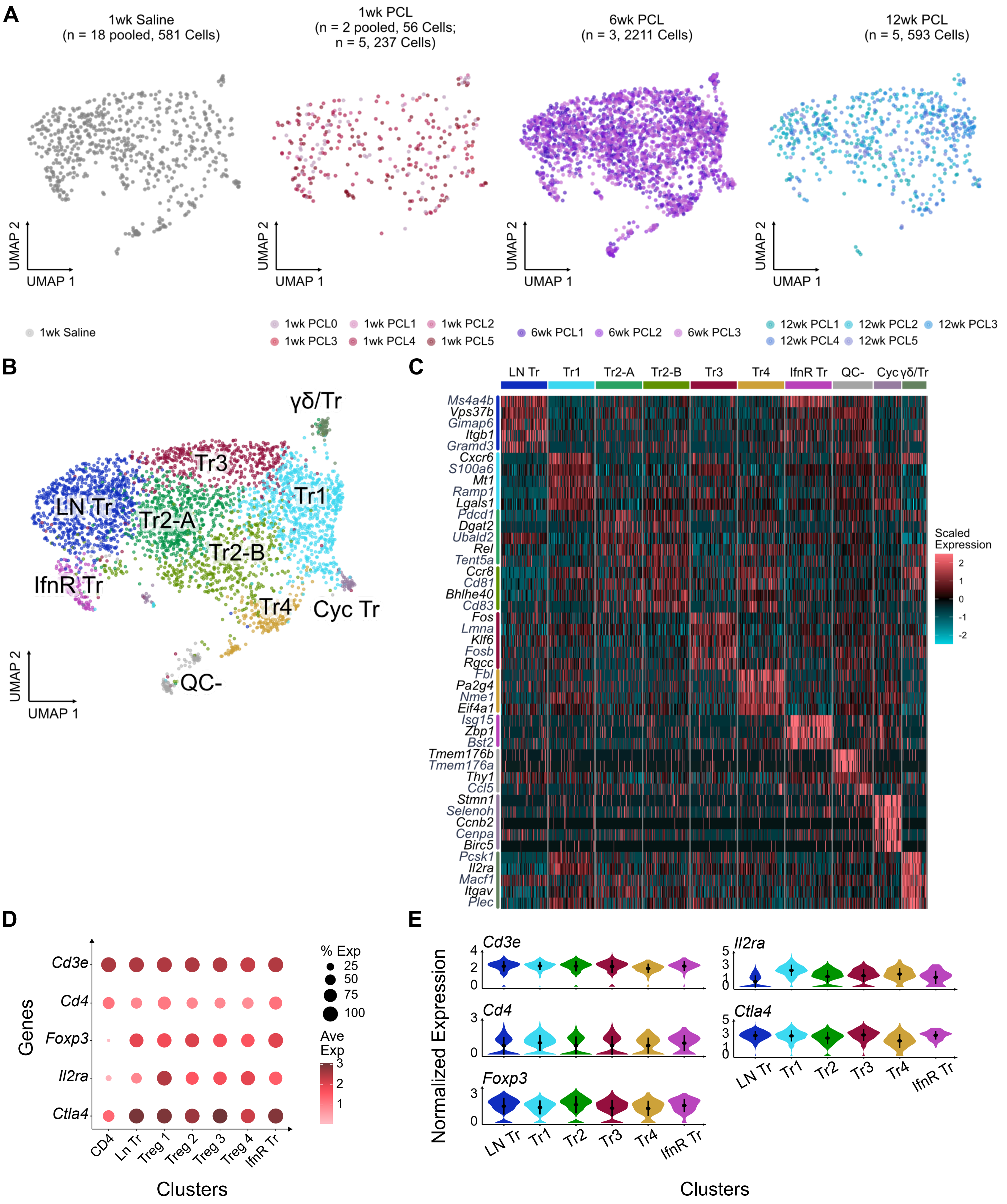

**Supplemental Figure 7. Single-cell RNAseq analysis of Treg subsets**
A) UMAP visualization of Tregs split by timepoint and material and colored by replicate. B) UMAP labeled with unsupervised clusters (subsequently merged and contaminants removed based on differential expression analysis). C) Heatmap showing differential gene expression for each unsupervised Treg cluster. D) Dot plot depicting average expression (color) and percent expression (size) of canonical Treg markers across CD4+ conventional T cells (reference) and Treg subpopulations. E) Violin plots showing distribution of normalized expression for canonical Treg marker genes among Treg subclusters. Point and line indicate mean ± standard deviation of expression.

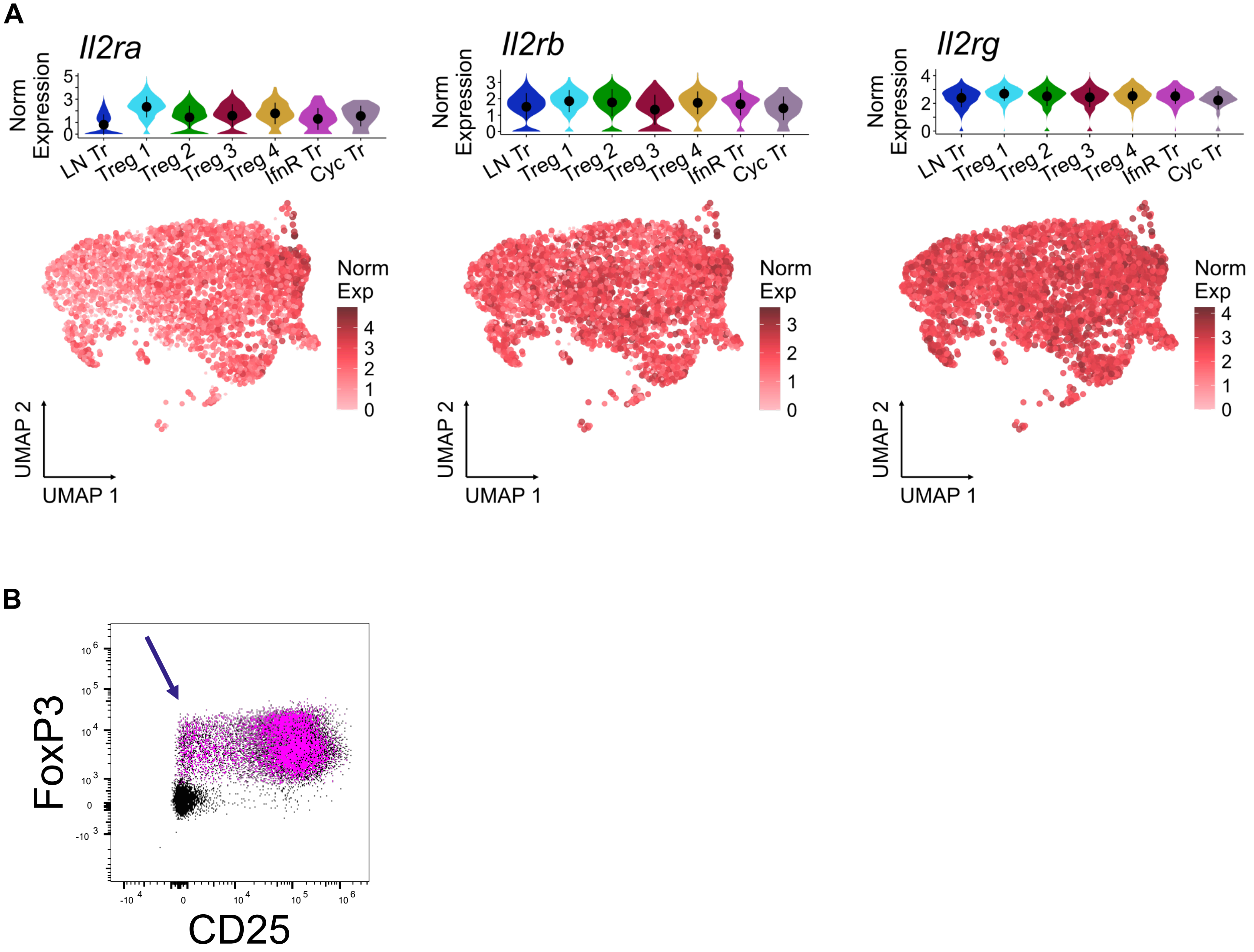


**Supplemental Figure 8. Detection of CD25 low Tregs in scRNAseq and flow cytometry**
A) Violin plots of normalized counts from Treg scRNAseq show LN-associated Tregs have low *Il2ra* expression but retain strong expression of other IL2 receptor subunits. B) Flow cytometry CD25 antibody PC-61 is specific to IL2Rα, and it identifies a population of Tregs with low CD25 surface expression (see STable 10 for panel).

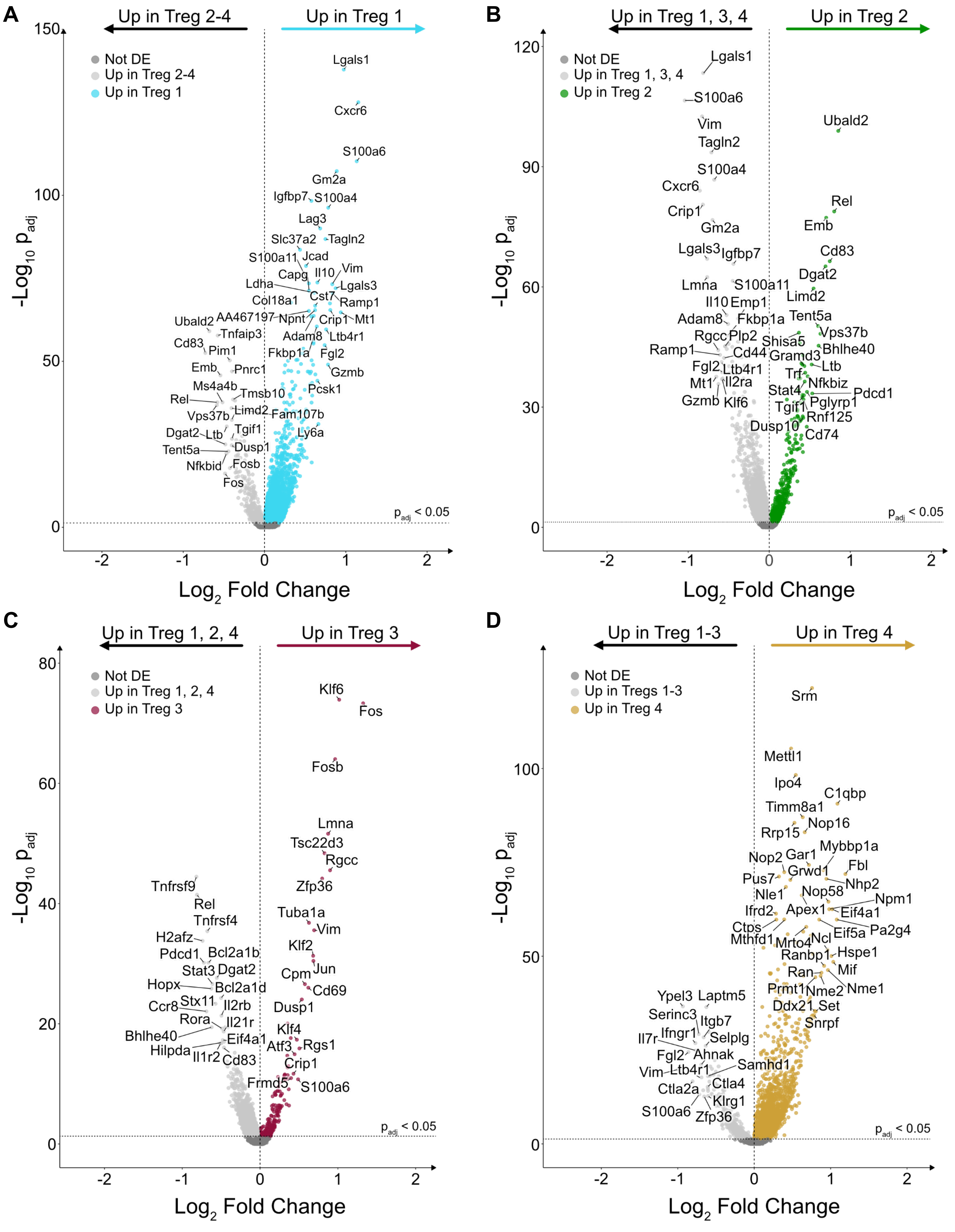


**Supplemental Figure 9. Differential gene expression identifies genes distinguishing effector Treg subpopulations.**
A-D) Volcano plots depicting differential gene expression profiles for each effector Treg subpopulation compared to the remainder of the effector Treg populations.

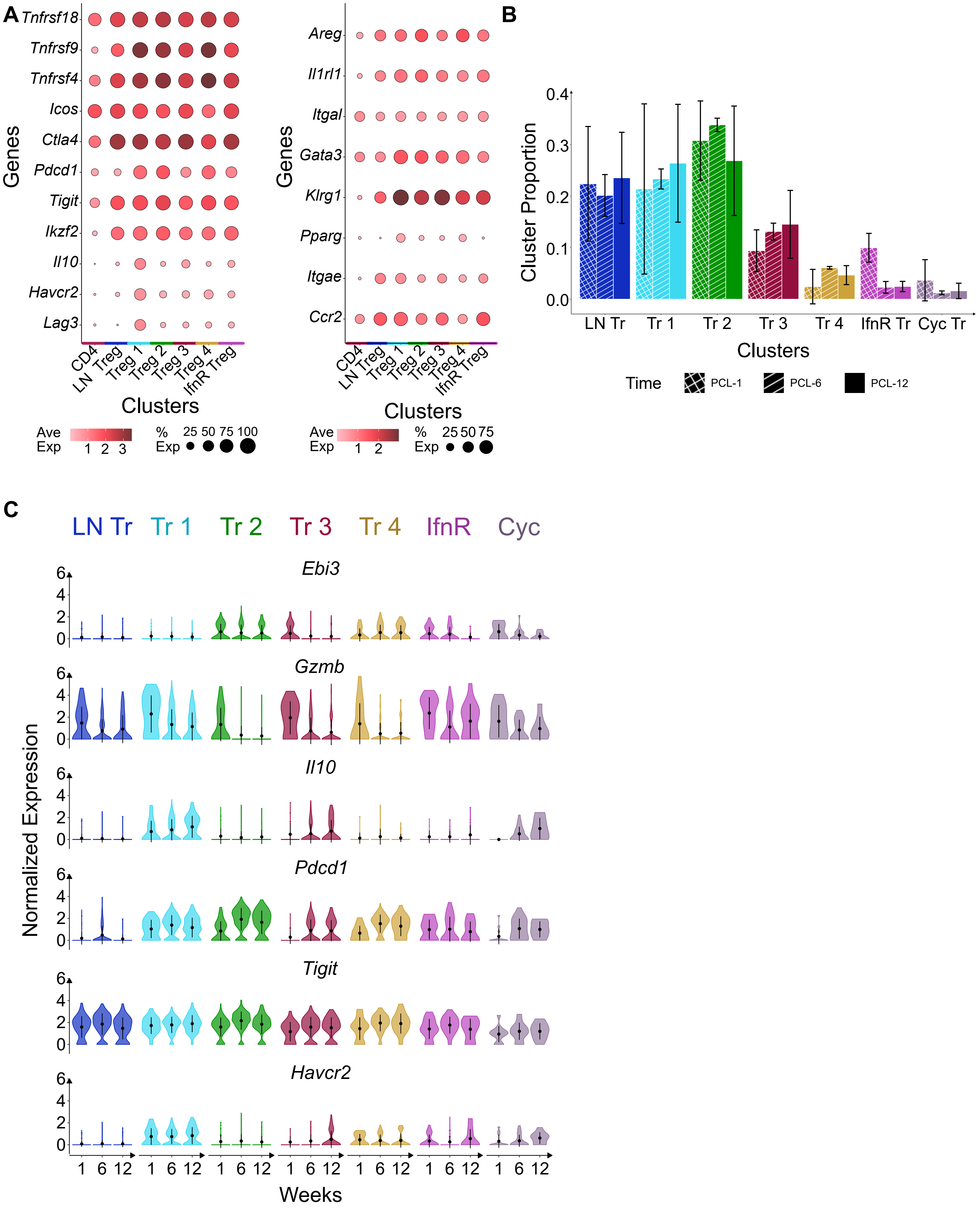


**Supplemental Figure 10. Treg populations all express high levels of activation and tissue residency markers with temporal shifts to population proportion and effector gene expression**
A) Average expression (color) and percent expression (size) of activation markers (left) and markers of tissue residence (right) show strong expression in all Treg populations relative to CD4+ conventional T cells. B) Mean and standard deviation of population proportions across replicates in each timepoint. C) Gene expression changes across all Treg subpopulations across timepoints.

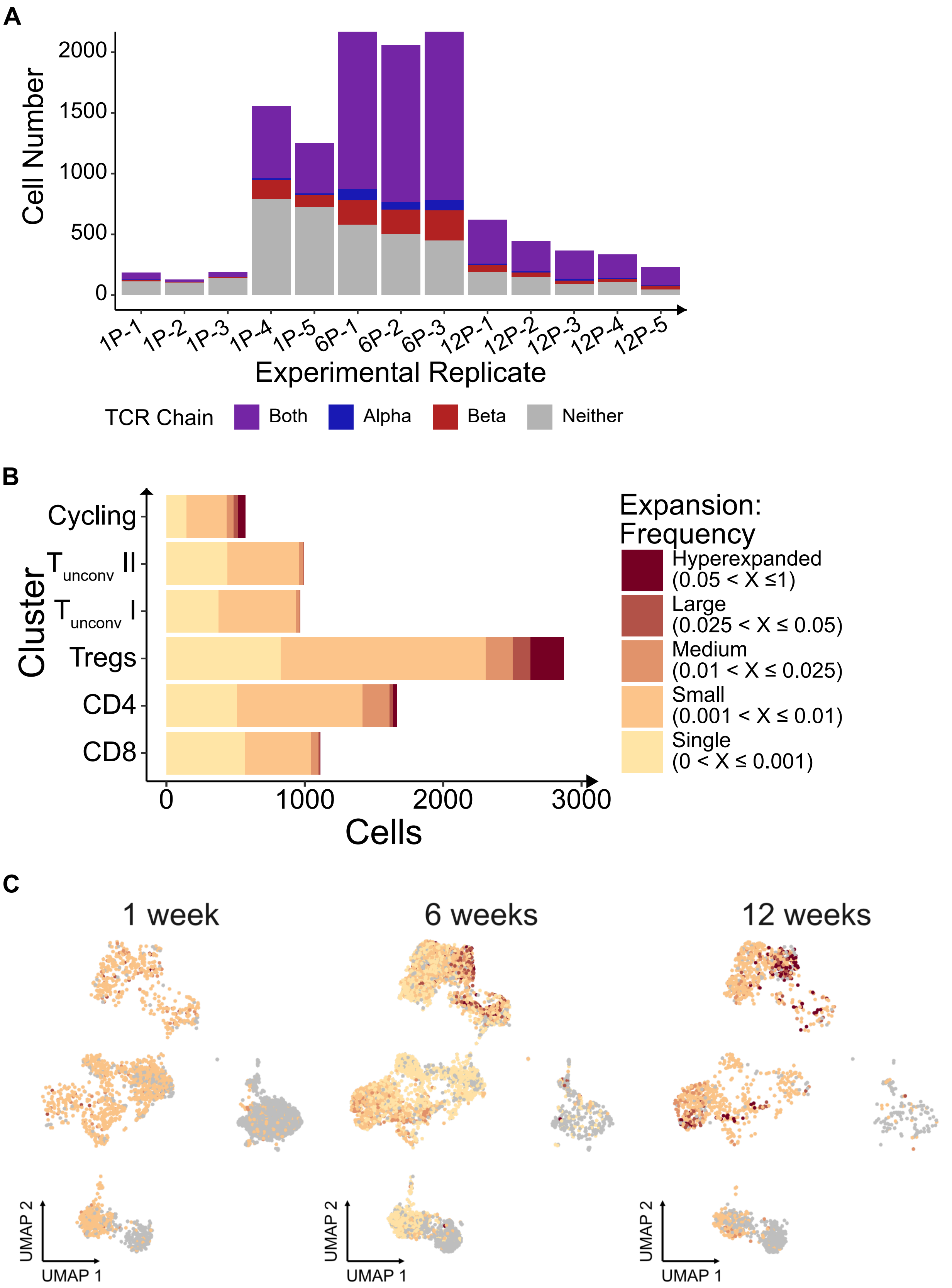


**Supplemental Figure 11. Single-cell TCR sequencing coverage and expansion analysis for sorted CD3+ T cells**
A) T cell receptor (TCR) α/β chain coverage for each experimental replicate. B) Cell counts by clonal expansion category (defined by clone frequency rather than number). C) UMAPs split by timepoint show cells colored by TCR expansion category (by frequency).

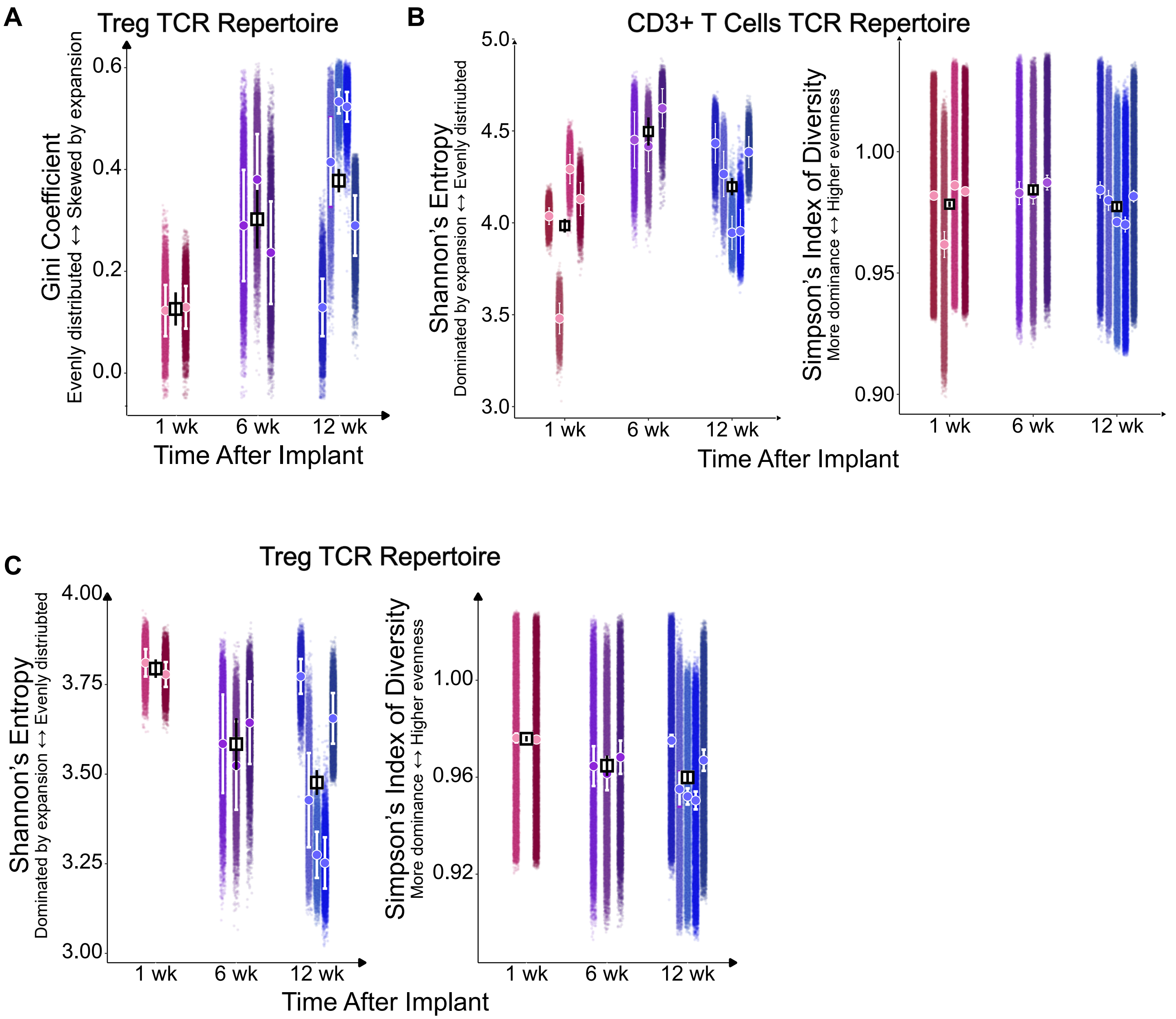


**Supplemental Figure 12. Downsampling analysis of TCR repertoire diversity in scTCRseq data**
A) Subsampling analysis of the Treg TCR repertoire with diversity quantified with Gini coefficient. B) CD3+ T cell TCR repertoire downsampling analysis assessed by Shannon's entropy and Simpson's diversity index (1-D). C) Treg TCR repertoire downsampling analyzed using Shannon's entropy and Simpson's diversity index. Repertoire downsampled 10,000 times; each point represents diversity coefficient value for a given downsampling. Circles and corresponding bars indicate mean ± sd for each replicate; square and bar indicate mean ± SEM for each timepoint.

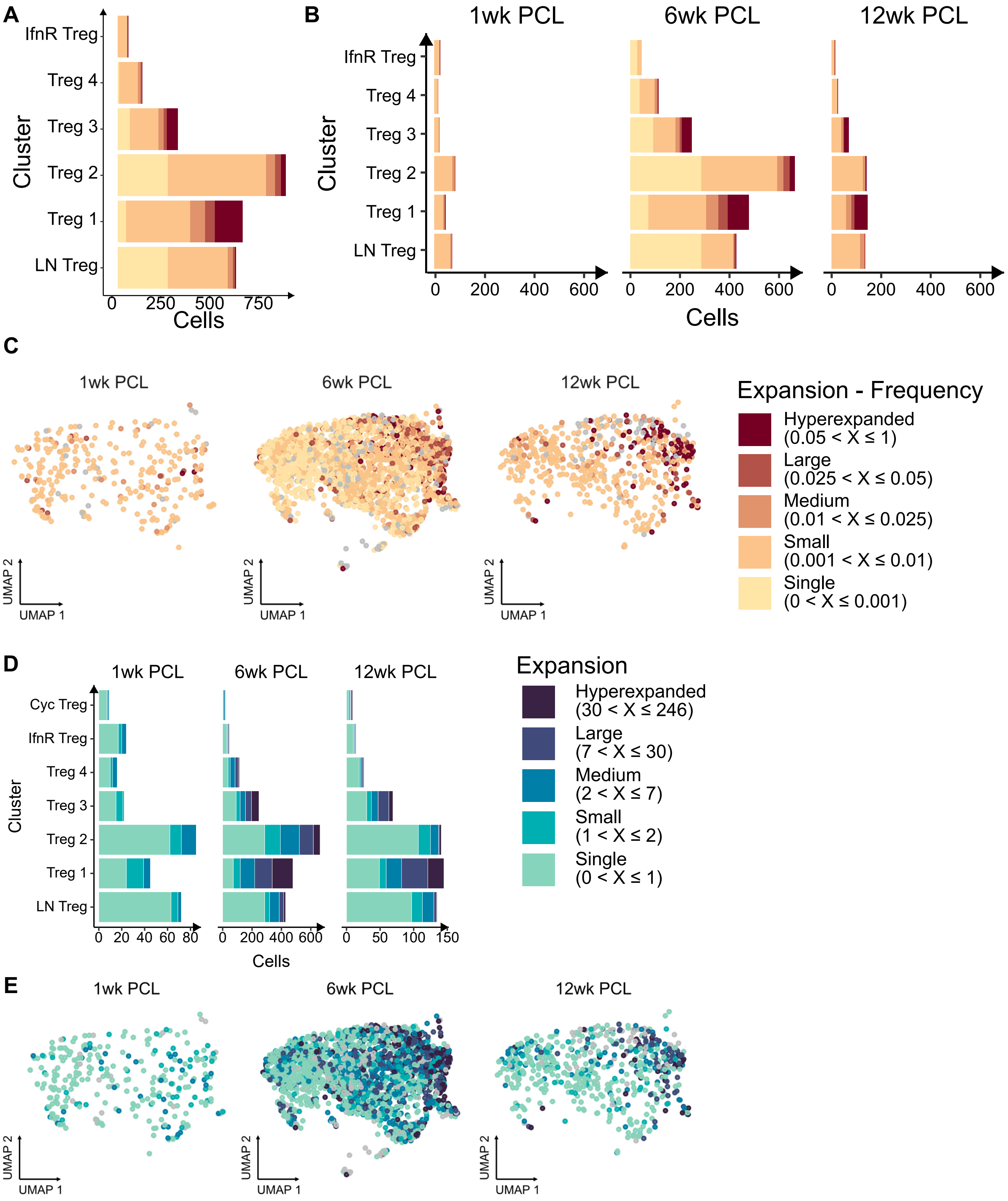


**Supplemental Figure 13. Quantification and visualization of clonal expansion within Treg subclusters**
A) Cell numbers for Treg subclusters by clonal expansion category (frequency-based). B) Cell counts for each expansion category split by time point. C) UMAPs, split by timepoint, colored by clonal expansion frequency category. D) Bar plots of cell number for each expansion category (by number) split by timepoint. E) UMAPs split by timepoint, colored by number-based expansion categories.

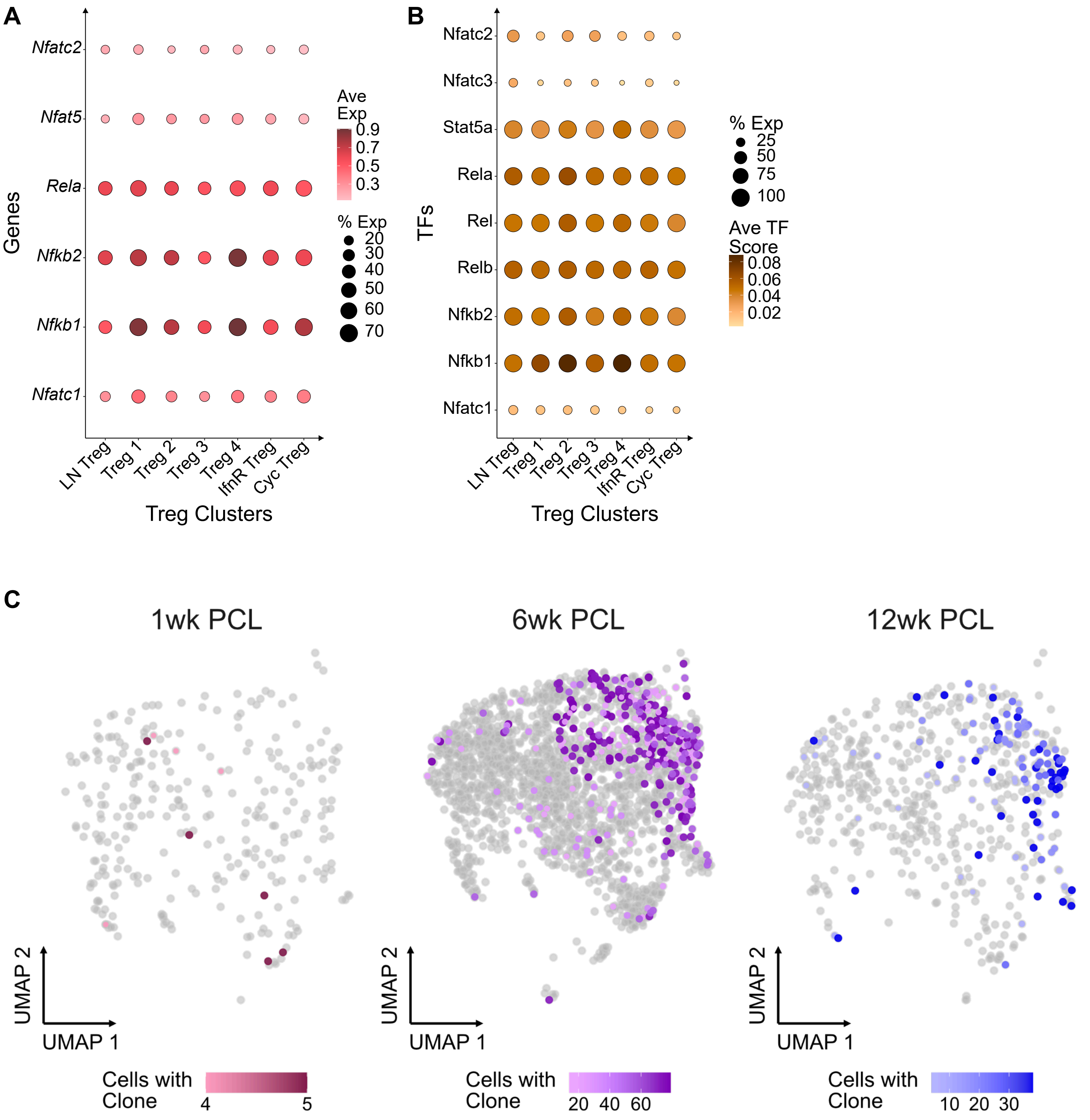


**Supplemental Figure 14. TCR signaling and top clones in Treg subclusters.**
A) Average (color) and percent of cells (size) for key TCR signaling gene expression and (B) TF activation scores shown across Treg subclusters. C) Top 5 clones (with ties when present) at each time point visualized on UMAP split by time point.

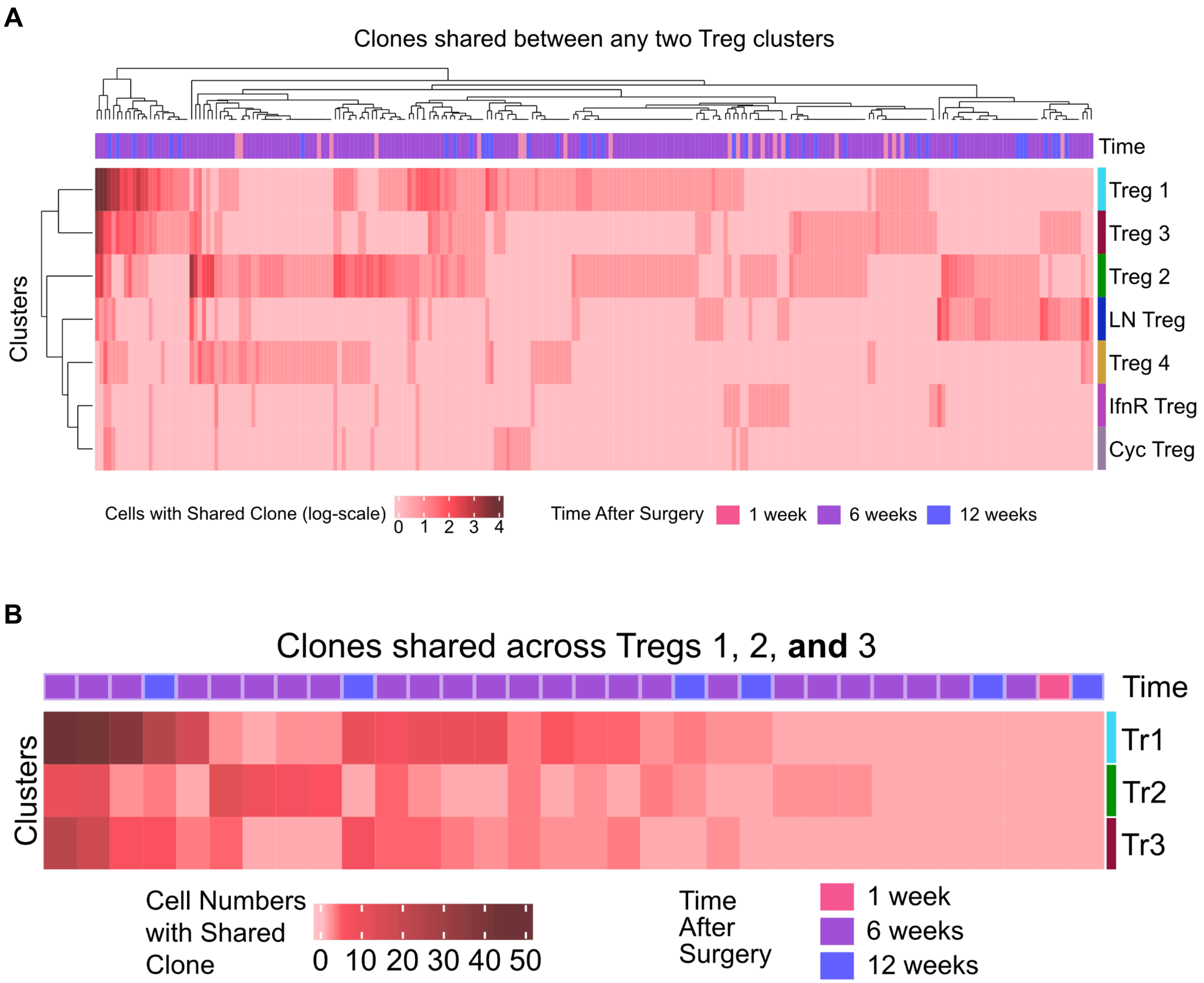


**Supplemental Figure 15. Clonal overlap among Treg subpopulations**
A) Heatmap displaying the incidence of clones shared between any 2 Treg clusters. Hierarchical clustering with complete linkages was performed using the incidence of each clone in each cluster. B) Heatmap showing clones shared between all 3 major effector Treg subclusters (Tregs 1, 2, and 3).

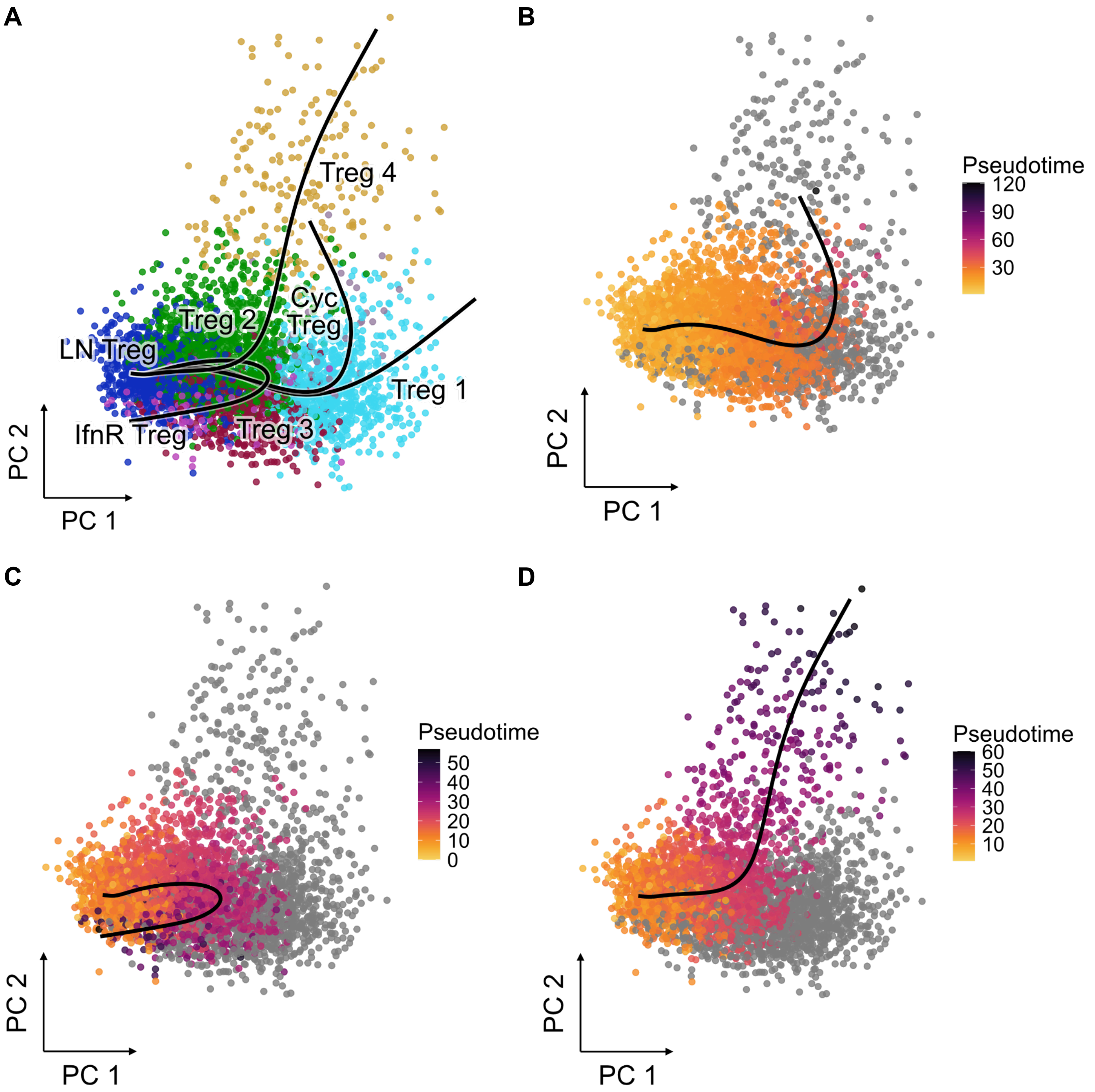


**Supplemental Figure 16. Trajectory inference identifies multiple pathways among Treg populations**
A) Principal component (PC) space visualization of all 4 inferred cell-state trajectories with cells colored by Treg cluster. B) Individual trajectories from LN associated Treg to Treg 2 to Treg 3 to Cycling Treg, C) LN associated Treg to Treg 2 to IFN Responsive Treg, D) LN associated Treg to Treg 2 to Treg 4 depicted on PC coordinates with cells colored by pseudotime values associated with each specific trajectory.

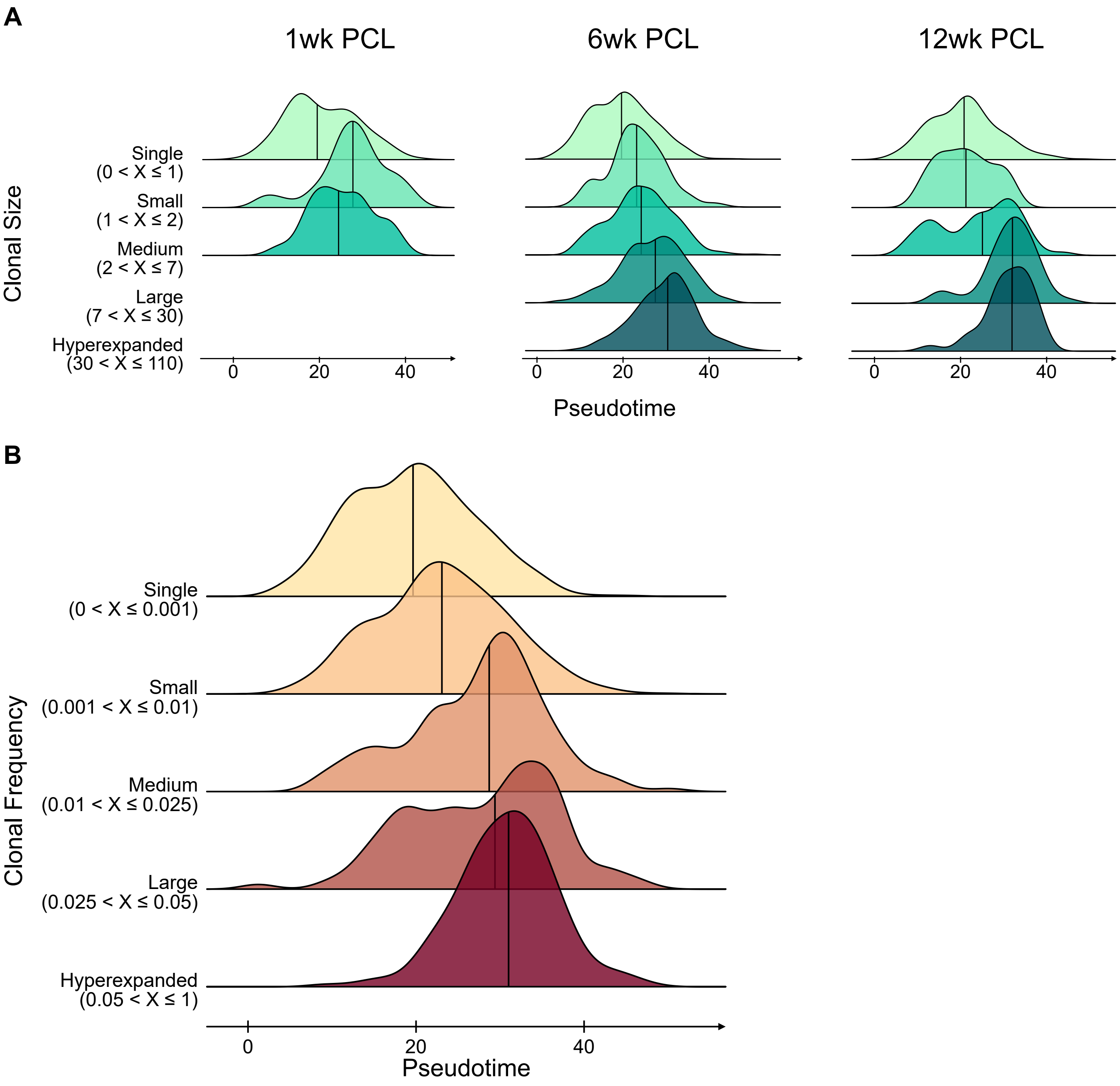


**Supplemental Figure 17. Association of pseudotime with Treg clonal expansion categories**
A) Distribution of pseudotime values for each clonal expansion category (defined by number), split by timepoint. B) Distribution of pseudotime values for each clonal expansion category (defined by frequency). Solid line within distribution indicates median value.

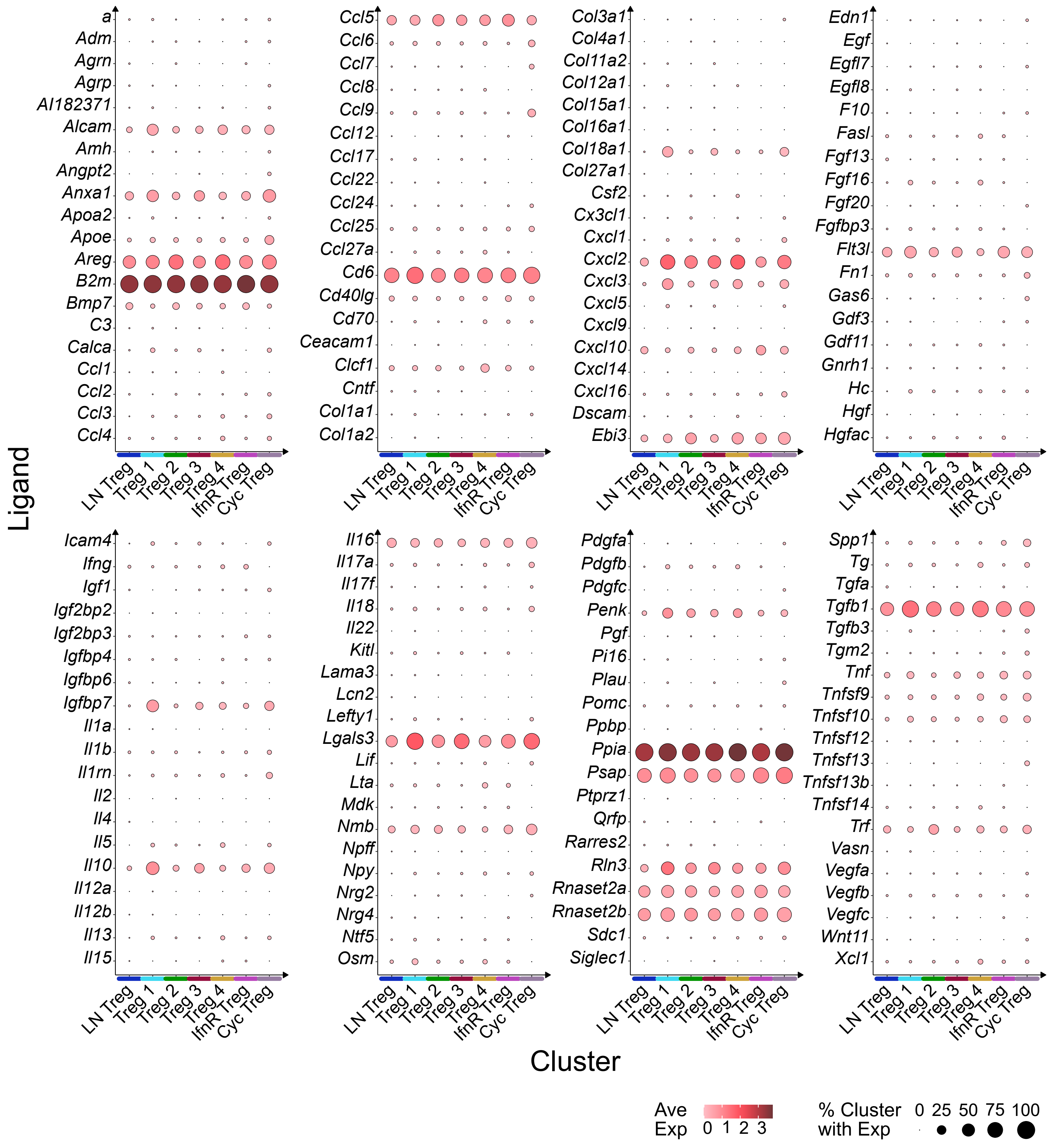


**Supplemental Figure 18. Secreted ligand expression profiles across Treg subclusters**
Dot plot displaying all secreted ligands expressed by Treg subclusters, where color indicates average normalized expression and dot size reflects percent of cells expressing the ligand within each subcluster.

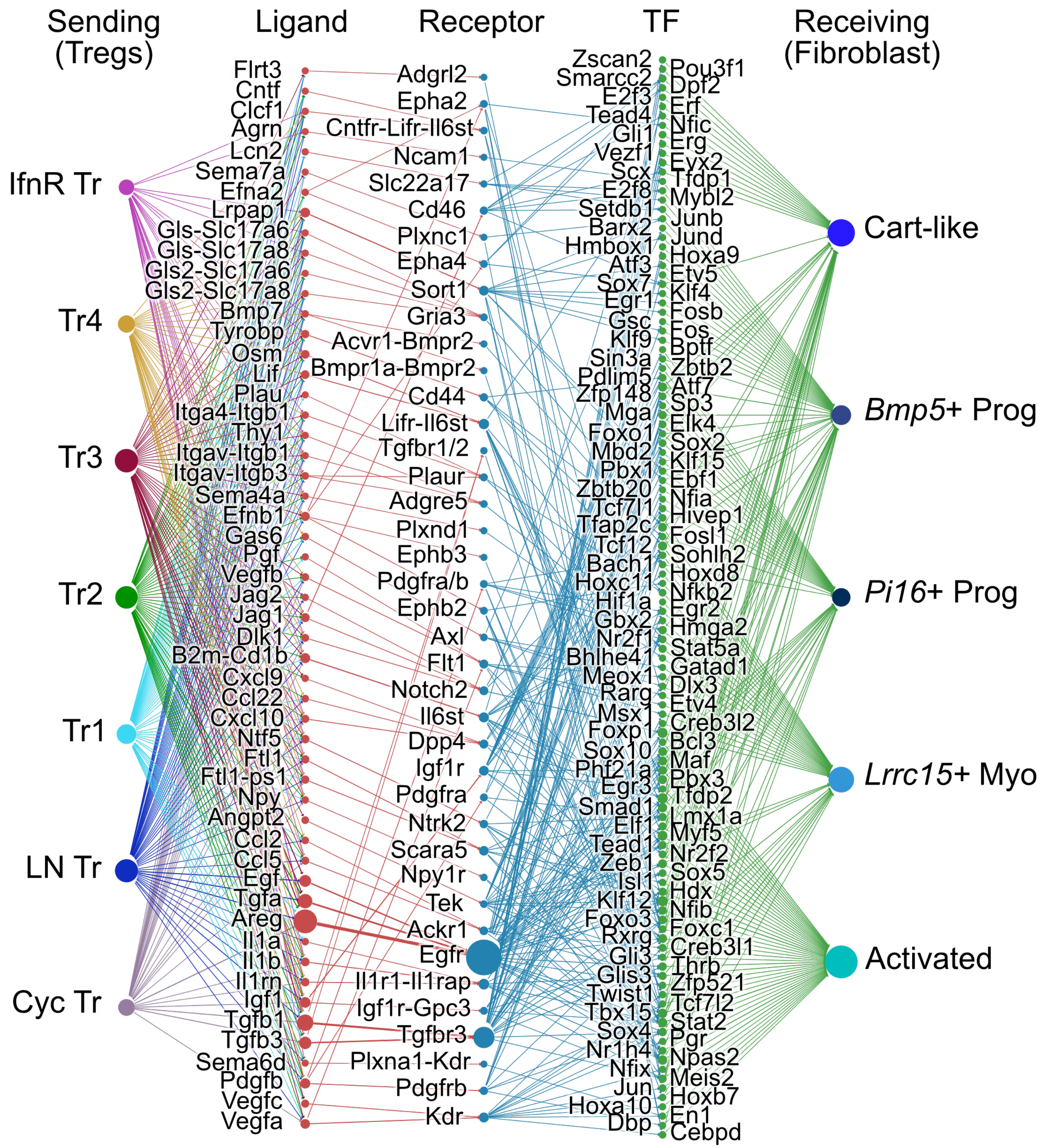


**Supplemental Figure 19. Comprehensive Treg-Fibroblast dominoSIgnal communication network**
Network diagram depicting all predicted signaling pathways linking Treg subpopulations to fibroblast clusters as identified in communication inference analysis. Node size and edge thickness correspond to number of pathways.

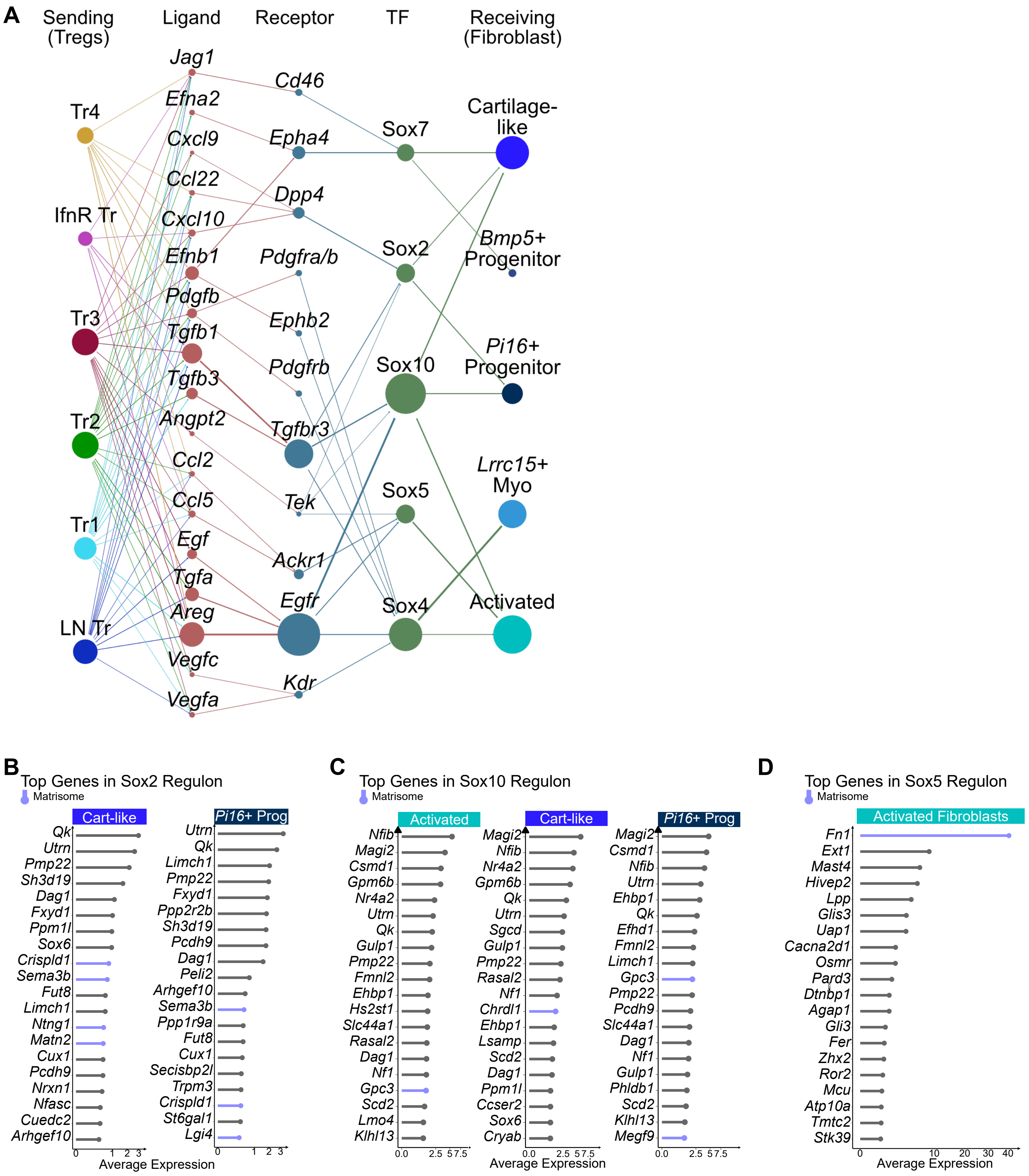


**Supplemental Figure 20. Sox-family mediated Treg-fibroblast network and target gene analysis**
A) Network of predicted Treg to fibroblast interactions mediated by Sox-family Tfs (all Treg clusters included). B) Top genes by average expression level in Sox2, C) Sox10, and D) Sox5 regulons shown for each fibroblast cluster with TF activation in Treg to fibroblast network; matrisome-associated genes annotated in blue.

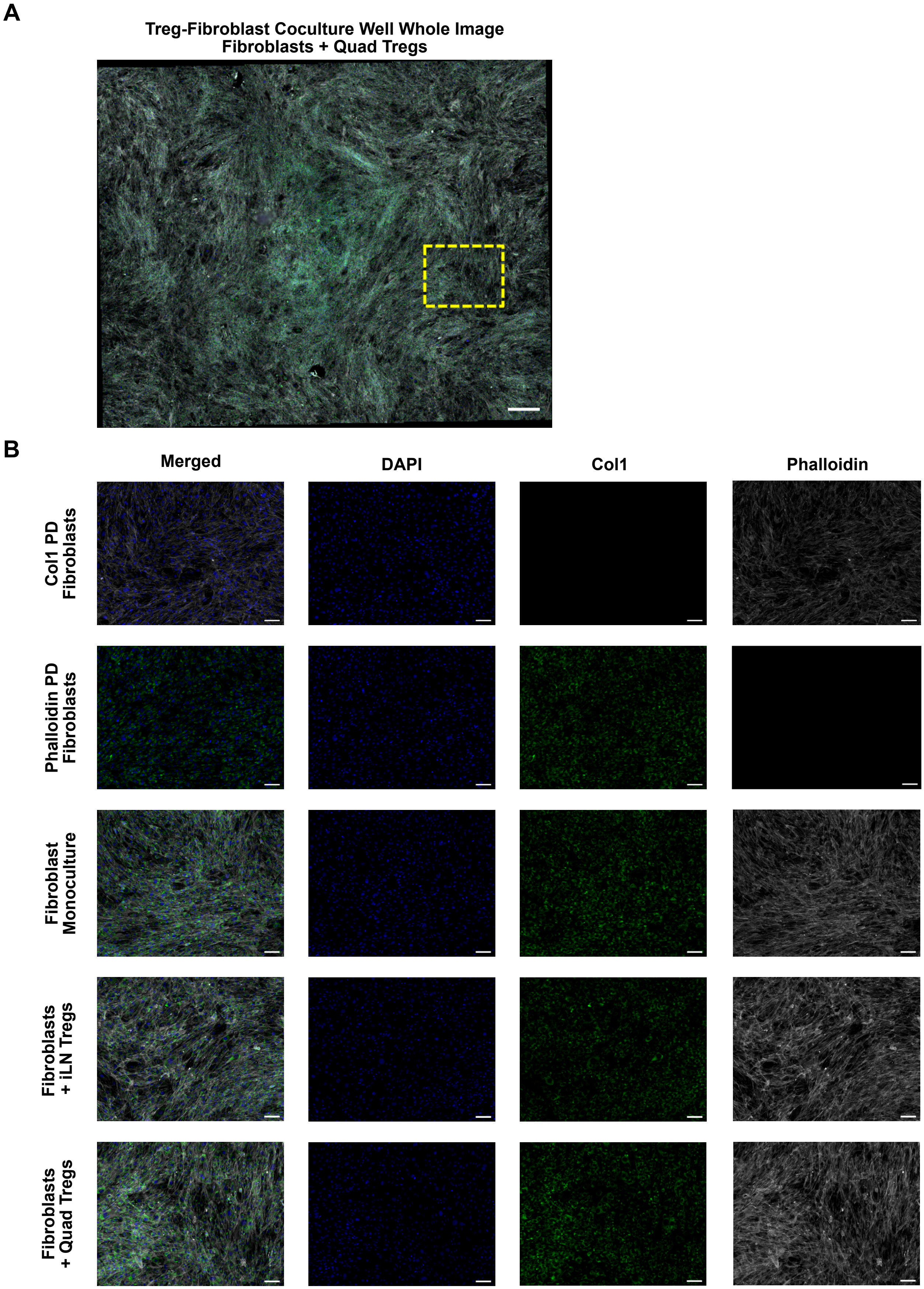


**Supplemental Figure 21. Imaging for phalloidin and collagen 1 after Treg-Fibroblast Coculture.**
A) Representative whole-well image of DAPI, Collagen1, and Phalloidin immunofluorescence staining in tissue culture from fibroblasts alone, fibroblasts + inguinal lymph node (iLN) Tregs, and fibroblasts + quadricep (quad) Tregs. Inset indicated with dashed yellow line. Scale bar = 500μm B) Merged and single channel images across matched primary delete (PD) controls and representative tissue insets. Scale bars = 100μm.

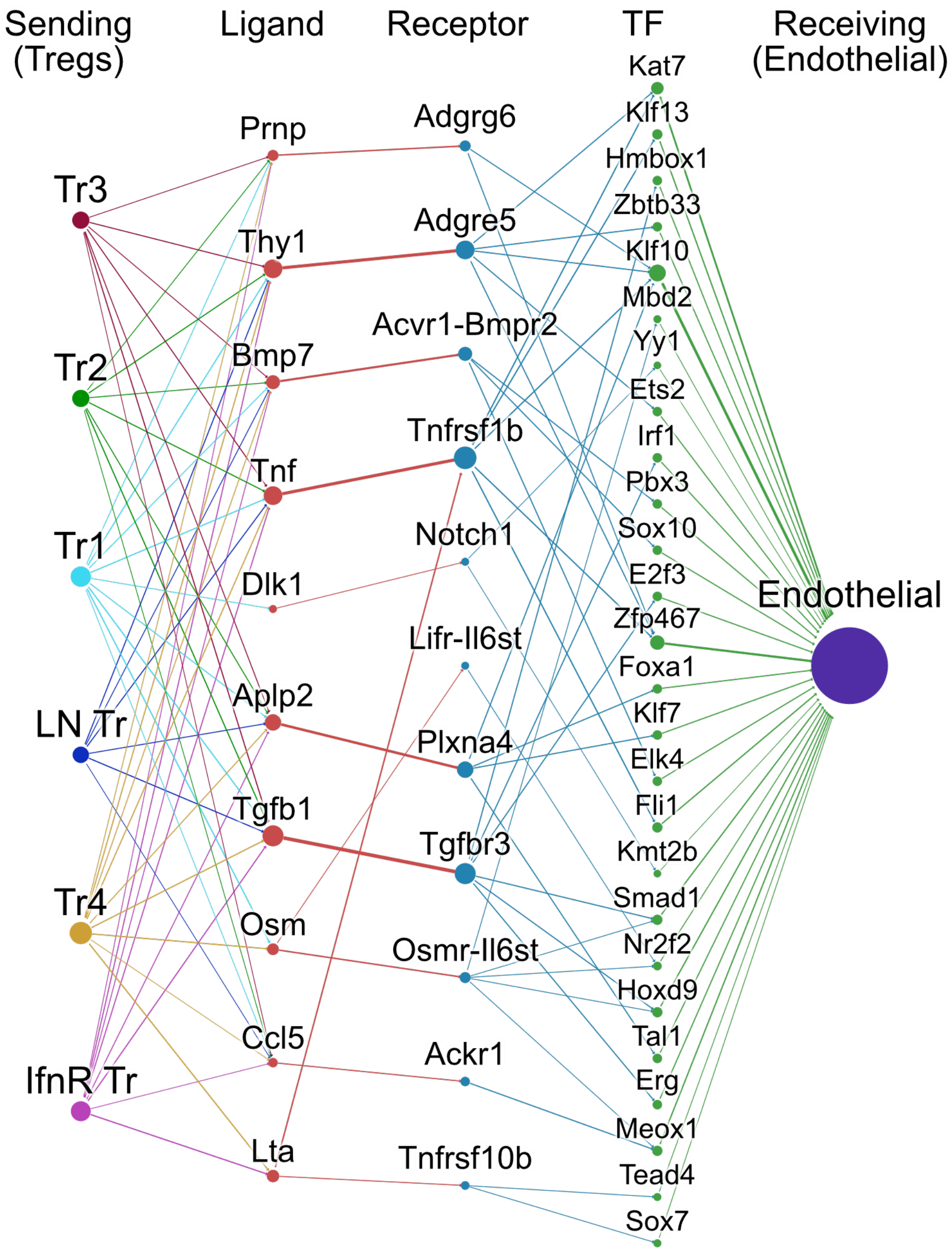


**Supplemental Figure 22. Treg to endothelial cell communication network.**
Network diagram depicting all predicted signaling pathways linking Treg subpopulations to endothelial cell population as identified in communication inference analysis. Node size and edge thickness correspond to number of pathways.

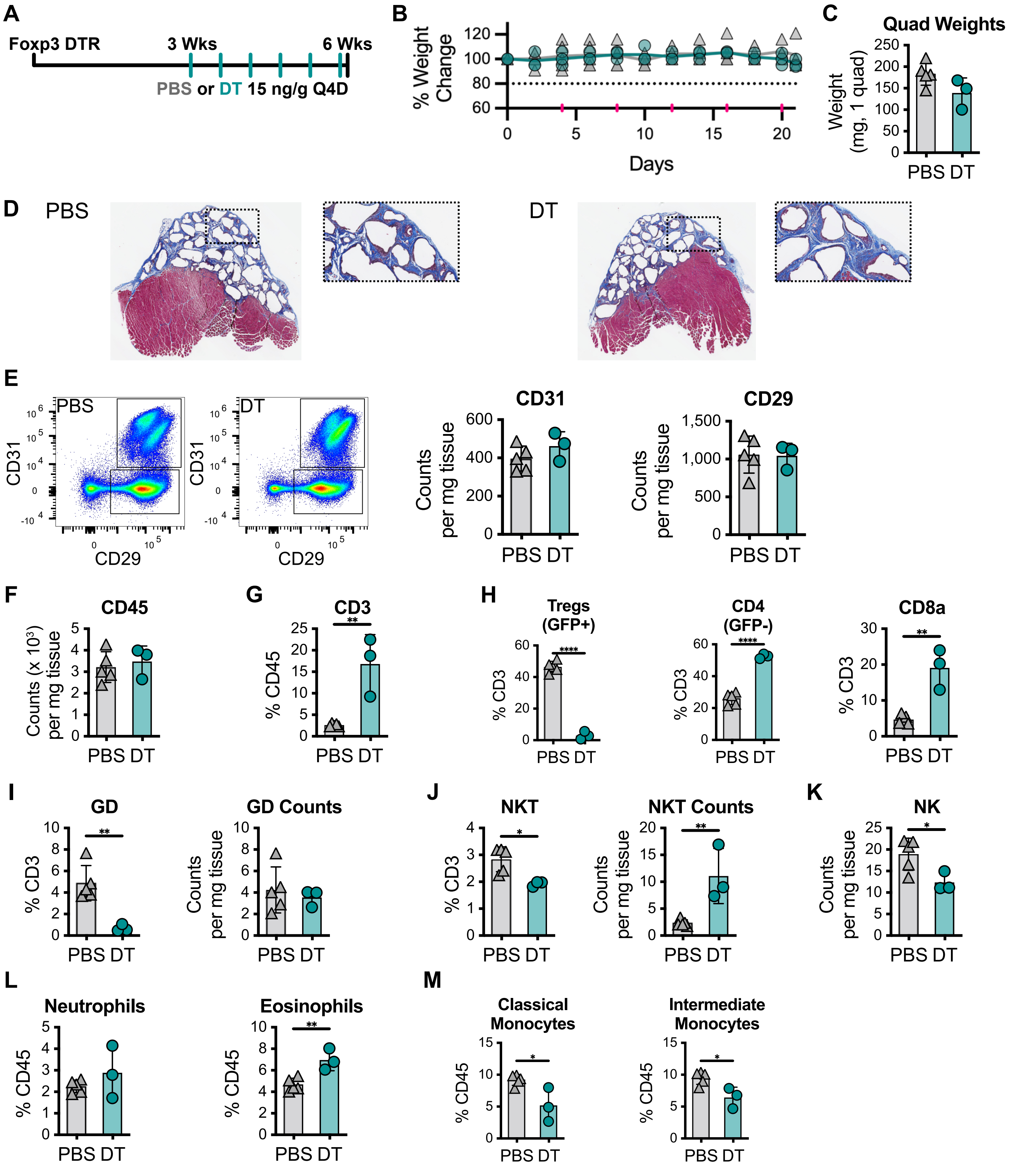


**Supplemental Figure 23. Q4D *in vivo* Treg depletion timeline, weights, histology, and immune quantification**
A) Experimental timeline for Q4D depletion protocol. B) Mouse weight changes over time for PBS vs DT treated groups. C) Quadriceps weight at harvest. D) Representative Masson's trichrome staining of whole quadreceps and fibrotic region inset for PBS (left) and DT (right) treated groups. E) Gating and quantification for CD29+ and CD29+CD31+ stromal populations by GFP Pan Stromal/Immune panel. F-M) Quantification of immune populations: F) CD45 counts, G) CD3 percentage of CD45+, H) Treg, CD4+ Tconv, CD8+ T cell percentages of CD3+, I) γδ T cell percentage of CD3+ and counts, J) NKT percentage of CD3+ and counts, K) Natural killer cell counts, L) Neutrophil and eosinophil percentages of CD45+, M) Classical and intermediate monocyte percentages of CD45+. Statistical significance assessed with unpaired two-sided t-test: $p < 0.05$ (*), $p < 0.01$ (**), $p < 0.001$ (***), $p < 0.0001$ (****).

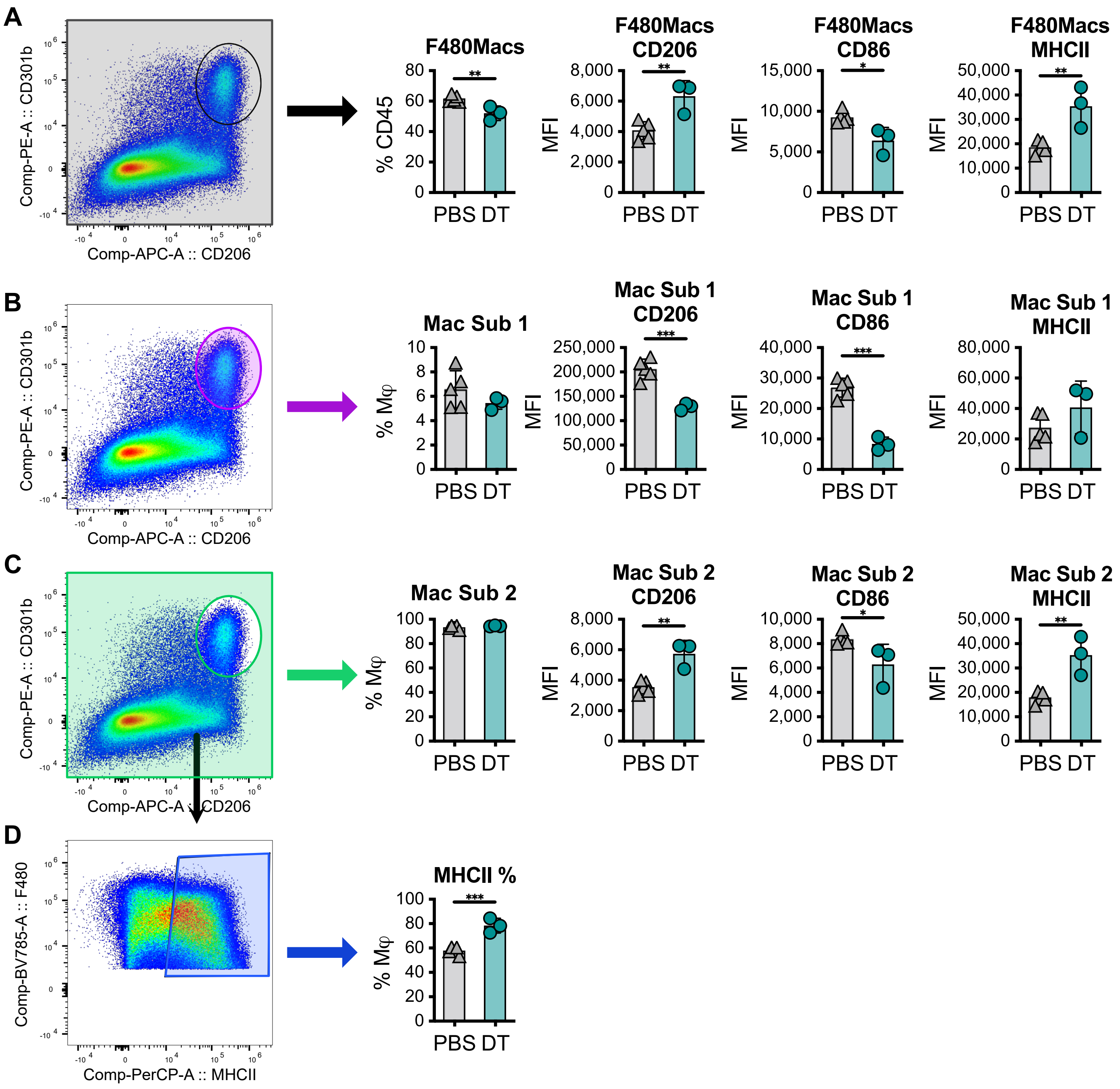


**Supplemental Figure 24. Macrophage subset quantification following Q4D Treg depletion**

A) Analysis of the macrophage population as a whole for CD206, CD86, and MHCII Median Fluorescence Intensity (MFI). B) Gating for macrophage subset 1 (CD206+CD301b+), percentage of macrophages, and MFI for CD206, CD86, and MHCII. C) Gating for macrophage subset 2 (complement to macrophage subset 1), percentage of macrophages, and MFI for CD206, CD86, and MHCII. D) Percentage of MHCII+ cells within macrophage subset 2. Transparent colored overlays are added over exported gating schemes to indicate gated subset.

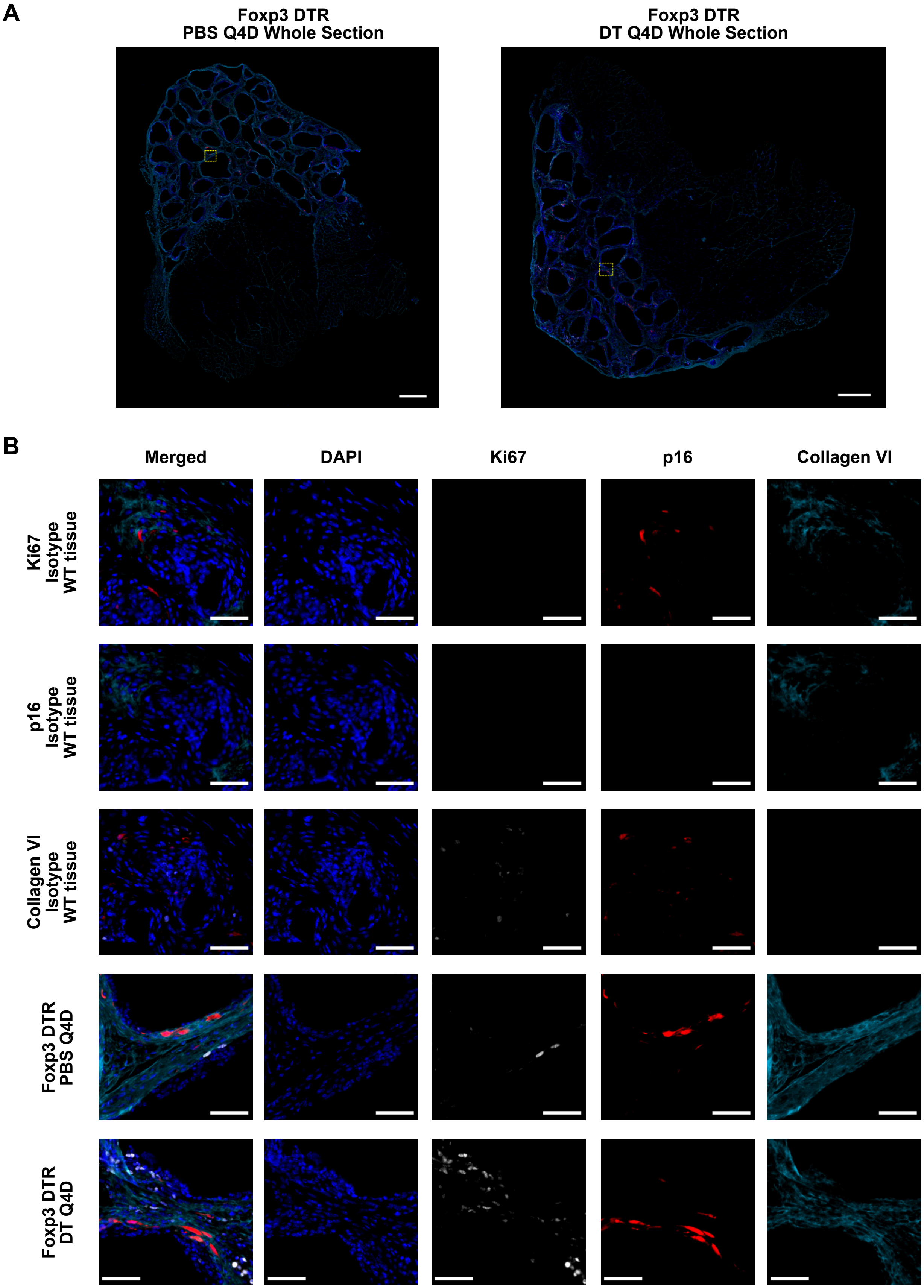


**Supplemental Figure 25. Imaging for proliferation and senescence markers following Q4D Treg depletion**
A) Whole-section images of DAPI/Ki67/p16 immunofluorescence staining in quadriceps from PBS and DT treated groups. Insets indicated with dashed yellow line and arrow. Scale bars = 500µm. B) Merged and single channel images across matched isotype controls and representative tissue insets. Scale bars = 50µm. Collagen VI staining not quantified in this analysis.

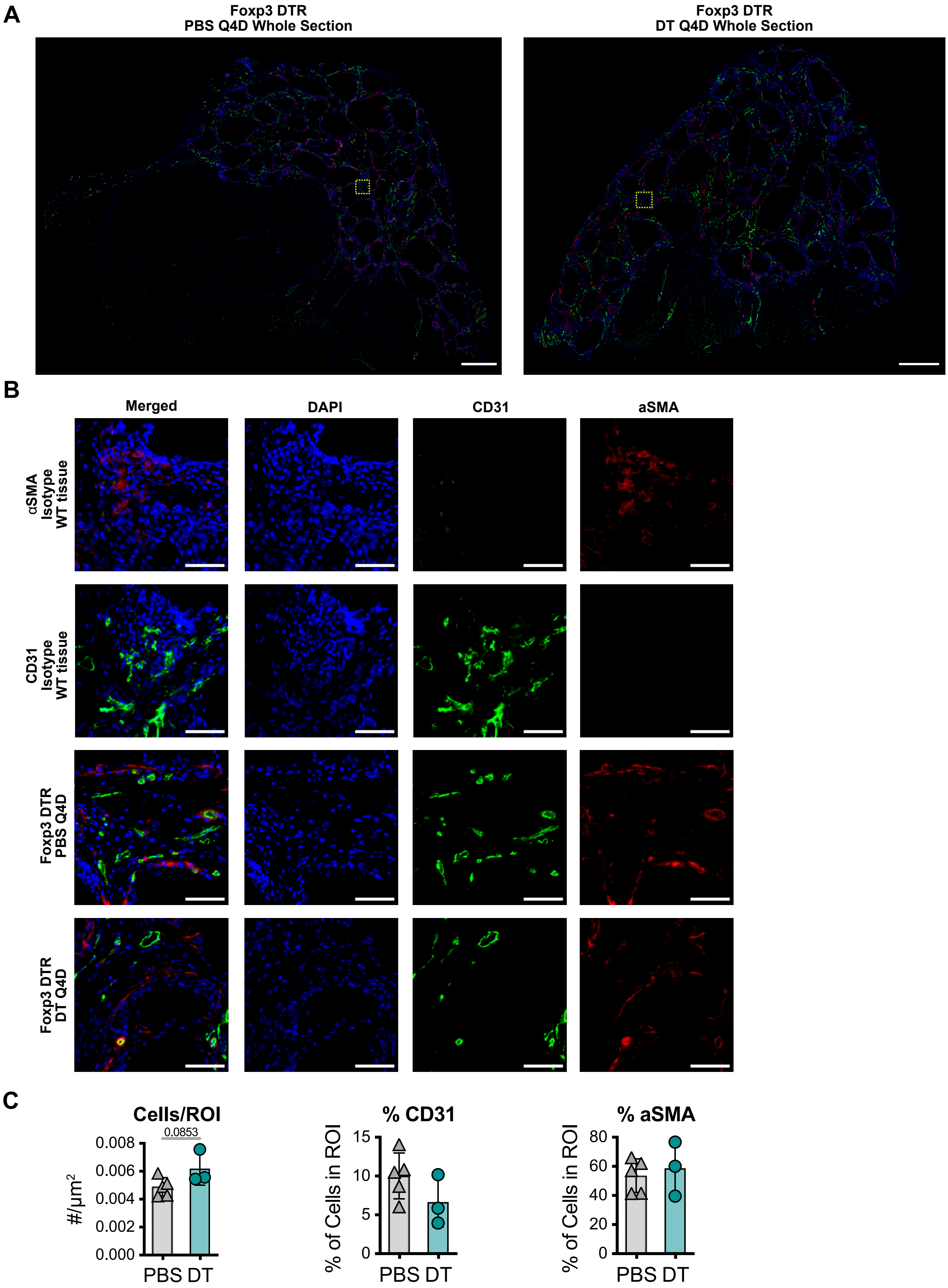


**Supplemental Figure 26. Imaging of vascular and stromal markers after Q4D Treg depletion**
A) Whole-section immunofluorescence images from PBS and DT treated quadriceps stained for DAPI, CD31, and αSMA. Inset indicated with yellow dashed line. Scale bars = 500μm. B) Merged and single channel images across matched isotype controls and representative tissue insets. C) Quantification of cells/ROI and percentage of CD31+ and αSMA+ cells within ROI. Scale bars = 50μm. Bar plots mean ± standard deviation.

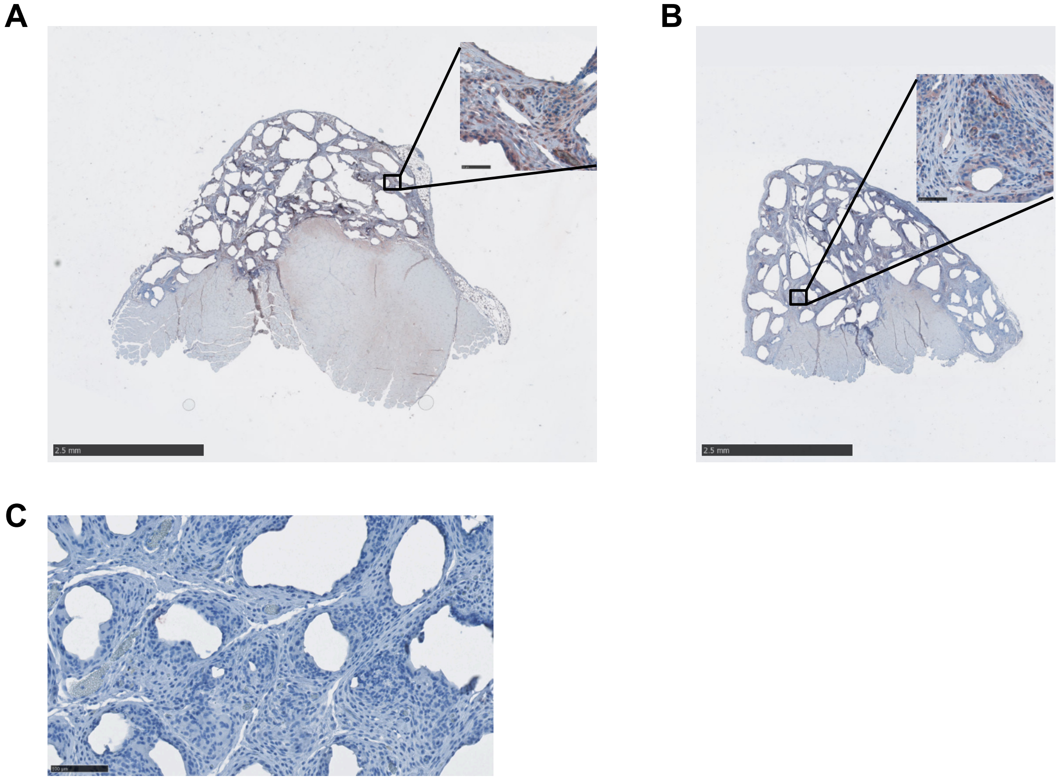


**Supplemental Figure 27. Immunohistochemistry staining of neovascular marker CD105 after Q4D Treg depletion**
A) Whole section immunohistochemistry images for CD105 staining of Q4D depletion study for PBS treated 6 week VML PCL Foxp3 DTR and (B) DT treated 6-week VML PCL Foxp3 DTR; scale bars = 2.5mm, inset scale bars = 50μm. (C) Representative image of CD105 staining control sample; scale bar = 100μm.

**Supplemental Table 1: T Cell Checkpoint Panel**

| Antigen | Clone | Fluor | Staining | Dilution | Manufacturer | Cat # | RRID |
|---|---|---|---|---|---|---|---|
| NK1.1 | PK136 | BV421 | Surface | 50 | BioLegend | 108732 | AB_10895916 |
| GITR | DTA-1 | BV480 | Surface | 200 | BD Biosciences | 746745 | AB_2744008 |
| CD45 | 30-F11 | BV510 | Surface | 200 | BioLegend | 103138 | AB_2563061 |
| CD4 | GK1.5 | BV605 | Surface | 300 | BioLegend | 100451 | AB_2564591 |
| GD TCR | GL3 | BV711 | Surface | 200 | BD Biosciences | 563994 | AB_2738531 |
| B220 | RA3-6B2 | BV750 | Surface | 200 | BioLegend | 103261 | AB_2734157 |
| PD1 | RMP1-30 | BV786 | Surface | 200 | BD Biosciences | 748264 | AB_2872692 |
| TCF | C63D9 | Alexa Fluor 488 | Intracellular | 100 | Cell Signaling | 6444S | AB_2797627 |
| CD3 | 17A2 | SparkBlue 550 | Surface | 100 | BioLegend | 100260 | AB_2832258 |
| CD8a | 53-6.7 | BB700 | Surface | 200 | BD Biosciences | 566409 | AB_2744467 |
| TIGIT | GIGD7 | PerCP-eFluor 710 | Surface | 100 | ThermoFisher | 46-9501-82 | AB_11150967 |
| RORgt | | PE | Intracellular | | | | |
| CTLA-4 | UC10-4B9 | PE-Dazzle 594 | Intracellular | 200 | BioLegend | 106318 | AB_2564496 |
| TIM3 | RMT3-23 | PE-Cy7 | Surface | 150 | BioLegend | 119716 | AB_2571932 |
| TOX | REA473 | APC | Intracellular | 150 | Miltenyi | 130-118-335 | AB_2751485 |
| Foxp3 | FJK-16s | eFluor 660 | Intracellular | 100 | ThermoFisher | 35-5773-82 | AB_11218094 |
| Viability | | Zombie NIR | Surface | 5000 | BioLegend | 423106 | |
| LAG3 | A7R34 | APC-eFluor 780 | Surface | 150 | ThermoFisher | 47-1271-82 | AB_1724012 |
| **Fixative:** True-Nuclear Transcription Factor Buffer Set | | | | | BioLegend | 424401 | |

**Supplemental Table 2: GFP T Cell Checkpoint Panel**

| Antigen | Clone | Fluor | Staining | Dilution | Manufacturer | Cat # | RRID |
|---|---|---|---|---|---|---|---|
| NK1.1 | PK136 | BV421 | Surface | 50 | BioLegend | 108732 | AB_10895916 |
| GITR | DTA-1 | BV480 | Surface | 200 | BD Biosciences | 746745 | AB_2744008 |
| CD45 | 30-F11 | BV510 | Surface | 200 | BioLegend | 103138 | AB_2563061 |
| CD4 | GK1.5 | BV605 | Surface | 300 | BioLegend | 100451 | AB_2564591 |
| GD TCR | GL3 | BV711 | Surface | 200 | BD Biosciences | 563994 | AB_2738531 |
| B220 | RA3-6B2 | BV750 | Surface | 200 | BioLegend | 103261 | AB_2734157 |
| PD1 | RMP1-30 | BV786 | Surface | 200 | BD Biosciences | 748264 | AB_2872692 |
| GFP | | GFP | Surface | | | | |
| CD3 | 17A2 | SparkBlue 550 | Surface | 100 | BioLegend | 100260 | AB_2832258 |
| CD8a | 53-6.7 | BB700 | Surface | 200 | BD Biosciences | 566409 | AB_2744467 |
| TIGIT | GIGD7 | PerCP-eFluor 710 | Surface | 100 | ThermoFisher | 46-9501-82 | AB_11150967 |
| CD25 | PC-61 | PE | Surface | 250 | BioLegend | 102008 | AB_312857 |
| LAG3 | C9B7W | PE-Dazzle 594 | Surface | 200 | BioLegend | 125224 | AB_2572081 |
| TIM3 | RMT3-23 | PE-Cy7 | Surface | 150 | BioLegend | 119716 | AB_2571932 |
| CD11b | M1/70 | AF700 | Surface | 400 | BioLegend | 101222 | AB_493705 |
| Viability | | Zombie NIR | Surface | 5000 | BioLegend | 423106 | |
| CD127 | A7R34 | APC eFluor 780 | Surface | 150 | eBioscience | 47-1271-82 | AB_1724012 |
| **Fixative:** FluoroFix Buffer | | | | | BioLegend | 422101 | |

**Supplemental Table 3: GFP Pan Immune Panel**

| Antigen | Clone | Fluor | Staining | Dilution | Manufacturer | Cat # | RRID |
|---|---|---|---|---|---|---|---|
| CD86 | GL-1 | BV421 | Surface | 200 | BioLegend | 105032 | AB_10898329 |
| CD19 | eBio1D3 | Super Bright 436 | Surface | 100 | eBioscience | 62-0193-82 | AB_2688107 |
| Ly6G | 1A8 | Pacific Blue | Surface | 200 | BioLegend | 127612 | AB_1877212 |
| Siglec-F | E50-2440 | BV480 | Surface | 150 | BD Biosciences | 746668 | AB_2743940 |
| CD45 | 30-F11 | BV605 | Surface | 300 | BioLegend | 103140 | AB_2562341 |
| Ly6C | HK1.4 | BV650 | Surface | 1200 | BioLegend | 128049 | AB_2800630 |
| GD TCR | GL3 | BV711 | Surface | 200 | BD Biosciences | 563994 | AB_2738531 |
| B220 | RA3-6B2 | BV750 | Surface | 200 | BioLegend | 103261 | AB_2734157 |
| F480 | BM8 | BV785 | Surface | 300 | BioLegend | 123141 | AB_2653667 |
| GFP | | | | | | | |
| CD3 | 17A2 | SparkBlue 550 | Surface | 100 | BioLegend | 100260 | AB_2832258 |
| MHCII | M5/114.15.2 | PerCP | Surface | 200 | BioLegend | 107624 | AB_2191073 |
| CD8a | 53-6.7 | BB700 | Surface | 200 | BD Biosciences | 566410 | AB_2744467 |
| CD301b | URA-1 | PE | Surface | 250 | BioLegend | 146804 | AB_2562944 |
| CD11c | N418 | PE-Dazzle 594 | Surface | 500 | BioLegend | 117348 | AB_2563654 |
| CD200R3 | Ba13 | PECy7 | Surface | 400 | BioLegend | 142212 | AB_2814045 |
| CD206 | C068C2 | APC | Surface | 200 | BioLegend | 141708 | AB_10896057 |
| NK1.1 | PK136 | AF647 | Surface | 200 | BioLegend | 108720 | AB_2132713 |
| CD11b | M1/70 | AF700 | Surface | 400 | BioLegend | 101222 | AB_493705 |
| Viability | | Zombie NIR | Surface | 3000 | BioLegend | 423106 | |
| CD9 | MZ3 | APCeFluor780 | Surface | 500 | BioLegend | 124814 | AB_2783073 |
| CD4 | GK1.5 | APC Fire 810 | Surface | 100 | BioLegend | 100480 | AB_2860583 |
| Fixative: FluoroFix Buffer | | | | | BioLegend | 422101 | |

**Supplemental Table 4: T Cell Sorting Panel**

| Antigen | Clone | Fluor | Staining | Dilution | Manufacturer | Cat # | RRID |
|---|---|---|---|---|---|---|---|
| CD45 | 30-F11 | BV605 | Surface | 150 | BioLegend | 103140 | AB_256341 |
| CD3 | 17A2 | Alexa Fluor 488 | Surface | 200 | BioLegend | 100210 | AB_389301 |
| Viability | | Zombie NIR | Surface | 1500 | BioLegend | 423106 | |

**Supplemental Table 5: GFP Pan Stromal Immune Panel**

| Antigen | Clone | Fluor | Staining | Dilution | Manufacturer | Cat # | RRID |
|---|---|---|---|---|---|---|---|
| CD86 | GL-1 | BV421 | Surface | 200 | BioLegend | 105032 | AB_10898329 |
| CD19 | eBio1D3 | Super Bright 436 | Surface | 100 | eBioscience | 62-0193-82 | AB_2688107 |
| Ly6G | 1A8 | Pacific Blue | Surface | 200 | BioLegend | 127612 | AB_1877212 |
| Siglec-F | E50-2440 | BV480 | Surface | 150 | BD Biosciences | 746668 | AB_2743940 |
| CD45 | 30-F11 | BV510 | Surface | 150 | BioLegend | | AB_ |
| GD TCR | GL3 | BV605 | Surface | 100 | | | AB_ |
| Ly6C | HK1.4 | BV650 | Surface | 1200 | BioLegend | 128049 | AB_2800630 |
| CD31 | | BV711 | Surface | 1200 | | | |
| B220 | RA3-6B2 | BV750 | Surface | 200 | BioLegend | 103261 | AB_2734157 |
| F480 | BM8 | BV785 | Surface | 300 | BioLegend | 123141 | AB_2653667 |
| GFP | | | | | | | |
| CD3 | 17A2 | SparkBlue 550 | Surface | 100 | BioLegend | 100260 | AB_2832258 |
| MHCII | M5/114.15.2 | PerCP | Surface | 200 | BioLegend | 107624 | AB_2191073 |
| CD8a | 53-6.7 | BB700 | Surface | 200 | BD Biosciences | 566410 | AB_2744467 |
| CD301b | URA-1 | PE | Surface | 250 | BioLegend | 146804 | AB_2562944 |
| CD11c | N418 | PE-Dazzle 594 | Surface | 500 | BioLegend | 117348 | AB_2563654 |
| CD200R3 | Ba13 | PECy7 | Surface | 400 | BioLegend | 142212 | AB_2814045 |
| CD206 | C068C2 | APC | Surface | 200 | BioLegend | 141708 | AB_10896057 |
| NK1.1 | PK136 | AF647 | Surface | 200 | BioLegend | 108720 | AB_2132713 |
| CD11b | M1/70 | AF700 | Surface | 400 | BioLegend | 101222 | AB_493705 |
| Viability | | Zombie NIR | Surface | 3000 | BioLegend | 423106 | |
| CD29 | MZ3 | | Surface | 300 | BioLegend | | AB_ |
| CD4 | GK1.5 | APC Fire 810 | Surface | 100 | BioLegend | 100480 | AB_2860583 |
| Fixative: FluoroFix Buffer | | | | | BioLegend | 422101 | |

**Supplemental Table 6: Immunofluorescence Staining Primary and Secondary**

| Antigen | Clone | Host species | Stock Concentration | Dilution | Manufacturer | Cat # | RRID |
|---|---|---|---|---|---|---|---|
| CD4 | EPR19514 | Rabbit | | 1000 | Abcam | ab183685 | AB_2686917 |
| Foxp3 | EPR22102-37 | Rabbit | | 100 | Abcam | ab215206 | AB_2860568 |
| CD31 | EPR17259 | Rabbit | 0.519 mg/mL | 2000 | Abcam | ab182981 | AB_2920881 |
| PDGFRβ | 28E1 | Rabbit | 4 μg/mL | 100 | Cell Signaling | 3169S | AB_2162497 |
| αSMA | EPR5368 | Rabbit | 0.147 mg/mL | 1000 | Abcam | ab124964 | AB_11129103 |
| DAPI | – | – | – | 10 | Akoya | FP1490 | |
| Opal 520 | – | – | – | 100 | Akoya | FP1487001KT | |
| Opal 570 | – | – | – | 150 | Akoya | FP1488001KT | |
| Opal 650 | – | – | – | 150 | Akoya | FP1496001KT | |

**Supplemental Table 7: TotalSeq-C for Hashing**

| Hashing Oligo | Antigen | Clone | Dilution | Manufacturer | Catalog Number |
|---|---|---|---|---|---|
| TotalSeq-C0301 | MHC Class I; CD45 | M1/42; 30-F11 | 100 | BioLegend | 155861 |
| TotalSeq-C0302 | MHC Class I; CD45 | M1/42; 30-F11 | 100 | BioLegend | 155863 |
| TotalSeq-C0303 | MHC Class I; CD45 | M1/42; 30-F11 | 100 | BioLegend | 155865 |
| TotalSeq-C0304 | MHC Class I; CD45 | M1/42; 30-F11 | 100 | BioLegend | 155867 |
| TotalSeq-C0305 | MHC Class I; CD45 | M1/42; 30-F11 | 100 | BioLegend | 155869 |

**Supplemental Table 8: Custom Primers for γδ VDJ sequencing**

| Name | Sequence | Final Concentration (μM) |
|---|---|---|
| 10X-FP1 | aatgatacggcgaccaccgagatctacactctttccctacacgacgctc | 0.5 |
| 10X-FP2 | aatgatacggcgaccaccgagatct | 1 |
| GDM-RP-1-1 | tcgaatctccatactgaccaagcttgac | 1 |
| GDM-RP-1-2 | gtcttcagcgtatccccttcctgg | 1 |
| GDM-RP-1-3 | ctttcaggcacagtaagccagc | 0.5 |
| GDM-RP-1-4 | tcttcagtcaccgtcagccaactaa | 0.5 |
| GDM-RP-2-1 | ccacaatcttcttggatgatctgagact | 0.5 |
| GDM-RP-2-2 | gtcccagtcttatggagatttgtttcagc | 0.5 |

**Supplemental Table 9: HTO Sequences for Demultiplexing Samples**

| ID | Read | Pattern | Sequence |
|---|---|---|---|
| HTO2 | R2 | 5PNNNNNNNNNN(BC) | GGTCGAGAGCATTCA |
| HTO3 | R2 | 5PNNNNNNNNNN(BC) | CTTGCCGCATGTCAT |
| HTO4 | R2 | 5PNNNNNNNNNN(BC) | AAAGCATTCTTCACG |
| HTO5 | R2 | 5PNNNNNNNNNN(BC) | CTTTGTCTTTGTGAG |
| HTO6 | R2 | 5PNNNNNNNNNN(BC) | TATGCTGCCACGGTA |

**Supplemental Table 10: Treg Canonical Markers Panel**

| Antigen | Clone | Fluor | Staining | Dilution | Manufacturer | Cat # | RRID |
|---|---|---|---|---|---|---|---|
| GITR | DTA-1 | BV480 | Surface | 200 | BD Biosciences | 746745 | AB_2744008 |
| CD45 | 30-F11 | BV510 | Surface | 200 | BioLegend | 103138 | AB_2563061 |
| CD4 | GK1.5 | BV605 | Surface | 300 | BioLegend | 100451 | AB_2564591 |
| CD11b | M1/70 | BV650 | Surface | 400 | BioLegend | 101259 | AB_2566568 |
| GD TCR | GL3 | BV711 | Surface | 200 | BD Biosciences | 563994 | AB_2738531 |
| CD3 | 17A2 | SparkBlue 550 | Surface | 100 | BioLegend | 100260 | AB_2832258 |
| CD8a | 53-6.7 | BB700 | Surface | 200 | BD Biosciences | 566409 | AB_2744467 |
| CD25 | PC-61 | PE | Surface | 250 | BioLegend | 102008 | AB_312857 |
| CTLA-4 | UC10-4B9 | PE-Dazzle 594 | Intracellular | 150 | BioLegend | 106318 | AB_2564496 |
| Foxp3 | FJK-16s | eFluor 660 | Intracellular | 100 | ThermoFisher | 35-5773-82 | AB_11218094 |
| Viability | | Zombie NIR | Surface | 5000 | BioLegend | 423106 | |
| CD127 | A7R34 | APC-eFluor 780 | Surface | 150 | ThermoFisher | 47-1271-82 | AB_1724012 |
| **Fixative**: True-Nuclear Transcription Factor Buffer Set | | | | | BioLegend | 424401 | |
